# *Peribacillus frigoritolerans* T7-IITJ, a potential biofertilizer, induces plant growth-promoting genes of *Arabidopsis thaliana*

**DOI:** 10.1101/2023.12.09.570910

**Authors:** Debankona Marik, Pinki Sharma, Nar Singh Chauhan, Neelam Jangir, Rajveer Singh Shekhawat, Devanshu Verma, Manasi Mukherjee, Moses Abiala, Chandan Roy, Pankaj Yadav, Ayan Sadhukhan

**Affiliations:** Department of Bioscience and Bioengineering, IIT Jodhpur, Jodhpur 342030, India; Department of Biochemistry, Maharshi Dayanand University, Rohtak 124001, India; Jodhpur City Knowledge and Innovation Foundation, IIT Jodhpur, Jodhpur 342030, India; Department of Biological Sciences, College of Basic and Applied Sciences, Mountain Top University, Prayer City, Nigeria; Department of Genetics and Plant Breeding, Agriculture University Jodhpur, Jodhpur 342304, India

**Keywords:** *Arabidopsis thaliana*, biofertilizer, drought, *Peribacillus frigoritolerans*, PGPR, transcriptome

## Abstract

This study aimed to isolate plant growth and drought tolerance-promoting bacteria from the nutrient-poor rhizosphere soil of several plant species from the Thar desert and unravel their molecular mechanisms of plant growth promotion, to develop effective biofertilizers for arid agriculture. Among our isolates of Thar desert rhizobacteria, *Enterobacter cloacae* C1P-IITJ, *Kalamiella piersonii* J4-IITJ, and *Peribacillus frigoritolerans* T7-IITJ, significantly enhanced root and shoot growth in the model plant *Arabidopsis thaliana* under PEG-induced drought stress in the lab. Whole genome sequencing and biochemical analyses of the non-pathogenic bacterium T7-IITJ revealed its plant growth-promoting traits, viz., solubilization of phosphate, iron, and nitrate and production of exopolysaccharides and auxin. Transcriptome analysis of *Arabidopsis thaliana* inoculated with T7-IITJ and exposure to drought revealed the induction of plant genes for photosynthesis, auxin and jasmonate signaling, nutrient mining and sequestration, redox homeostasis, and secondary metabolite biosynthesis pathways related to beneficial bacteria-plant interaction, but repression of many stress-responsive genes. Biochemical analyses indicated enhanced proline, chlorophyll, iron, phosphorous, and nitrogen content and reduced reactive oxygen species in plant tissues due to T7-IITJ inoculation. This bacterium could also improve the germination and seedling growth of *Tephrosia purpurea*, *Triticum aestivum,* and *Setaria italica* under drought. Additionally, T7-IITJ inhibited the growth of two plant pathogenic fungi, *Rhizoctonia solani,* and *Fusarium oxysporum*. These results suggest *P. frigoritolerans* T7-IITJ is a potent biofertilizer which can regulate plant genes promoting growth and drought tolerance.

## Introduction

Plant growth-promoting rhizobacteria (PGPR) ushered in a fresh green revolution in today’s world for their ability to increase crop productivity in the face of global warming, climate change, and increasing droughts and desertification (Lyu et al. 2020; Shah et al. 2021). PGPRs as “biofertilizers” provide a sustainable solution for global food security in the backdrop of the human population explosion. Biofertilizers are preferred over chemical fertilizers because of the high cost and environmental hazards associated with the latter (Dasgupta et al. 2021). PGPRs have diverse mechanisms of plant growth promotion, viz., increasing the soil phosphate and nitrate availability for plants, producing the hormone auxin, which results in faster root growth, and producing antibiotics to help the plant survive against pathogens (Pieterse *et al*. 2012; Olanrewaju, Glick and Babalola 2017). Phosphate-solubilizing PGPRs produce organic acids that chelate cations by their carboxyl or hydroxyl groups, releasing phosphate from the soil and making it available for plants (Billah *et al*. 2019). Another mechanism of action of rhizobacteria involves conferring induced systemic resistance to plants, achieved by the expression of *nonexpresser of pathogenesis-related genes 1* and through hormonal crosstalk (Borah *et al*. 2023). Apart from growth promotion and fighting phytopathogens, PGPRs also enhance drought stress tolerance by regulating stress-tolerance genes, producing protective osmolytes, siderophores, exopolysaccharides (EPS), and other volatile organic compounds (Ahmad *et al*. 2022; Abiala *et al*. 2023).

Deserts, hubs of stress-tolerant microbes, are potential hotspots for bioprospecting PGPRs. Two bacteria, *Pseudomonas* sp. M30-35 and *Bacillus* sp. WM13-24 from the roots of the Tengger desert plant, *Haloxylon ammodendron*, contributed to the increased drought tolerance of ryegrass by inhibiting reactive oxygen species (ROS), regulating plant hormone pathways, increasing relative water content, photosynthetic ability, and osmolyte content (He et al. 2021). *Cronobacter sakazakii*, *Proteus mirabilis*, and *Pseudomonas balearica* were isolated from the rhizosphere of *Aerva tomentosa* and *Panicum turgidum* in the Cholistan Desert in Pakistan. They showed promising growth-promoting traits in wheat, resulting in the upregulation of stress tolerance genes and increased cell stability (Zia *et al*. 2021). Another study in the Thar Desert, Rajasthan, India, highlighted the potential of *Serratia marcescens* to promote growth and confer systemic resistance in wheat (Singh and Jha 2016). However, exploring stress-tolerant PGPR strains from extreme desert environments is not exhaustive and holds immense promise for improving agricultural productivity in arid regions.

Motivated by the previous studies, we aimed at isolating stress-tolerant bacteria from the rhizosphere of Thar desert plants, which could be used for the development of biofertilizers for arid agriculture, as well as biocontrol agents for mitigating plant diseases in Rajasthan. We employed pure culture techniques, biochemical screening, and whole genome sequencing of the isolated bacterial strain to identify traits for plant growth promotion. Finally, phenotyping and transcriptome profiling were conducted in *Arabidopsis thaliana* to explore our isolated desert rhizobacteria’s molecular mechanisms of drought tolerance conferred to plants (Fig. 1).

**Fig. 1.**
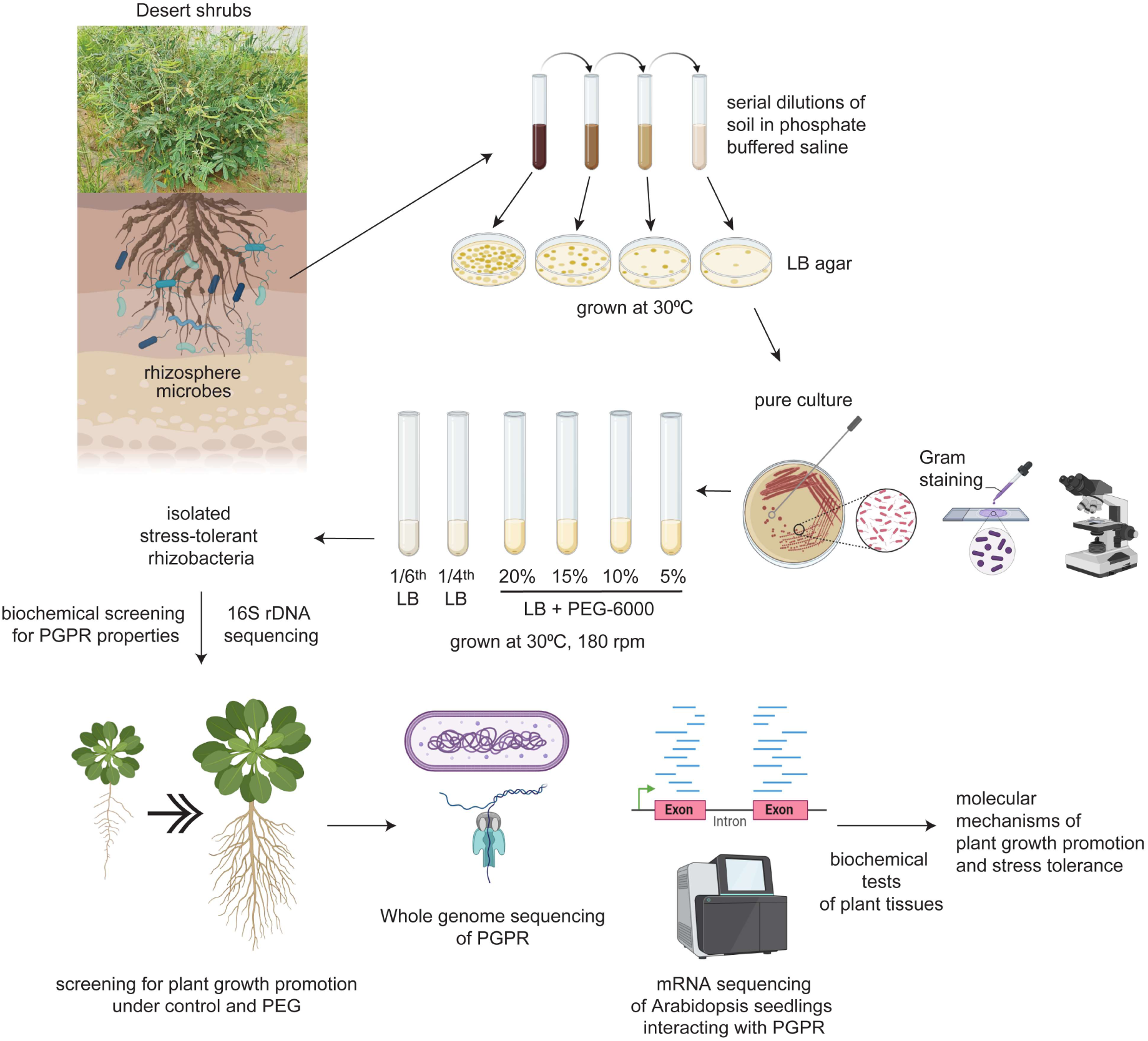
Schema of isolation and characterization of plant growth-promoting rhizobacteria from the Thar desert. Desert shrub rhizosphere soil samples at 12-15 cm depth were collected, diluted in phosphate-buffered saline, and plated on Luria-Bertani agar plates. Single colonies were streaked several times to obtain pure cultures, verified by Gram staining, microscopy, and 16S rDNA Sanger sequencing. The stress-tolerant bacteria were screened based on their ability to grow in LB media containing 5%, 10%, 15%, and 20% PEG-6000, which mimicked drought stress, and in diluted media (1/4^th^ and 1/6^th^ LB), which simulated nutrient deficiency stresses. Different biochemical tests were conducted to shortlist the stress-tolerant rhizobacteria based on plant-growth-promoting rhizobacteria (PGPR) properties, including phosphate solubilization, auxin and exopolysaccharide production, etc. Shortlisted bacteria were tested for their efficacy in promoting the seedling growth of *Arabidopsis thaliana* in hydroponic media containing PEG. The non-pathogenic safe strain was subjected to whole genome sequencing to understand its genes for plant growth promotion. Finally, transcriptome analysis by mRNA sequencing of *A. thaliana* seedlings inoculated with PGPR, and some biochemical tests were conducted to understand the molecular mechanisms of plant growth promotion and drought tolerance conferred by the PGPR. The figure was created using BioRender (https://www.biorender.com).

## Materials and Methods

### Isolation of stress-tolerant rhizobacteria

We collected soil samples from the rhizosphere of Thar desert plants at a depth of 10-15 cm at the IIT Jodhpur campus (26°28′34.38″N and 73°6′47.77″E), Rajasthan, India. The rhizosoil and control soil from the same site, where no plants were growing, were tested for pH and nutrient content at the Soil Testing Laboratory, Krishi Bhawan, Paota, Jodhpur. One gram of each rhizosoil sample was resuspended in 10 ml of 100 mM phosphate-buffered saline (PBS), pH 7.4, then serially diluted to 10^-1^, 10^-2^, and 10^-3^ in PBS. Fifty microliters of each dilution were evenly spread on a Luria Bertani (LB) agar plate and incubated overnight at 30^°^C. Bacterial colonies observed after the incubation were re-streaked to obtain pure cultures, verified by Gram staining and observation under a light microscope with 100x magnification. Pure isolates were grown on LB broth supplemented with 15% and 20% polyethylene glycol (PEG-6000) (Rajeswar and Narasimhan, 2021), as well as in 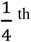 and 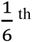 strength LB broth to screen drought and nutrient-deficiency-tolerant bacteria (Abiala et al. 2023). The bacterial growth was measured by reading the absorbance at 600 nm (A_600 nm_) with a UV-1800 double-beam spectrophotometer (Shimadzu Corporation, New Delhi, India) of the microbial culture grown at 30°C with continuous shaking at 180 rpm for 16 h. The extreme temperature tolerance of the isolates was determined by inoculating them in LB media and observing overnight growth at 5^°^C, 15^°^C, 30^°^C, 45^°^C, and 60^°^C with shaking at 180 rpm. The pH tolerance was determined by observing overnight growth at 30^°^C and 180 rpm in LB media with pH 3, 7, 9, and 11.

### Genomic DNA isolation from rhizobacteria and 16S rDNA sequencing

Bacteria were inoculated in 2 ml of LB broth and grown overnight at 30^°^C with 180 rpm shaking in an incubator (iGene Lab Serve, New Delhi, India). The culture was centrifuged for 15 min at 5,000 rpm in a Sorvall™ Legend™ centrifuge (Thermo Fisher Scientific, New Delhi, India), and the bacterial pellet was resuspended in CTAB buffer containing 2% SDS. It was incubated at 60^°^C for 0.5 h in a Thermomixer C (Eppendorf, Chennai, India). The samples were RNase-treated, then further extracted using phenol, chloroform, and isoamyl alcohol 24:24:1, and finally precipitated using isopropanol (Kumar, Kaur and Sandhu 2014). Quantitative and qualitative analysis of the isolated DNA was performed using a VWR mySPEC microvolume UV-visible spectrophotometer (VWR Lab Products Private Limited, Bangalore, India). The isolated genomic DNA was subjected to a PCR with forward and reverse 16S rDNA universal primers (16S_Fw: 5’-AGRGTTYGATYMTGGCTCAG-3’ and 16S_Rv: 5’-RGYTACCTTGTTACGACTT-3’) to get an amplicon of 1.5 kb, in a T100 Thermal Cycler (Bio-Rad, Gurugram, India) at 95^°^C for 5 min, followed by 34 cycles of 95^°^C for 30 s, 55^°^C for 45 s, and 72^°^C for 1.5 min, and a final extension at 72^°^C for 10 min. The 16S rDNA PCR-amplified fragments were purified by Wizard^®^ SV Gel and PCR Clean-Up System (Promega, New Delhi, India) and digested by HaeIII (SRL, New Delhi, India) at 37^°^C for 1 h. Duplicate samples were eliminated by observing similar banding patterns of the digested 16S rDNA fragments in a 2% agarose gel. De-duplicated 16S rDNA amplicons were sequenced by Eurofins Genomics India Private Ltd. (Bangalore, India) using Sanger sequencing chemistry. The sequences were analyzed by FinchTV chromatogram viewer, and non-chimeric sequences were uploaded to the EzBiocloud database (https://www.ezbiocloud.net/) to identify the pure bacterial isolates. The database hit strain with 100% sequence similarity, and completeness with the submitted sequence was considered the identification result.

### Biochemical characterization of rhizobacteria

The catalase, oxidase, starch hydrolysis, and citrate utilization tests for bacteria were conducted as described earlier (Alves *et al*. 2002; MacWilliams 2009; Reiner 2010; Shields and Cathcart 2013). A standardized procedure was followed for the Methyl Red-Voges Proskauer (MR-VP) test to assess the type of fermentation pathways (Mcdevitt 2009).

### Identification of plant growth-promoting traits of rhizobacteria

For testing phosphate solubilizing abilities, overnight liquid cultures of the bacterial isolates were centrifuged, and the pellets were resuspended in different volumes of PBS to attain a final A_600 nm_ = 1, corresponding to 8 × 10^8^ cells/ml. One microliter of the inoculum, thus prepared, was spotted on phosphate solubilization media and incubated for seven days at 30^°^C (Abiala, Sadhukhan and Sahoo 2023). After seven days, the diameters of the colony (CD) and the halo zone around the colony (HZD) were noted to calculate the solubilization index (SI) by the formula SI = (HZD + CD)/ CD. Indole acetic acid (IAA) production in liquid bacterial growth media was detected spectrophotometrically by a standard procedure (Chandra, Askari and Kumari 2018). The EPS production ability of the isolates was determined by the anthrone reagent using a glucose standard curve (Kvíderová *et al*. 2019). Bacteria were screened for nitrate reductase activity following the conversion of nitrate to nitrite, as described earlier (Kim and Seo 2018). The siderophore-producing abilities of bacteria were determined by the chrome azurol S (CAS) and hexadecyltrimethylammonium bromide (HDTMA) assay (Louden, Haarmann and Lynne 2011).

### Assessment of anti-fungal activity of rhizobacteria

The biocontrol potential of rhizobacteria was studied against two plant pathogenic fungi, *Rhizoctonia solani* and *Fusarium oxysporum,* by measuring the disk assay (Berkow et al. 2020). The fungal strains were grown in YEPD broth with continuous shaking at 180 rpm for 24 hours at 28°C in separate tubes. One hundred microliters of overnight grown fungal culture (A_600 nm_ = 1.0) were spread on solid LB agar plates. A sterile filter paper disc was placed on the center of the scale, and 50 µl of an overnight-grown bacterial culture (OD_600nm_ = 1.0) was added to the disc. Finally, the plate was incubated at 28°C for 48 hours and photographed to measure the diameter of the halo zone of fungal growth inhibition.

### Assessment of plant growth-promoting ability of rhizobacteria

The promotion of plant growth and drought tolerance by the bacterial isolates C1P-IITJ, J4-IITJ, and T7-IITJ was assessed on *A. thaliana* (ecotype Columbia; Col-0). Plant seeds were surface sterilized with 80% ethanol and 1% sodium hypochlorite by rinsing them 2 min in each, then washed five times with deionized water. For the germination assay, surface-sterilized seeds of *Tephrosia purpurea*, *Setaria italica*, and *Triticum aestivum* were placed on a filter paper in a sterile glass petri plate, soaked in 20 ml autoclaved deionized water without or with 5% PEG-6000. In half the plates, 1 ml overnight grown bacterial culture resuspended in PBS (8 × 10^8^ cells) was added, and sterile PBS in the other. The plates were incubated at 28°C in the dark, and germination rates were noted after 5 d.

For the plant growth assay in soil, sterilized and imbibed Col-0 seeds were sown in sterilized soil mix in 2.5-inch pots. The soil mix was prepared by mixing desert soil collected from the exact sampling location with soil-rite mix (KELTECH Energies Ltd., Bangalore, India) in a ratio of 3:1, autoclaving, and keeping aside for a few days. The pots were inoculated with 1 ml of the bacterial suspension 10 d after germination. Similarly, sterile PBS was added to control pots. Plants were grown at 22 ± 2°C under 55% relative humidity and 80 µmole m^−2^ s^−1^ of light intensity in a 12 h day/12h night cycle. After one month, the plants were removed from the soil, washed, and photographed.

For the seedling growth assay in hydroponics, sterilized Col-0 seeds were imbibed in deionized water for three days at 4°C, then sown uniformly on N50 nylon meshes, mounted on photographic frames (Sadhukhan *et al*. 2017) and floated on 300 ml of modified ¼^th^ Hoagland media, pH 5.8, without or with 5% PEG-6000 in sterile magenta boxes (Abiala *et al*. 2023). Twenty seeds were used for each sample. After a day, the media was inoculated separately with 1 ml of the bacterial isolates, and the seedlings were allowed to grow for 14 d. For the control samples, media were inoculated with sterile PBS without bacteria. The seedlings were transferred to solidified agar plates and photographed with a scale at the end of 14 d. The root length and shoot diameter were measured from the images as described previously (Sadhukhan *et al*. 2017). On water-soaked cotton, *T. purpurea* seeds were imbibed at 30°C in a sterile petri dish. The sprouted seedlings were transferred to hydroponic growth conditions in 300 ml ¼^th^ Hoagland media without or with 10% PEG-6000 in magenta boxes. Six seeds were used for each condition; in this case, the inoculation of bacteria was performed after two days of transfer to the hydroponic media. All experiments were repeated three times.

### Whole genome sequencing of rhizobacteria

The genomic DNA of T7-IITJ was purified by HiPura^®^ bacterial genomic DNA isolation kit (Himedia). Sequencing was performed on Nanopore MinION MK1C with midnight pipeline (Oxford Nanopore Technologies, Oxford, UK). The generated long reads were assembled using the Unicycler assembler, which uses SPAdes as a *de novo* assembler (Wick *et al*. 2017). Prokka 1.12. J-species software (http://jspecies.ribohost.com/jspeciesws/) was used to annotate the functions of the genes. Genomes of *Peribacillus* strains were selected based on average nucleotide identity (ANI), downloaded from the NCBI web server. The phylogenetic relationship with reference genomes of *Peribacillus* species was determined using the Roary platform version 3.11.2. Assembled contigs were used to draw a circular genomic map using the Proksee tool (https://proksee.ca/). The antipathogenic gene clusters were identified by the antiSMASH version 7.0.1 (Medema *et al*. 2011). The Comprehensive Antibiotic Resistance Database (CARD) identifier predicted the antibiotic resistance genes in the bacterial genome. The absence of virulence and antibiotic resistance genes within the mobile genetic elements was confirmed using the VRprofile2 web tool (Wang *et al*. 2022a).

### Measurement of chlorophyll, proline, lipid peroxidation, reactive oxygen species, and nutrient levels in plant tissues

Col-0 seeds were grown hydroponically in ¼^th^ Hoagland media for seven days to obtain uniform-sized seedlings, and after that, they were inoculated with *Peribacillus frigoritolerans* T7-IITJ. After seven days of plant-bacteria interaction, 5% PEG-6000 was applied to both inoculated and non-inoculated plants for five days, and tissues were harvested and weighed. The control seedlings were grown for a similar time in media without PEG. The chlorophyll and proline content of the Col-0 leaves were measured as per standard protocols (Arnon 1949; Bates, Waldren and Teare 1973). Malonaldehyde (MDA) levels in Col-0 seedlings, indicative of the membrane lipid peroxidation, were determined by Heath and Packer’s method(1968). The hydrogen peroxide (H_2_O_2_) and superoxide radical levels (O_2_^•–^) in seedlings were measured as described earlier (Elstner and Heupel, 1976; Sagisaka, 1976). Total nitrogen, phosphorous, and iron of the plant tissues were determined as described previously (Hartmann and Asch, 2018; Koistinen et al. 2020; Pradhan and Pokhrel, 2013).

### Transcriptome analysis of rhizobacteria-inoculated Arabidopsis

For transcriptome analysis, 200 Col-0 seeds were grown hydroponically and inoculated with T7-IITJ after seven days, as described above. Seedlings were snap-frozen in liquid nitrogen after seven days of plant-bacteria interaction and further PEG treatment for five more days. RNA was isolated by the CTAB-LiCl method, and its quality was checked using the Agilent 4150 TapeStation system (Agilent Technologies, CA, USA) and Qubit 4 fluorometer (Thermo Fisher). RNA-Seq libraries were prepared using the NebNext Ultra II RNA library prep kit, following the manufacturer’s instructions (New England Biolabs, MA, USA). High-quality RNA-Seq libraries were sequenced on the NovaSeq 6000 V1.5 platform (Illumina, Gurgaon, India) in paired-end reads (2×150 bp). The raw transcriptome sequences were submitted to the NCBI SRA database with a BioProject accession number PRJNA1020649. The quality of the raw sequence reads GC content and library complexity were ascertained by the FastQC software (Andrews 2010). The adapter trimming and quality trimming at Q30 were performed using AdapterRemoval ver. 2.3.2. (Schubert, Lindgreen and Orlando 2016). The clean reads were aligned to the reference *A. thaliana* genome (GCF000001735.4) using hisat2 version 2.2.1 (Zhang *et al*. 2021). The expression estimation was performed using featureCounts v2.0.3 (Liao, Smyth and Shi 2014). The differential gene expression (DEG) analysis was done in R using edgeR, ggplot2, and DEseq2 packages (Love, Huber and Anders 2014; Chen, Lun and Smyth 2016; Wickham 2016). Furthermore, the genes were filtered based on false discovery rate (*FDR*) < 0.05 and log_2_(fold-change) ≥ 1 or ≤ -1 to obtain the differentially expressed genes. The gene set enrichment analysis was performed using KOBAS 3.0 using the Kyoto Encyclopaedia of Genes and Genomes (KEGG), BioCyc, and Gene Ontology (GO) databases (Bu *et al*. 2021). A gene co-expression network was constructed with strongly upregulated genes in the transcriptome analysis (log_2_FC > 2.5) with the GENEMANIA web application (Warde-Farley *et al*. 2010). GENEMANIA automatically assigned the weightage to find the maximum interconnectivity between the query genes based on known interactions. The public *A. thaliana* gene expression datasets GSE10576, GSE39384, GSE24348, GSE4684, GSE7641, GSE577, GSE16722, GSE14502, and GSE31158 were mined by the software to construct the co-expression network. We expanded the network analysis into a functional protein-protein interaction (PPI) network and enrichment analysis with all the differentially expressed genes with -1 < log_2_FC > 1 (Table S5) with Cytoscape 3.10.1 (Excoffier *et al*. 2017) using the Search Tool for the Retrieval of Interacting Genes (STRING) database (Szklarczyk *et al*. 2023). A threshold interaction score of 7.5 was deemed statistically significant for considering pairwise interactions. We explored hub genes in the network by ranking the nodes based on 11 topological features and identifying common nodes for all features with the cytoHubba application in the Cytoscape tool.

### Validation of the transcriptome results by quantitative real-time PCR

The expression of the six most significantly induced genes, based on *FDR* values, was validated by quantitative real-time PCR (qPCR). *A. thaliana* seedlings were co-cultivated with T7-IITJ in hydroponics and treated with PEG like the transcriptome experiment, and samples snap frozen with liquid nitrogen. Tissues were crushed using mortars and pestles to isolate RNA using the SV total RNA isolation system (Promega) and converted to cDNA using PrimeScript™ IV 1st strand cDNA Synthesis Mix (DSS Takara Bio India Pvt. Ltd., New Delhi, India). QPCR was carried out using Brilliant II SYBR master mix (Agilent Technologies India Pvt Ltd, Chandigarh, India) in an Mx3000P qPCR System (Agilent). The relative expression levels between T7-IITJ-inoculated and non-inoculated (control) seedling samples were estimated using the standard curve method. The *AtUBQ1* gene was used as an internal standard to normalize between cDNA samples. All gene-specific oligonucleotide primers used for the qPCR analysis were designed using the Primer3web tool version 4.1.0 (Koressaar *et al*. 2018) and are listed in Table S1.

## Results

### The rhizospheres of Thar desert plants contain stress-tolerant bacteria

Several native and invasive desert plants, viz., *Crotolaria burhia*, *Prosopis cineraria*, *Prosopis juliflora*, *Senna alexandrina*, *Solanum virginianum*, *and Tephrosia purpurea,* were found to grow in the IIT Jodhpur campus located on the outskirts of the Thar desert. We isolated 30 pure bacterial cultures from the rhizosphere of these plants in LB media (Table 1). The soil around these plants was alkaline (pH 8.5-8.7) and contained low nitrogen levels, organic carbon, iron, zinc, molybdenum, boron, and medium levels of potassium and phosphorous (Table S2). To test the stress tolerance of the rhizobacterial isolates obtained from the above soil, they were grown in drought and nutrient deficiency stress conditions in the laboratory. Among them, 12 isolates grew in PEG-containing LB media, 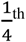, and 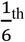 LB, while no growth was observed in the laboratory strain *E. coli* under similar conditions. This indicated the tolerance of these 12 isolates to drought and nutrient deficiency conditions (Table 1). Among the 12 isolates, J6-IITJ was highly resistant to drought stress, and C1P-IITJ, J4-IITJ, and T7-IITJ could tolerate moderate to high levels of drought stress under 15% and 20% PEG. Under 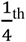 and 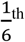 LB, T7-IITJ, J3-IITJ, and K4-IITJ showed maximum growth, whereas J4-IITJ and C1P-IITJ exhibited moderate tolerance to nutrient-deficient conditions. Our isolates C1P-IITJ, J4-IITJ, and T7-IITJ showed maximum growth under pH 9, and three exhibited tolerances to high (45°C) and low (15°C) temperatures.

**Table 1.** Growth of Thar desert rhizobacteria under extreme conditions.

| Isolate | Source plant | Morphology | Cell density ($10^8$ cells/ml) | | | | | | | | | | |
| --- | --- | --- | --- | --- | --- | --- | --- | --- | --- | --- | --- | --- | --- |
| | | | PEG | | Nutrient deficiency | | pH | | | Temperature ($^{\circ}$C) | | | |
|  |  |  | 15% | 20% | 1/4<sup>th</sup> LB | 1/6<sup>th</sup> LB | 3 | 9 | 11 | 5 | 15 | 45 | 60 |
| C1P-IITJ | *Crotolaria burhia* | Gram -ve bacillus | 8.53 ± 0.06 | 3.62 ± 0.01 | 6.58 ± 0.02 | 6 ± 0.12 | 1.1 ± 0.11 | 2.78 ± 0.03 | 1.66 ± 0.01 | 0.66 ± 0.04 | 1.73 ± 0.043 | 2.04 ± 0.02 | 0.7 ± 0.032 |
| S1P-IITJ | *Senna alexandrina* | Gram -ve coccus | 2.17 ± 0.02 | 1.72 ± 0.45 | 5.76 ± 0.02 | 4.32 ± 0.06 | 0.584 ± 0.12 | 2.86 ± 0.02 | 1.65 ± 0.04 | 0.44 ± 0.08 | 1.98 ± 0.087 | 2.24 ± 0.0654 | 0.63 ± 0.076 |
| T7-IITJ | *Tephrosia purpurea* | Gram +ve bacillus | 5.09 ± 0.1 | 4.46 ± 0.014 | 13.4 ± 0.14 | 8.63 ± 0.09 | 3.35 ± 0.098 | 9.62 ± 0.021 | 0.624 ± 0.23 | 1.5 ± 0.07 | 5.52 ± 0.43 | 4.25 ± 0.87 | 1.86 ± 0.87 |
| T3-IITJ | | Gram -ve bacillus | 6.06 ± 0.09 | 5.96 ± 0.56 | 1.68 ± 0.65 | 0.96 ± 0.16 | 1.99 ± 0.19 | 3.06 ± 0.198 | 3.48 ± 0.98 | 0.38 ± 0.67 | 0.8 ± 0.12 | 2.4 ± 0.87 | 3.29 ± 0.87 |
| S2-IITJ | *Solanum virginianum* | Gram -ve bacillus | 7.21 ± 0.01 | 4.49 ± 0.13 | 7.12 ± 0.45 | 6.24 ± 0.56 | 0 | 7.74 ± 0.18 | 6.58 ± 0.12 | 0.8 ± 0.9 | 1.3 ± 1.2 | 3.32 ± 0.8 | 2.45 ± 0.1 |
| S3-IITJ | | Gram -ve bacillus | 8.41 ± 0.03 | 4.87 ± 1.6 | 7.2 ± 0.6 | 7.04 ± 0.4 | 0 | 4.46 ± 0.23 | 4.71 ± 0.67 | 0.44 ± 0.45 | 2.2 ± 1.5 | 2.31 ± 0.7 | 1.54 ± 0.6 |
| S4-IITJ | | Gram -ve bacillus | 7.9 ± 0.02 | 4.18 ± 1.2 | 7.68 ± 1.7 | 6.64 ± 0.87 | 0.52 ± 0.43 | 6.16 ± 0.67 | 6.85 ± 0.43 | 0.81 ± 0.45 | 1.56 ± 0.32 | 3.24 ± 0.98 | 2.1 ± 0.456 |
| J1-IITJ | *Prosopis juliflora* | Gram +ve bacillus | 2.97 ± 0.1 | 0.6 ± 1.4 | 4.65 ± 1.2 | 1.89 ± 0.5 | 2.22 ± 1.2 | 7.1 ± 0.67 | 0.792 ± 0.34 | 3.46 ± 0.67 | 2.4 ± 0.78 | 4.03 ± 0.67 | 1.71 ± 0.98 |
| J3-IITJ | | Gram +ve coccus | 7.96 ± 0.12 | 3.72 ± 0.4 | 12 ± 1.2 | 8.21 ± 1.3 | 1.98 ± 0.99 | 2.55 ± 0.98 | 1.17 ± 0.65 | 1.91 ± 0.34 | 1.61 ± 0.69 | 4.42 ± 0.29 | 2.25 ± 0.19 |
| J4-IITJ | | Gram -ve bacillus | 7.37 ± 0.11 | 4.58 ± 0.11 | 7.27 ± 0.81 | 3.35 ± 0.93 | 0.264 ± 0.44 | 1.75 ± 0.998 | 0.288 ± 0.22 | 0.24 ± 0.12 | 2.16 ± 0.188 | 1.7 ± 0.18 | 0.39 ± 1.5 |
| J6-IITJ | | Gram +ve bacillus | 10.8 ± 0.02 | 10.4 ± 0.95 | 5.81 ± 0.78 | 1.24 ± 0.788 | 2.52 ± 0.88 | 9.14 ± 0.42 | 0.88 ± 0.56 | 0.696 ± 0.94 | 4.05 ± 0.32 | 6.35 ± 0.987 | 0.96 ± 0.555 |
| K4-IITJ | *Prosopis cineraria* | Gram -ve coccus | 6.06 ± 0.023 | 3.07 ± 0.129 | 13.2 ± 0.88 | 8.66 ± 0.55 | 0.0808 ± 0.63 | 2.92 ± 1.2 | 1.7 ± 0.9 | 1.18 ± 0.32 | 1.87 ± 0.76 | 1.16 ± 0.34 | 0.66 ± 0.23 |

The growth of different bacterial isolates from the rhizospheres of Thar desert plants under drought, nutrient deficiency and extremes of pH and temperature are shown as number of cells/ ml. Luria-Bertani (LB) media, pH 7.0, were supplemented with 15% and 20% w/v PEG-6000 to mimic drought stress, while 1/4<sup>th</sup> and 1/6<sup>th</sup> LB were used to mimic nutrient deficiency conditions. The respective media were seeded with equal amount of primary inoculum of each isolate, grown overnight at 30 $^{\circ}$ C, and the absorbance was observed at 600 nm. The pH of the LB media was adjusted after autoclaving to 3, 9, and 11, to study growth under extremes of pH, and the cultures were incubated at 5 $^{\circ}$ C, 15 $^{\circ}$ C, 45 $^{\circ}$ C and 60 $^{\circ}$ C to study growth under extremes of temperature. The growth is presented as the observed cell density ( $10^8$ cells/ml) after overnight growth of each culture, starting from equal inoculum size. $A_{600\text{ nm}} = 1$ corresponds to $8 \times 10^8$ cells/ml.

Deduplication analysis of the 12 rhizobacterial isolates by digestion of the amplified 16S rDNA fragments with the four-base cutter HaeIII indicated six unique band patterns (Fig. S1). Sanger sequencing revealed three unique non-chimeric sequences, 16S rDNA genes of the strains C1P-IITJ, J4-IITJ and T7-IITJ, sharing a homology of 99.89%, 98.9% and 100% with *Enterobacter cloacae*, *Kalamiella piersonii*, and *Peribacillus frigoritolerans*, respectively. Based on their database homology, strains C1P-IITJ, J4-IITJ, and T7-IITJ were labelled as *Enterobacter cloacae* C1P-IITJ, *Kalamiella piersonii* J4-IITJ, and *Peribacillus frigoritolerans* T7-IITJ, respectively. The 16S rDNA sequences of *Enterobacter cloacae* C1P-IITJ, *Kalamiella piersonii* J4-IITJ, and *Peribacillus frigoritolerans* T7-IITJ were submitted to NCBI GenBank with accession numbers OQ991942, OQ992202, and OQ991918, respectively.

Biochemical characterization of C1P-IITJ, J4-IITJ, and T7-IITJ showed that all three bacteria hydrolyzed starch produced oxidase and utilized citrate to produce pyruvate and carbon dioxide (Fig. 2). C1P-IITJ could not produce cytochrome oxidase and showed a negative MR but a positive VP test. Similarly, J4-IITJ also resulted in negative oxidase activity, but it gave positive MR and VP tests. T7-IITJ produced cytochrome oxidase, but it exhibited negative MR and VP tests. Among the three isolates, J4-IITJ produced a maximum halo zone (SI 4.28) in the phosphate solubilization test, followed by T7-IITJ (3.86) and C1P-IITJ (3.15). In the EPS production test, C1P-IITJ produced the maximum EPS, i.e., 282.5 µg/ml, followed by J4-IITJ (233.75 µg/ml) and T7-IITJ (133.12 µg/ml) respectively. T7-IITJ produced the greatest amount of IAA, the plant hormone facilitating root growth, 42.61 µg/ml, then C1P-IITJ (34.15 µg/ml) and J4-IITJ (18.92 µg/ml).

**Fig. 2.**
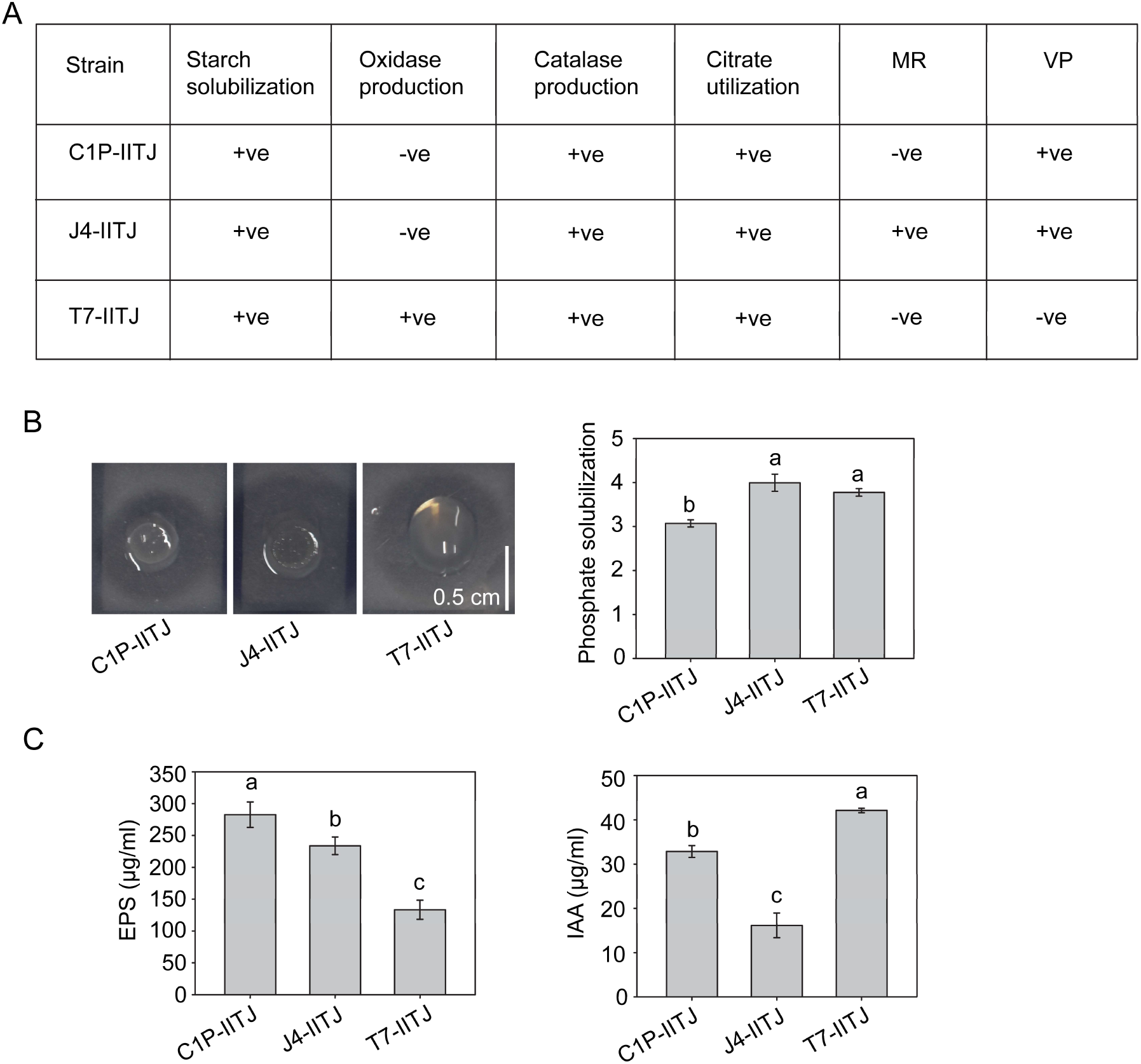
Biochemical characteristics and plant growth-promoting traits of three Thar desert rhizobacteria. **A** Some biochemical characteristics of the three rhizobacteria viz., *Enterobacter cloacae* C1P-IITJ, *Kalamiella piersonii* J4-IITJ, and *Peribacillus frigoritolerans* T7-IITJ, are shown. Starch solubilization, oxidase and catalase production, citrate utilization, methyl red (MR), and Voges-Proskauer (VP) test results are shown as positive (+ve) or negative (-ve) responses by each bacterium (see Methods). All experiments were repeated three times with similar results. **B** A plant growth-promoting trait, phosphate solubilization potential of three bacterial isolates, C1P-IITJ, J4-IITJ, and T7-IITJ, are shown. The bacterial cultures were spotted on phosphate solubilization media (see Methods), and halo formation around the colony was observed after seven days. The solubilization index measured by the (halo zone diameter + colony diameter)/ (colony diameter) of each bacterium is shown on the right. Bars indicate average solubilization indices with a standard error of six bacterial colonies. **C** Levels of exopolysaccharide (EPS) and indole acetic acid (IAA) produced by each bacterium are shown in µg/ml (see Methods). Different lower-case alphabets above the error bars represent significant differences (*P* < 0.05; Tukey’s test).

### Thar desert rhizobacteria promotes plant growth under non-stress and drought stress conditions

All three identified rhizobacteria, viz., C1P-IITJ, J4-IITJ, and T7-IITJ, promoted the growth of *A. thaliana* in one-month soil culture, as observed by a significant improvement in root length and rosette diameter (Fig. 3A, 3B). In hydroponic growth conditions using ¼^th^ Hoagland media alone (non-stress control) as well as 5% PEG-supplemented ¼^th^ Hoagland media (drought stress), both the shoot diameter and root length improved significantly within two weeks upon bacterial inoculation (Fig. 3C, 3D). Among the isolated rhizobacterial strains improving plant growth and stress tolerance, only T7-IITJ was considered for further analyses, as the other two isolates, *Enterobacter cloacae* C1P-IITJ and *Kalamiella piersonii* J4-IITJ were found to be potential human pathogens (Davin-Regli and Pagès, 2015; Rekha et al. 2020).

**Fig. 3.**
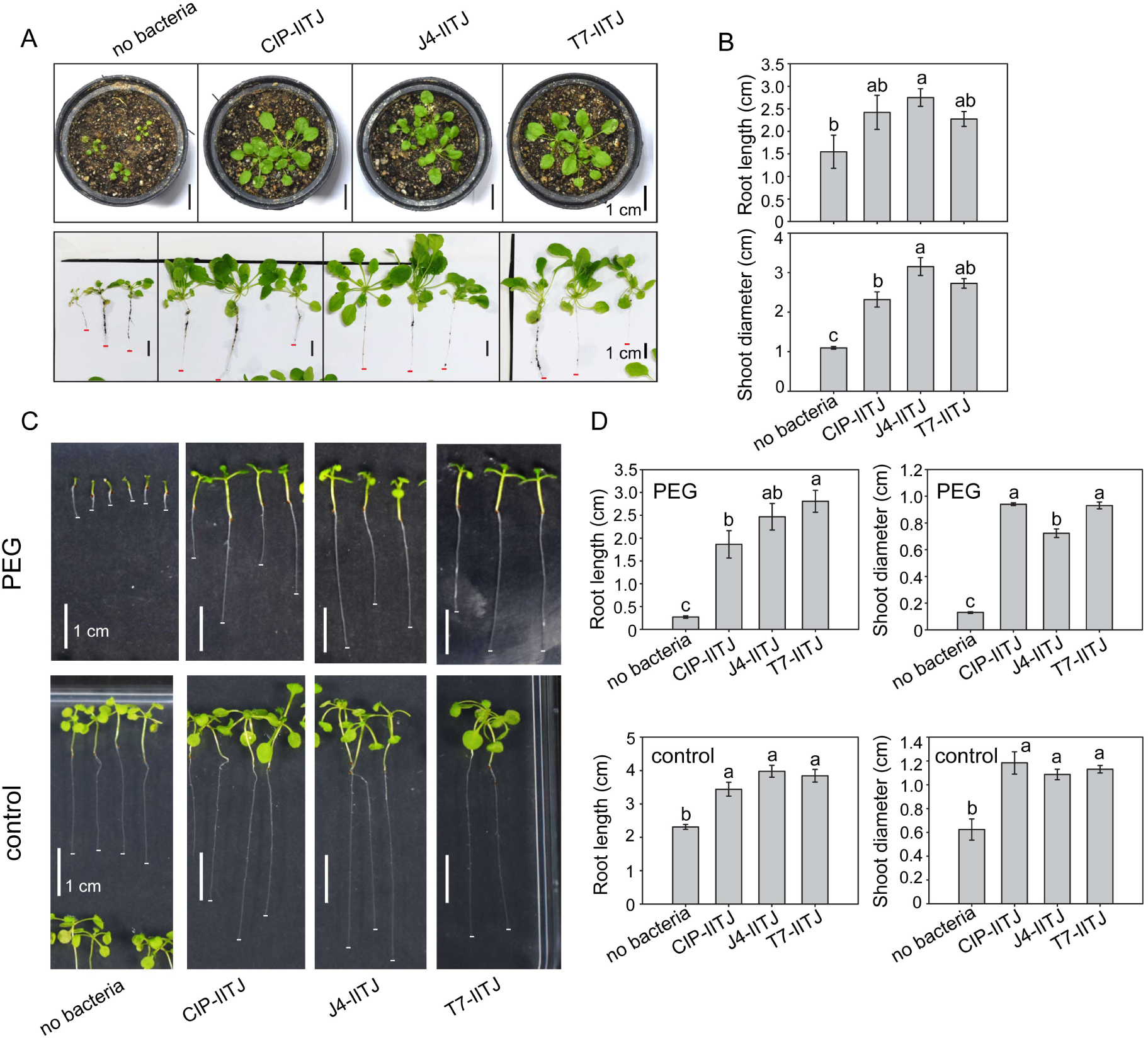
Promotion of plant growth and drought stress tolerance by three Thar desert rhizobacteria. **A** Growth promotion of *Arabidopsis thaliana* in soil, after inoculation by *Enterobacter cloacae* C1P-IITJ, *Kalamiella piersonii* J4-IITJ, and *Peribacillus frigoritolerans* T7-IITJ, is shown. Plant seeds were sown in soil mix and inoculated with bacteria after 10 d, and growth was observed after a month (see Methods). Plants were carefully removed from the soil, washed, and photographed. Black scale bars indicate 1 cm. **B** The difference in root length and shoot (rosette) diameter between the inoculated and non-inoculated plants in the soil culture (**A**) are shown. Bars indicate an average of six plants, with standard error. Asterisks indicate significant differences between inoculated and non-inoculated plants (*p* < 0.05; Student’s *t*-test). **C** Improvement of drought stress tolerance of *A. thaliana* under hydroponic growth conditions, without and with inoculation by C1P-IITJ, J4-IITJ, and T7-IITJ, after a day of sowing. The plants were grown in ¼^th^ Hoagland media, pH 5.8 (control), or media supplemented with 5% PEG-6000 for 14 d, transferred to solidified agar plates, and photographed. White scale bars indicate 1 cm. **D** The difference in root length and shoot diameter between the inoculated and non-inoculated plants in the hydroponic culture (**C**) are shown. Bars indicate an average of six plants, with standard error. Different lower-case alphabets above the error bars represent significant differences (*P* < 0.05; Tukey’s test).

### The genome of *Peribacillus frigoritolerans* T7-IITJ contains genes promoting the growth and stress tolerance of plants

T7-IITJ genome sequence, obtained through long read sequencing, was assembled into three contigs of 5375132 bp having 40.40% GC with N50 of 3677948 bp. Functional annotation of the genome revealed 7150 coding sequences, 39 rRNA, 83 tRNA, and one tmRNA gene sequence. The average nucleotide identity (ANI) among different *Peribacilli* was low, ∼67-99%, suggesting significant interspecific genomic variation (Table S3). A z-score of 0.99852 in the tetra correlation analysis confirmed our Sanger sequencing results that the strain T7-IITJ is a *Peribacillus frigoritolerans*, no other *Peribacillus* species (Table S4). Further, the ANI scores suggested that T7-IITJ is close to *Peribacillus frigoritolerans* CF13 in comparison to other members of the species (Table S3). While phylogenomic analysis showed that T7-IITJ shares the highest similarity with *Peribacillus frigoritolerans* HMB 20428, the genomic matrix also revealed that the genomes of all selected Peribacillus strains share only a few genes, suggesting considerable genomic variation between the strains, and the unique nature of T7-IITJ (Fig. 4A). The whole genome sequence of T7-IITJ was submitted to the NCBI with an accession number JAUPFL000000000. An *in-silico* analysis of the T7-IITJ genome revealed its plant growth-promoting potential, inferred from the occurrence of genes producing the phytohormone auxin, EPS, and those promoting nutrient mining from the soil (Ahmad *et al*. 2022). The T7-IITJ genome contained three genes for auxin biosynthesis, five for siderophore production, nine for nitrogen fixation and metabolism, five for phosphate solubilization, and other genes associated with EPS and antibiotic production (Table 2). We manually confirmed the absence of virulence genes for plant or human pathogenesis in the genome of this bacterium (Fig. 4B). The antiSMASH software predicted six gene clusters for the secondary metabolite biosynthesis within the genome of T7-IITJ, among which five clusters were found to have antipathogenic activity (Table 2). A non-ribosomal peptide synthase (NRPS) gene cluster showed a high 87% similarity with a known cluster encoding koranimine biosynthesis, a compound with nematicidal activity (Martho *et al*. 2016). The NRPS-independent, IucA/IucC-like siderophore cluster was found to have 50% similarity with the schizokinen biosynthesis cluster, encoding a siderophore with antifungal properties (Maindad et al. 2014). The beta lactone gene cluster was 40% similar to fengycin, another antifungal compound (Vanittanakom et al. 1986). An opine-like metallophore gene cluster found in T7-IITJ had 87% similarity with the antimicrobial compound bacillopaline-producing gene cluster (Bellotti, Rowińska-Żyrek and Remelli 2021). Lastly, the RRE element-containing cluster showed 60% similarity with that of an antimicrobial compound, paeninodin (Semenzato et al. 2022).

**Fig. 4.**
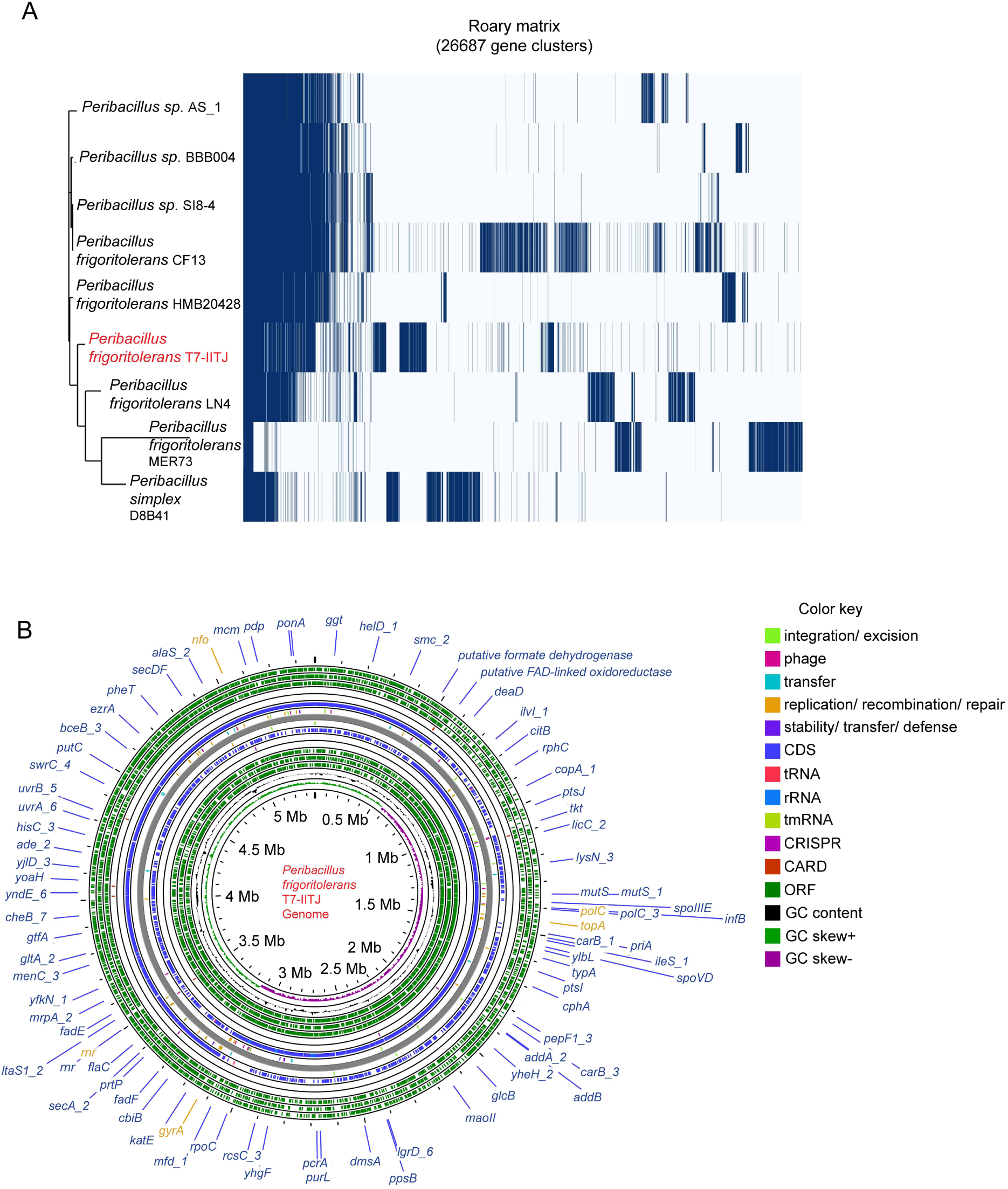
Genome analysis of *Peribacillus frigoritolerans* T7-IITJ. **A** Phylogenomic tree of *Peribacillus frigoritolerans* T7-IITJ, drawn using the FastTree v2.1.10 tool and Roary pipeline, based on nucleotide identity with closely related bacterial species (see Supplemental Tables 3, 4). **B** Genome map of *Peribacillus frigoritolerans* T7-IITJ. The genome map was drawn from the whole genome sequence of *P. frigoritolerans* using the Proksee tool and annotated using PROKKA. Antibiotic resistance genes were identified using the Comprehensive Antibiotic Resistance Database (CARD) identifier.

**Table 2.** *Peribacillus frigoritolerans* genes for the promotion of plant growth and stress tolerance.

| Protein | Function | Start (bp) | Stop (bp) |
| --- | --- | --- | --- |
| **Auxin biosynthesis** |  |  |  |
| Indole-3-glycerol phosphate synthase | Branchpoint of indole-3-acetic acid biosynthesis from the tryptophan biosynthetic pathway | 1587487 | 1586711 |
| Auxin efflux carrier family protein | Component of auxin efflux | 379587 | 380012 |
| Auxin efflux carrier family protein |  | 380048 | 380215 |
| **Siderophore biosynthesis** |  |  |  |
| Putative siderophore transport system ATP-binding protein YusV | Siderophore production | 2785410 | 2785820 |
| Siderophore biosynthesis diaminobutyrate-2-oxoglutarate aminotransferase |  | 3183640 | 3183987 |
| Siderophore biosynthesis diaminobutyrate-2-oxoglutarate aminotransferase |  | 3184008 | 3184478 |
| Siderophore biosynthesis L-2,4-diaminobutyrate decarboxylase |  | 3184486 | 3185994 |
| Siderophore transport protein | Siderophore transport from the cell | 4407281 | 4408513 |
| **Nitrogen fixation** |  |  |  |
| Nitrate transporter | Nitrate/nitrite assimilation | 517739 | 518656 |
| Nitrite reductase [NAD(P)H] | Reduction of nitrate to nitrite via a two-electron transfer | 518707 | 520011 |
| Nitrite reductase [NAD(P)H] |  | 520077 | 520409 |
| Assimilatory nitrite reductase [NAD(P)H] small subunit | Rate-limiting reduction of nitrate to nitrite | 520406 | 521071 |
| Nitrate/nitrite transporter NarT | Transport | 881313 | 881053 |
| Nitrogen regulatory PII-like protein | Regulation of nitrogen fixation | 3102353 | 3102024 |
| Nitric oxide reductase activation protein NorQ | Denitrification | 3815270 | 3816136 |
| Nitric oxide reductase activation protein NorD | Conversion of nitric oxide to nitrous oxide | 3816582 | 3817571 |
| Nitric oxide reductase activation protein NorD |  | 3817801 | 3818064 |
| **Phosphate solubilization** |  |  |  |
| Phosphate transport system regulatory protein PhoU | Phosphate homeostasis | 5043318 | 5043971 |
| Phosphate regulon transcriptional regulatory protein PhoB (SphR) | Positive regulator for the phosphate regulon | 2016776 | 2016663 |
| Phosphate ABC transporter, ATP-binding protein PstB | Phosphate transport | 5042471 | 5043298 |
| Phosphate ABC transporter, permease protein PstC |  | 5040636 | 5041574 |
| Phosphate ABC transporter, permease protein PstA |  | 5041613 | 5042455 |
| **Exopolysaccharides (EPS) production** |  |  |  |
| Peptidoglycan glycosyltransferase FtsW | Peptidoglycan polymerase promoting exopolysaccharide synthesis | 1532648 | 1532472 |
| Peptidoglycan glycosyltransferase<br>FtsW |  | 1533130 | 1532651 |
| Peptidoglycan glycosyltransferase<br>FtsW |  | 1533570 | 1533193 |
| **Gene clusters with anti-pathogenic activity** |  |  |  |
| Non-ribosomal peptide synthetase | Nematicidal activity | 1707072 | 1757675 |
| NRPS-independent, IucA/IucC-like<br>siderophore | Antifungal activity | 955668 | 967468 |
| Betalactone |  | 3449676 | 3473843 |
| Opine-like metallophore | Antimicrobial activity | 3409397 | 3431536 |
| RRE-element containing cluster |  | 768683 | 788967 |

Proteins encoded by the *Peribacillus frigoritolerans* genome, which function in plant growth promotion, are shown. The proteins were categorized based on their functions, viz, auxin biosynthesis, siderophore biosynthesis, nitrogen fixation, phosphate solubilization, exopolysaccharide production, and resistance against antibiotics. The physical positions of the corresponding genes on the genome are shown by the starting and ending nucleotide positions (in base pairs). The genes were functionally annotated using Prokka 1.12. J-species software. Gene clusters for the biosynthesis of secondary metabolites having anti-pathogenic activity were predicted using the antiSMASH v. 7.0.1 software.

Taking cues from the T7-IITJ genome, we carried out additional biochemical tests to detect this bacterium’s nitrogen fixation and siderophore production abilities, which are vital for its biofertilization potential. This bacterium produced 1.6 µg/ml of nitrite, indicating its nitrate reductase activity. When patched on CAS agar, T7-IITJ produced orange halo zones of 24 mm around its colonies due to the chelation of iron by siderophores from the iron-CAS/HDTMA blue dye complex of the media. In addition, T7-IITJ produced clear zones of growth inhibition of two fungal plant pathogens, *Fusarium oxysporum* and *Rhizoctonia solani*, around its colony. The growth inhibition zone of 18 ± 0.58 mm and 25 mm was observed against *Rhizoctonia solani* and *Fusarium oxysporum*, respectively (Fig. S2).

### *Peribacillus frigoritolerans* T7-IITJ increases chlorophyll content and protects Arabidopsis tissues from desiccation and oxidative stress

We observed a significant 2.5-2.8-fold increase in chlorophyll a, b, and total chlorophyll in *A. thaliana* seedlings inoculated with T7-IITJ compared to non-inoculated seedlings under PEG-induced drought, which is indicative of higher photosynthetic functions leading to an increase in growth under stress. Chlorophyll contents decreased under drought than non-stressed control. In control samples too, chlorophyll a, and total chlorophyll increased slightly but significantly by T7-IITJ (Fig. 5A). The leaf proline levels increased by a significant 2.5-fold due to T7-IITJ which suggests that the bacterium triggered osmolyte biosynthesis to protect the tissues from drought. Proline levels were higher under drought. A significant increase in proline due to T7-IITJ inoculation was also found under control conditions (Fig. 5B). On the other hand, tissue ROS, viz., H_2_O_2,_ was reduced by half in seedlings due to T7-IITJ inoculation under drought and by 5-fold under control conditions (Fig. 5C). The levels of O_2_.^-^ were significantly reduced 4.7-fold under drought and by half under control conditions due to T7-IITJ inoculation (Fig. 5D). MDA levels, indicative of the extent of membrane lipid peroxidation, were significantly reduced 5-fold in the inoculated plants under PEG and by 2-fold under control conditions (Fig. 5E). This suggests that T7-IITJ improves the protection of plant tissues from oxidative damage under drought as well as control conditions.

**Fig. 5.**
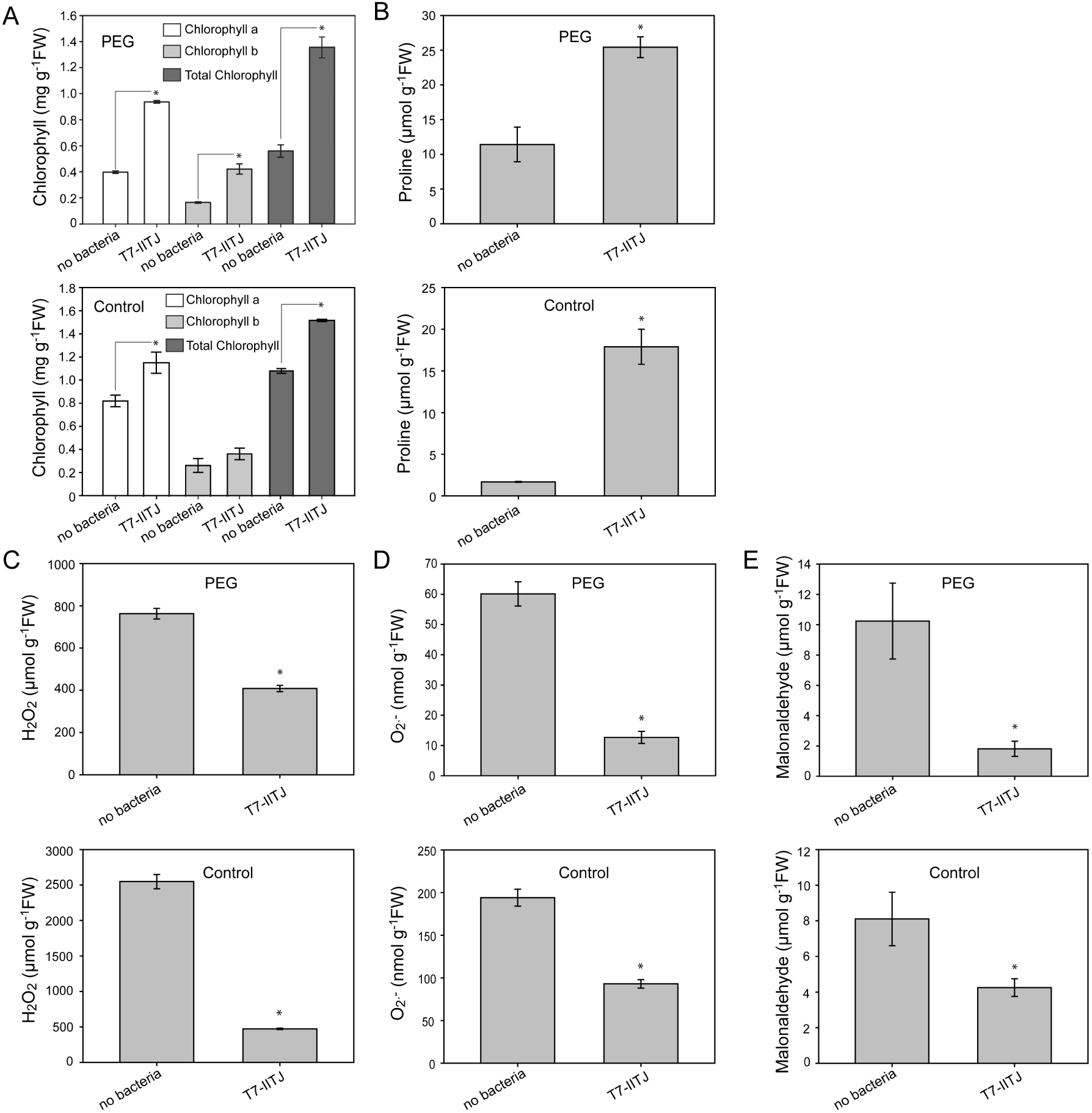
Chlorophyll, proline, and reactive oxygen species content in Arabidopsis seedlings upon *Peribacillus frigoritolerans* T7-IITJ inoculation. One-week-old *Arabidopsis thaliana* seedlings grown in ¼^th^ Hoagland media, pH 5.8, were inoculated with *Peribacillus frigoritolerans* T7-IITJ, allowed to interact for another week, and treated with 5% PEG-6000 for five days. Some seedlings were not inoculated with bacteria but subjected to the same PEG treatment. Bacteria-inoculated and non-inoculated seedlings were grown in ¼^th^ Hoagland media without PEG for a similar duration for the control experiments. After that, the shoots were harvested and weighed, homogenized using mortar and pestle, and extracted with acetone and sulfosalicylic acid to measure the chlorophyll (**A**) and proline contents (**B**), respectively, by spectrophotometry (for details, see Methods). Reactive oxygen species, viz. hydrogen peroxide (H_2_O_2_) (**C**) and superoxide (O_2_^.-^) (**D**), and malonaldehyde (**E)** levels were measured in whole seedlings by specific biochemical tests using spectrophotometry (see Methods). The quantity of each measured parameter was normalized by grams of the sample fresh weight (FW). The bars indicate average values with a standard error of three biological replicates, each comprising ten seedlings. Asterisks indicate significant differences between the bacteria-inoculated and non-inoculated seedling samples (*P* < 0.05; Student’s *t*-test).

### *Peribacillus frigoritolerans* T7-IITJ increases total nitrogen, iron, and phosphate content of Arabidopsis tissues

As T7-IITJ was found to solubilize nitrogen, iron, and phosphate, we were intrigued to explore whether the levels of these nutrients in *A. thaliana* tissues altered upon T7-IITJ inoculation. The total tissue nitrogen increased 4-fold under drought and 3-fold under control conditions due to T7-IITJ inoculation. Total tissue iron, ferrous (Fe^2+^), and ferric (Fe^3+^) also increased twice under drought conditions due to T7-IITJ. On the other hand, phosphate levels increased by a significant 1.6-fold in plant tissues under drought (Fig. 6). These results indicated that T7-IITJ enhanced the mining of essential nutrients in plants to boost growth under control and drought conditions.

**Fig. 6.**
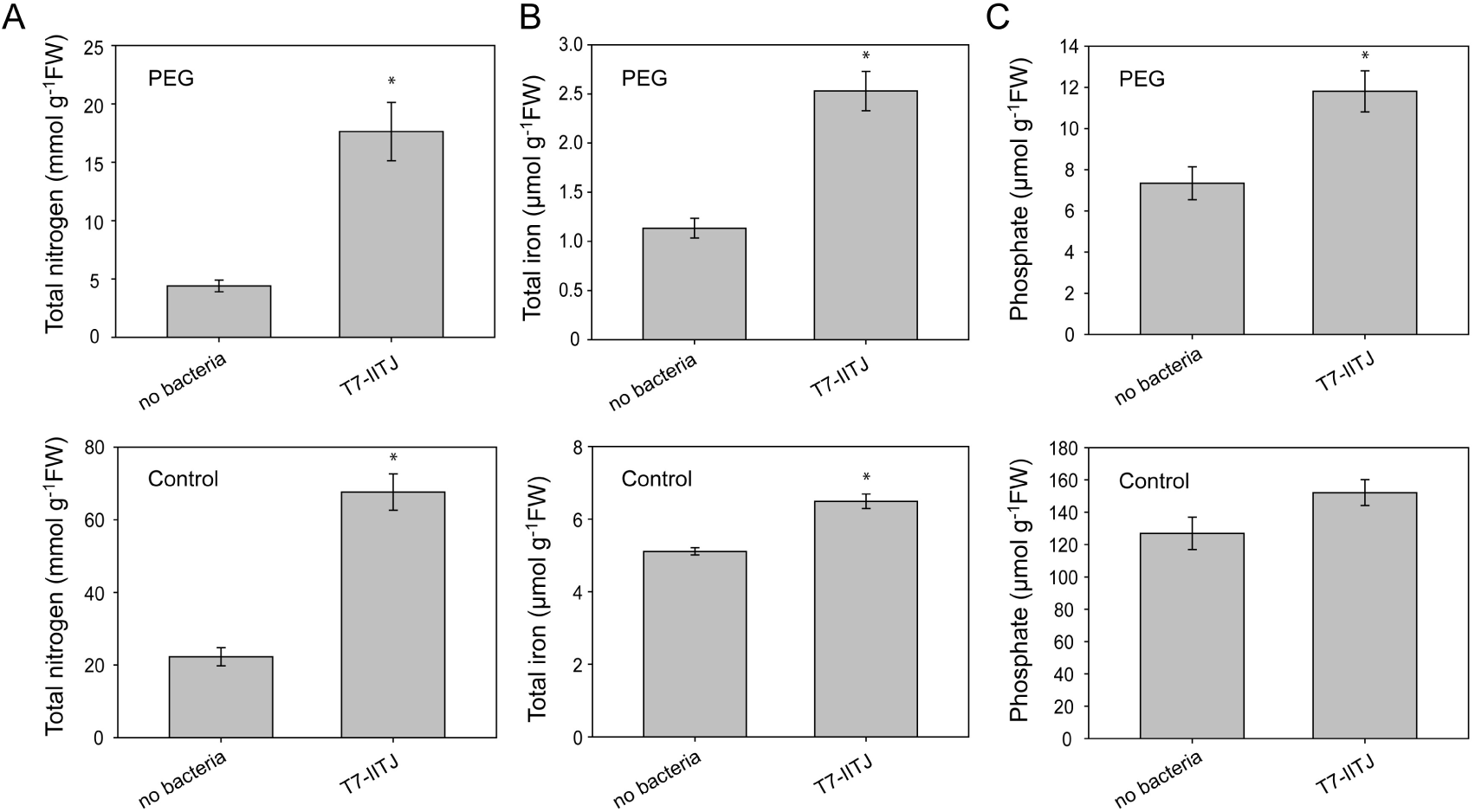
Nutrient content in Arabidopsis seedlings upon *Peribacillus frigoritolerans* T7-IITJ inoculation. One-week-old *Arabidopsis thaliana* seedlings grown in ¼^th^ Hoagland media, pH 5.8, were inoculated with *Peribacillus frigoritolerans* T7-IITJ, allowed to interact for another week, and treated with 5% PEG-6000 for five days. Some seedlings were not inoculated with bacteria but subjected to the same PEG treatment. Bacteria-inoculated and non-inoculated seedlings were grown in ¼^th^ Hoagland media without PEG for a similar duration for the control experiments. After that, the whole seedlings were harvested, weighed, and homogenized using mortar and pestle, to measure the total tissue nitrogen (**A**), total iron (Fe^II^ and Fe^III^) (**B**), and phosphate contents (**C**), respectively, by spectrophotometry (see Methods). The quantity of each measured parameter was normalized by grams of the sample fresh weight (FW). The bars indicate average values with a standard error of three biological replicates, each comprising ten seedlings. Asterisks indicate significant differences between the bacteria-inoculated and non-inoculated seedling samples (*P* < 0.05; Student’s *t*-test).

### *Peribacillus frigoritolerans* T7-IITJ alters global gene expression in Arabidopsis seedlings under drought stress

To gain further insights into the molecular mechanisms in the bacteria-inoculated plants leading to increased growth and stress tolerance, we compared the mRNA levels of inoculated versus non-inoculated plants treated with PEG by RNA sequencing. Transcriptome analysis of two-weeks-old *A. thaliana* seedlings inoculated with T7-IITJ in PEG-containing hydroponic media indicated the upregulation of 445 genes (log_2_FC > 1) and downregulation of 503 genes (log_2_FC < -1), *FDR* < 0.05 (Table S5). Twenty-six protein-coding genes were most strongly upregulated with log_2_FC > 2.5 (Table 3). These included two vacuolar-membrane-localized iron and manganese transporter genes, *VTL1* and *VTL5*, whose paralogs supply iron to beneficial bacteria (Walton *et al*. 2020). T7-IITJ induced several genes encoding defense-related proteins, viz., *beta-galactosidase 7* (*BGAL7*; *AT5G20710*) (Hrubá *et al*. 2005; Thatcher *et al*. 2015), contributing to root development (Hrubá *et al*. 2005), *jacalin-like lectin domain-containing protein* (*JAL10*; *AT1G52070*) (Häffner, Konietzki and Diederichsen 2015) of the jasmonate (JA)-signaling pathway, and a defense-related aquaporin *NOD26-LIKE INTRINSIC PROTEIN 1;1* (*NLM1*; *AT4G19030*) involved in H_2_O_2_ signaling (Sadhukhan *et al*. 2017). Genes encoding hybrid proline-rich protein (HyPRP) superfamily proteins *AZL14* and *AZI5* were also strongly induced by T7-IITJ, which helps colonize beneficial bacteria in the root, stimulates root growth, and induces systemic resistance against pathogens. The *azl14* mutant inoculated with *Pseudomonas simiae* showed reduced root length and less colonization of the strain in roots (Banday *et al*. 2022). *Xyloglucan endotransglucosylase/hydrolase 12*, induced by T7-IITJ, is a protein expressed in *A. thaliana* roots and is involved in root hair development (Song et al. 2016). Another gene induced by T7-IITJ, *phosphate starvation-induced glycerol-3-phosphate permease gene/ root hair specific 15* (*G3PP2*/ *RHS15*), can help in root development and more efficient phosphate mining (Ramaiah *et al*. 2011). *SMALL AUXIN UPREGULATED RNA 69* (*SAUR69*), belonging to the gene family involved in root development through auxin and other phytohormone-mediated signaling (Stortenbeker and Bemer 2019), was also induced by T7-IITJ. The strong upregulation of genes involved in root development in the transcriptome can explain the root elongation phenotype in the presence of T7-IITJ under drought stress. Seven of the 26 strongly induced genes were induced by drought and salt stresses in the public gene expression data, while the others were not typical drought-inducible genes (Table 3). We confirmed the expression levels of the transcriptome’s six most significantly induced genes (log_2_FC > 2.5; *FDR* < 10^-15^) in T7-IITJ-inoculated *A. thaliana* seedlings by qPCR. We conducted transcriptome analysis in whole seedlings but checked the expression levels separately in shoots and roots by qPCR. We found significant induction of the six genes in roots. Only *AT5G46890* was induced in the shoots as well (Fig. 6).

**Table 3.**
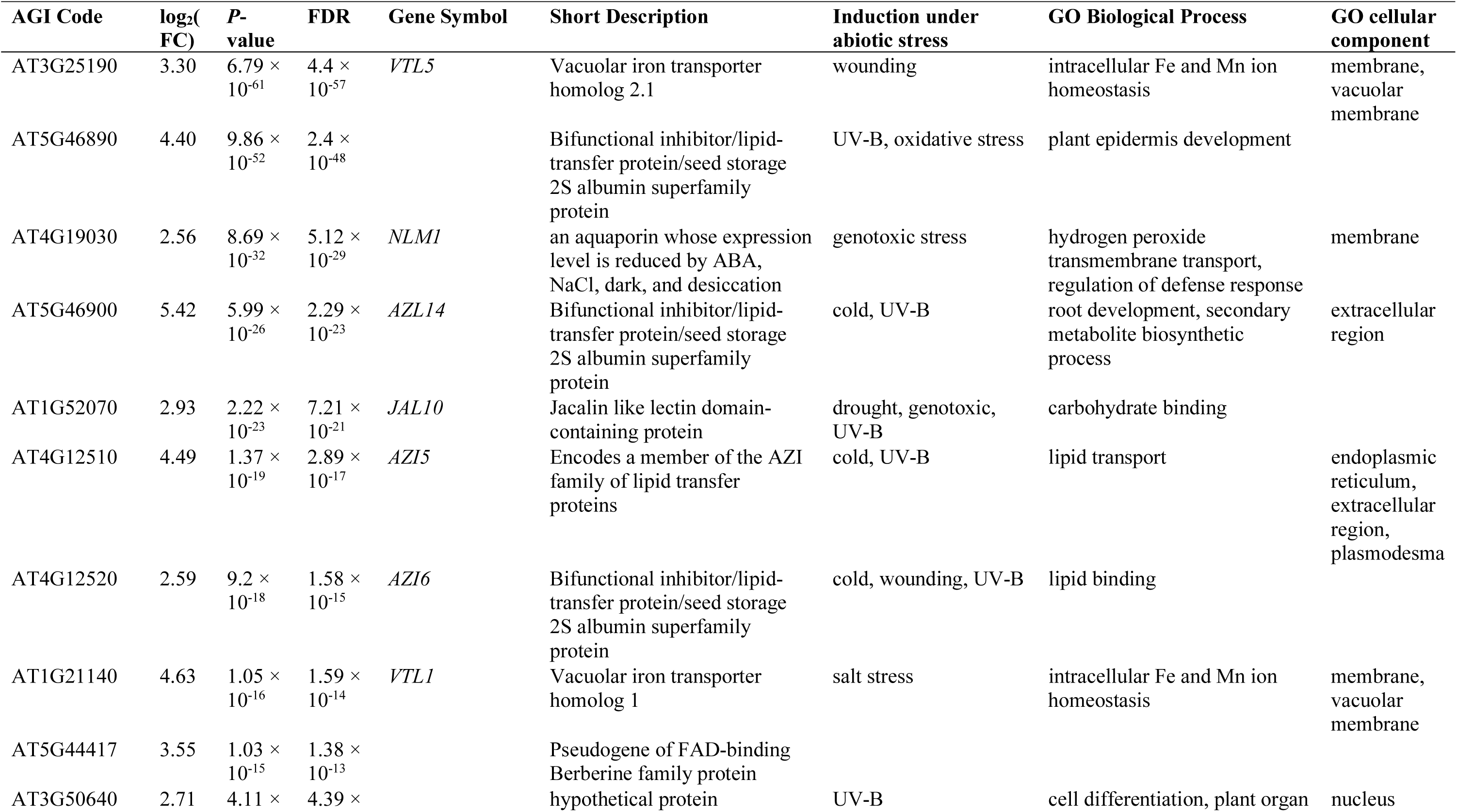
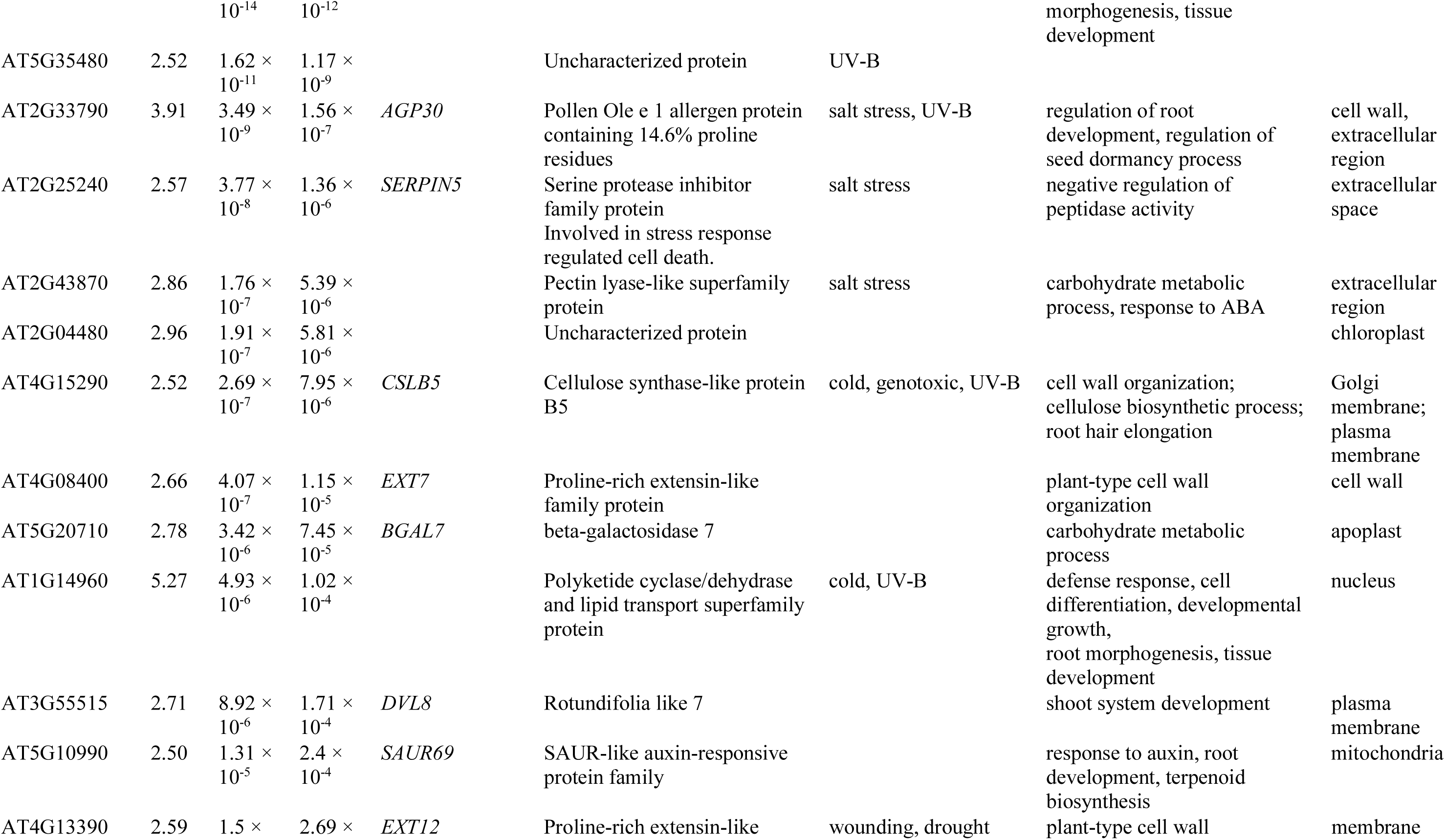
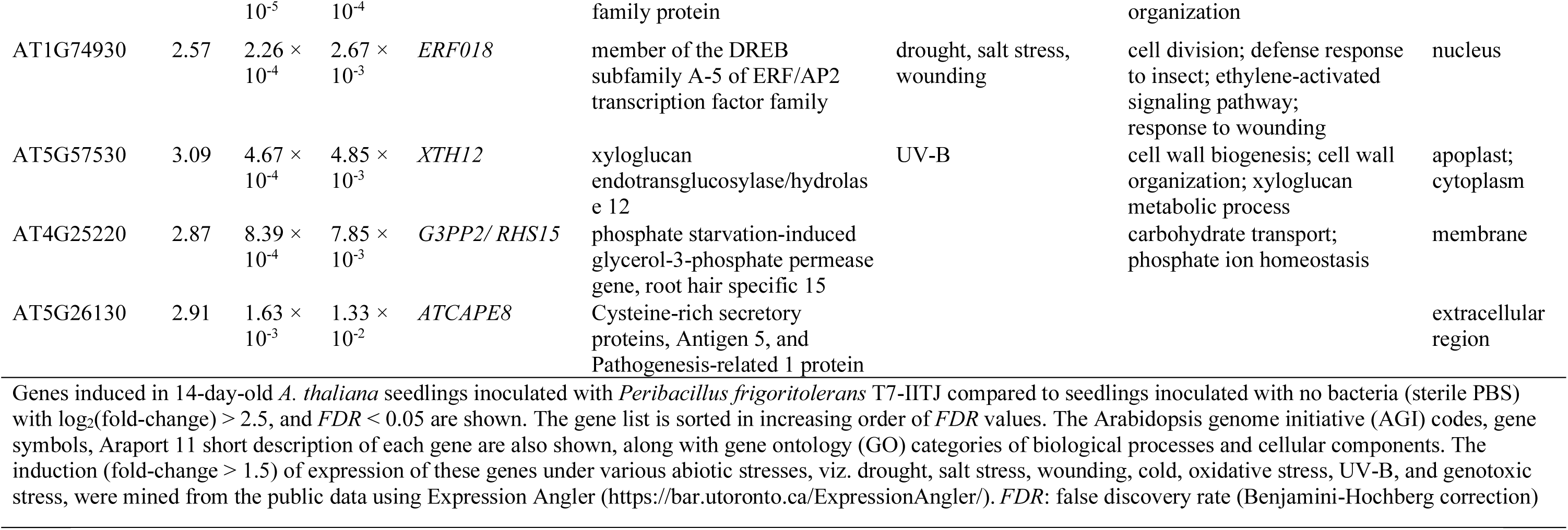
Genes highly induced in *Arabidopsis thaliana* seedlings inoculated with Peribacillus frigoritolerans T7-IITJ.

We conducted an enrichment analysis of KEGG pathways and GO categories to understand the functions of the induced genes. Photosynthesis, light harvesting, and electron transport in photosystem I and II, phylloquinone biosynthesis, carbon metabolism, iron homeostasis, and plant hormones auxin and salicylic acid (SA) were the enriched biological processes in the 445 genes induced by T7-IITJ. Enrichment of intracellular compartments like chloroplast, thylakoid, and photosystem light-harvesting complexes again indicated the importance of photosynthesis-related processes in improving plant growth by T7-IITJ (Table 4, Table S6). In addition, several secondary metabolite biosynthesis pathways were enriched among the induced genes. These included phylloquinol, terpenoid-quinone, trans-farnesyl diphosphate, geranylgeranyl diphosphate via mevalonate (MEV) and methylerythritol 4-phosphate (MEP), and polyisoprenoid biosynthesis (Table 4).

**Table 4.**
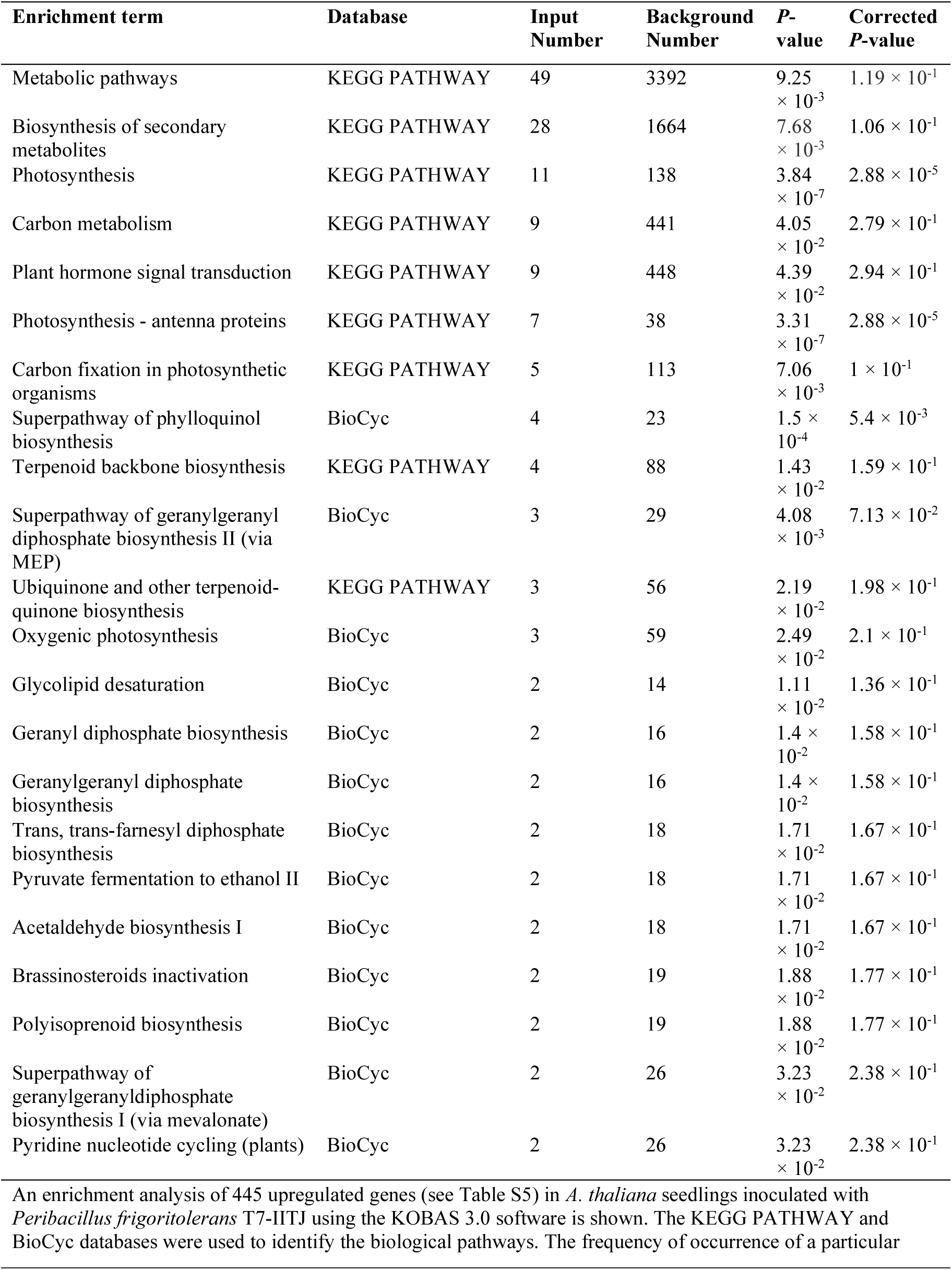
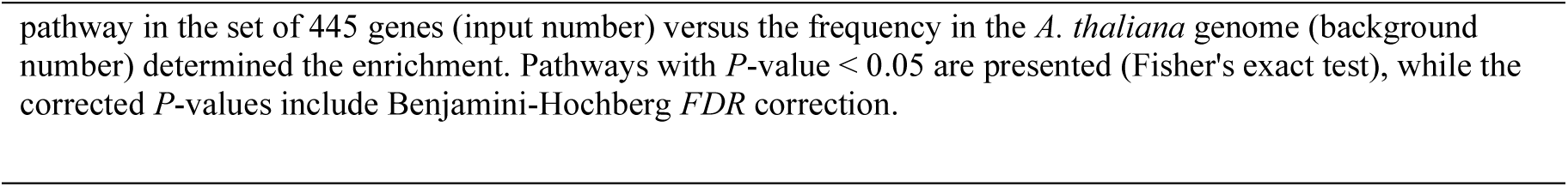
Biological pathways induced in *Arabidopsis thaliana* seedlings inoculated with *Peribacillus frigoritolerans* T7-IITJ.

A gene co-expression network analysis was conducted to interpret further the functions of the highly induced genes (log_2_FC > 2.5) from our transcriptome and to confirm if they worked together in common biological processes leading to the enhanced drought tolerance in *A. thaliana* upon T7-IITJ inoculation (Fig. 7). Interestingly the most strongly induced genes were highly interconnected through co-expression networks. A total of 18 most strongly induced genes (Table 3) were connected to nine additional genes with related functions. Furthermore, co-expressed genes shared functional protein domains in three network clusters. Functional enrichment analysis of the genes in the network indicated that iron and manganese homeostasis were the significantly enriched biological processes. Involved in these processes, *VTL1* and *VTL5* connected with four additional paralogs, *VIT1* (*AT2G01770*), *VTL2* (*AT1G76800*), *VIT homolog 3* (*AT3G43630*) and *VIT homolog 4* (*AT3G43660*).

**Fig. 7.**
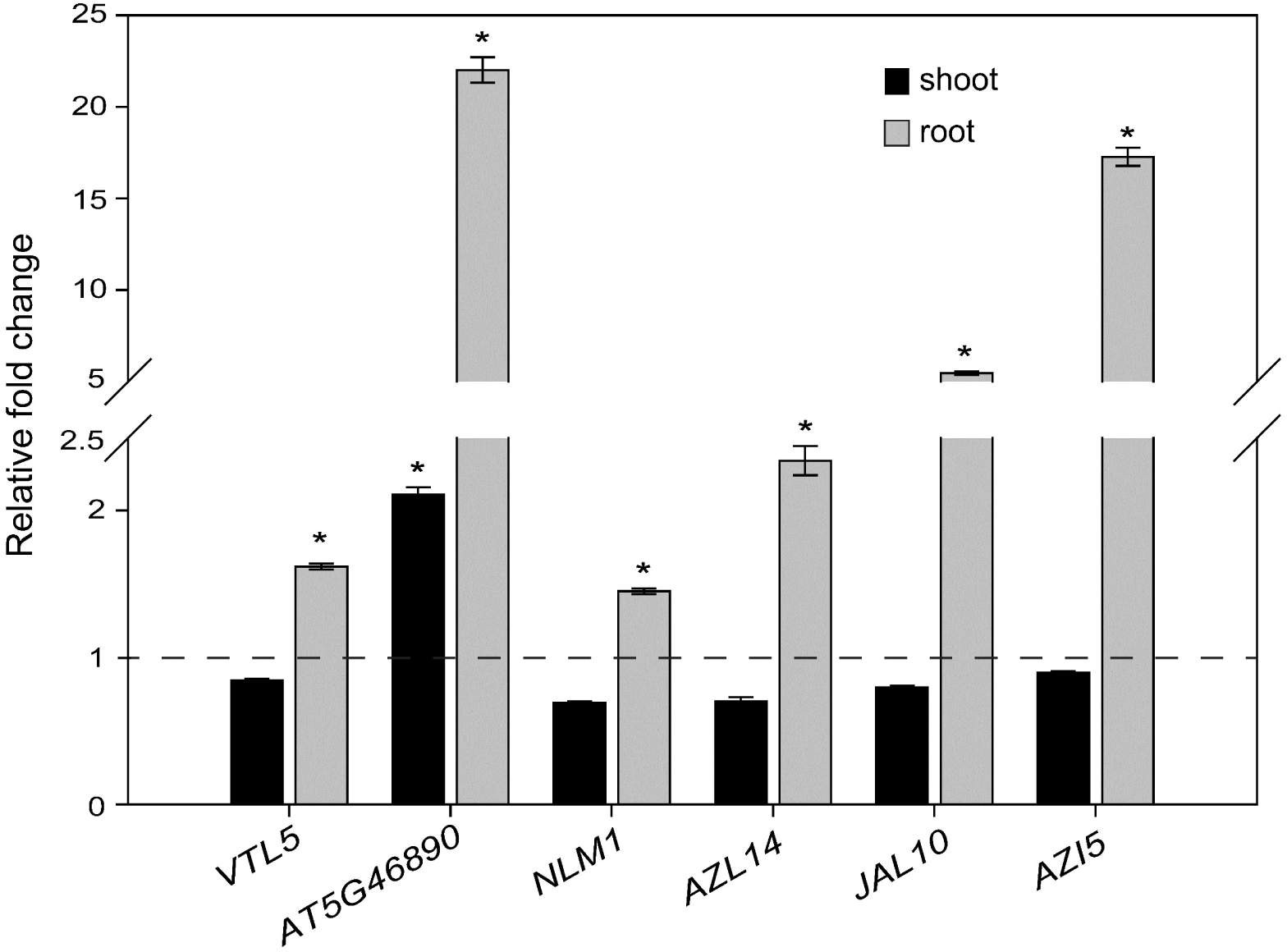
Real-time quantitative PCR of genes highly induced in Arabidopsis seedlings by *Peribacillus frigoritolerans* T7-IITJ under drought. One-week-old *Arabidopsis thaliana* seedlings were allowed to interact with *Peribacillus frigoritolerans* T7-IITJ for another week and treated with 5% PEG-6000 for five days. After that, the shoots and roots were separately harvested to isolate RNA and convert it to cDNA. Real-time PCR was conducted with the cDNA, and relative expression levels were estimated using the standard curve method (see Methods). The relative fold-change in expression levels of the top six highly induced genes in the transcriptome analysis (Table 3), viz., *VTL5*, *AT5G46890*, *NLM1*, *JAL1*, and *AZI5*, between T7-IITJ-inoculated versus non-inoculated control *A. thaliana* seedling samples are shown in the graph. A broken line shows the control levels. The expression levels in each sample were further normalized using *AtUBQ1* as the housekeeping gene internal standard. The bars indicate an average of three biological replicates, each comprising ten seedlings, with standard error. Asterisks indicate significant induction in gene expression between T7-IITJ-inoculated versus non-inoculated control (*P* < 0.05, Student’s *t*-test).

**Fig. 8.**
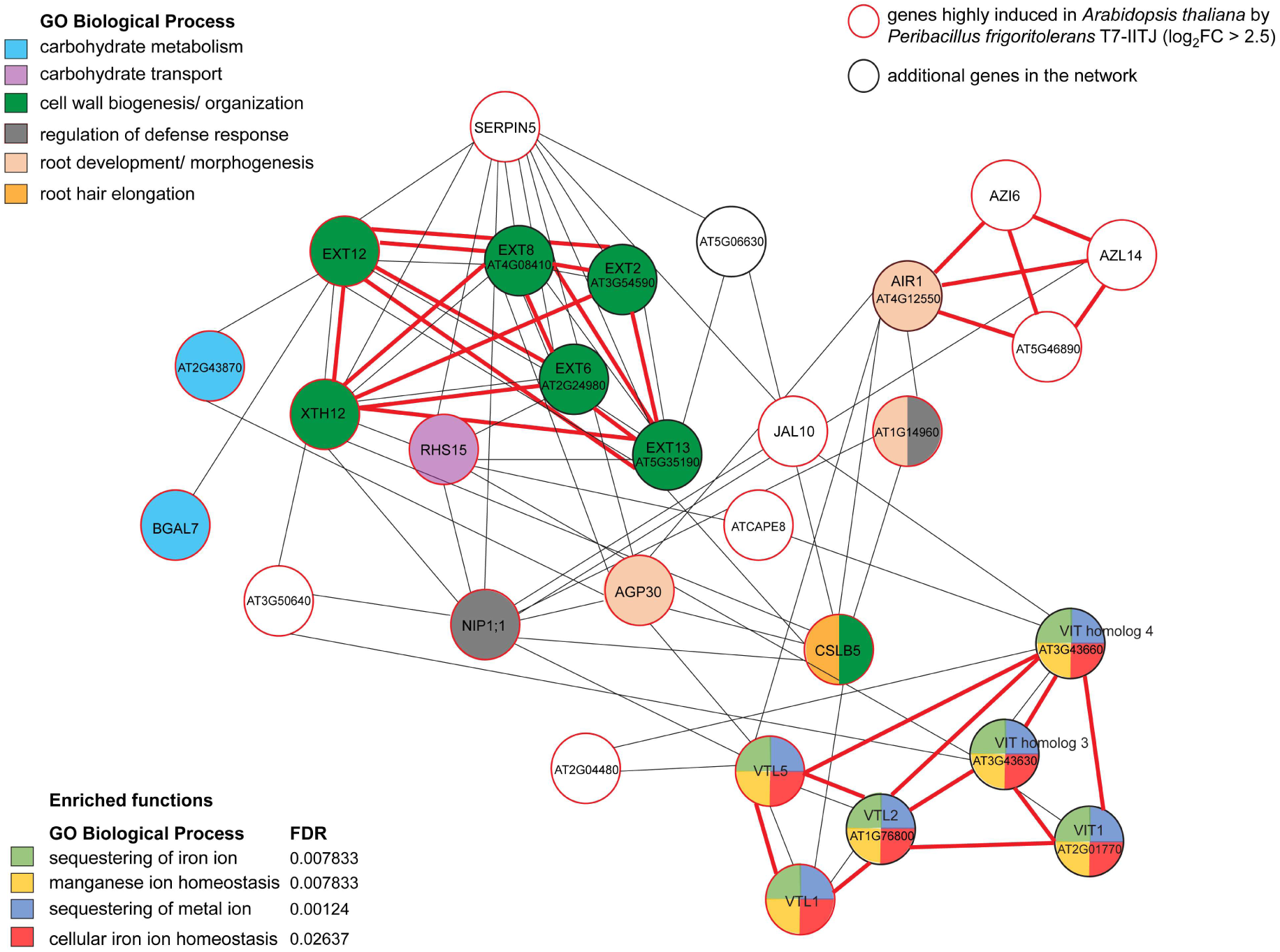
Co-expression network of genes highly induced in Arabidopsis seedlings by *Peribacillus frigoritolerans* T7-IITJ under drought. Genes induced more than the threshold of log_2_(fold-change) > 2.5, *FDR* < 0.05, in the transcriptome analysis (Table 3) were used for gene co-expression network construction using GENEMANIA web application (https://genemania.org/). Circles depict the network nodes and denote genes with AGI codes/ gene symbols (Table 3). The genes induced in our transcriptome analysis are circled in red, and additional genes in the network are circled in black. Lines denote the edges of the network. The black lines indicate co-expression, and the red lines indicate shared protein domains. The colored circles indicate the GO biological processes categorization of each gene. The enriched gene functions detected by the application are shown at the bottom. Enrichment was calculated by the frequency of occurrence within the 26 highly induced genes in our transcriptome versus the occurrence within the entire genome (*FDR* < 0.05). GO: gene ontology; *FDR*: false discovery rate (Benjamini-Hochberg correction).

Proline-rich extensin-like family proteins *EXT7, EXT12,* and another gene with a shared protein domain, *XTH12* were co-expressed with other paralogs *AT3G54590* (*EXT2*), *AT2G24980* (*EXT6*), *AT4G08410* (*EXT8*), and *AT5G35190* (*EXT13*), and functioned in cell wall biogenesis and organization. These genes were additionally connected to genes for carbohydrate metabolism and transport, *AT2G43870*, *BGAL7*, and *RHS15*, which were highly induced in our transcriptome. *AZI5*, *AZL14*, and *AT5G46890*, induced in our transcriptome, shared protein domains with *AT4G12550* (*AIR1*), which function in root development via auxin signaling and plant defense responses. Interestingly, *AIR1* co-expressed with the aquaporin *NLM1*, the proline-rich arabinogalactan protein *AGP30*, and *AT1G14960* induced in our transcriptome, which again functions in root development and defense responses. Another gene, *cellulose synthase-like protein B5* (*CSLB5*), working in both carbohydrate metabolism and root hair elongation, was interconnected to the other root development/ auxin signaling-related genes. The network analysis suggested possible mechanisms of growth promotion by T7-IITJ through the synergistic action of induced gene families, which included enhancement of iron and manganese uptake and vacuolar sequestration, root development through auxin signaling, and carbohydrate metabolism and cell wall biogenesis essential for the maintenance of growth processes. Expanding the network analysis to all upregulated genes (log_2_FC > 1, *FDR* < 0.05) based on predicted protein-protein interactions unraveled a more extensive network (Fig. S3). KEGG signaling pathway and GO categorization analysis of all network genes showed enrichment of photosynthesis, photosynthesis - antenna proteins, ROS biosynthesis, and metabolic pathways. We could identify seven hub genes for the network, viz., *PSAF*, *PSAK*, *LHCA1*, *PSAN*, *LHCB2.2*, *PSBQ1*, and *Lhb1B1*, all encoding proteins involved in photosynthesis.

Lastly, we were intrigued to examine the enriched biological processes in the 503 genes downregulated by T7-IITJ. The KEGG pathway enrichment analysis indicated that a subset of genes for the biosynthesis of secondary metabolites, metabolism of specific amino acids, phenylpropanoid biosynthesis, plant hormone and MAPK signaling pathways, and protein processing in the endoplasmic reticulum were suppressed by T7-IITJ (Table S7). Surprisingly, the enriched GO categories indicated that the expression of many genes functioning in response to ABA and to several stresses, including water deprivation, osmotic stress, salt stress, cold, hypoxia, and oxidative stress, were suppressed by T7-IITJ (Table S8). A predicted PPI network for downregulated genes (log_2_FC < -1, *FDR* < 0.05) indicated six hub genes, viz., *ABI5*, *LTI65*, *RD29A, PP2CA, ABI2,* and *PXG3*. KEGG pathway analysis showed the enrichment of secondary metabolite biosynthesis, hormone signal transduction, MAPK signaling, starch and sucrose metabolism, and metabolic pathways for the downregulated gene PPI network. The GO analysis showed significant enrichment of the response to ABA, abiotic stimulus, osmotic stress, and temperature stimulus (Fig. S3). The enrichment of metabolic pathways and response to light stimulus were common for both up and downregulated gene networks.

## Discussion

The plants in the Thar desert are naturally adapted to different abiotic stresses, including drought, nutrient deficiency, heat, high pH, etc. We have undertaken this study to understand the possible roles of PGPRs in this adaptation and to bioprospect strains for application as biofertilizers in arid agriculture. We started by culturing 30 bacterial strains from the rhizosoil of some Thar desert plants. We narrowed it down to the 12 most stress-tolerant pure cultures, further reduced to three after deduplication and 16S rDNA fragment sequencing analysis. All three strains significantly improved growth and drought tolerance in *A. thaliana* (Fig. 3). While two out of the three identified strains were found to be human pathogens and risked possible incorporation in the food chain if used as a biofertilizer, we focused our further study on the safest strain, T7-IITJ, identified as *Peribacillus frigoritolerans* basonym *Brevibacterium frigoritolerans*, with no virulence genes (Montecillo and Bae 2022a). T7-IITJ showed many plant growth- and drought-tolerance-promoting traits, including auxin production, phosphate solubilization, and EPS production (Fig. 2). Previously, *B. frigoritolerans* strains have been isolated from drought- and salt-stressed rhizospheres, as well as from the extreme cold environments of Antarctica (Zhang *et al*. 2019; Choe, Lee and Kim 2022). T7-IITJ increased the early seedling root growth of its source plant, *T. purpurea*, as well as the germination rates of two arid-region crops, *Triticum aestivum,* and *Setaria italica*, in both control and PEG-supplemented media (Fig. S3-5), suggesting its role in the natural germination and seedling growth of arid-region plants, especially during a lack of moisture. Microbial inoculation promotes seed germination in normal and stressed conditions via increasing the vigor index and accumulating antioxidant enzymes under stress, among other mechanisms. (Cardarelli *et al*. 2022). In PEG-supplemented hydroponic media, T7-IITJ led to a drastic increase in the root length of *A. thaliana* seedlings, which could be partly due to auxin biosynthesis by the bacterium (Fig. 2, 3). T7-IITJ increased the plant’s leaf chlorophyll and proline content under drought and decreased the ROS content of the plant tissues (Fig. 4), which are the properties of many PGPR bacilli (Abiala *et al*. 2023). The proline content increased, and ROS levels reduced under non-stressful conditions as well, due to T7-IITJ, also observed in the case of a few other PGPRs (Bano and Muqarab 2017; Khan and Bano 2019). Improvement of plant growth could also be attributed to the enhanced nutrient mining conferred by T7-IITJ, reflected in the tissue nitrogen, iron, and phosphate contents under drought and control conditions (Fig. 6).

To improve the performance of PGPRs, knowledge of their genetic makeup is essential, which can lead to further improvement of their biofertilization or biocontrol potential by genetic engineering (Koby *et al*. 1994) and genome editing (Elmore *et al*. 2023). Hence, we sequenced the entire genome of T7-IITJ and functionally annotated the genes *in silico*. The occurrence of auxin biosynthesis genes, genes encoding phosphate solubilization, nitrogen fixation, siderophore biosynthesis, and EPS production in its genome could be correlated with the results of the biochemical tests (Fig. 2, Table 2). The T7-IITJ genome also contains genes that can make nitrogen and iron available to desert plants, which was validated by our further biochemical experiments (Fig. S2A, B). The observed antagonistic effect of T7-IITJ against the plant pathogens *Rhizoctonia solani* and *Fusarium oxysporum* (Fig. S2), along with the detection of antifungal and antimicrobial metabolite-synthesizing gene clusters in T7-IITJ (Table 2) indicate the role of T7-IITJ in inhibiting plant pathogens in the desert rhizospheres (Kaushik *et al*. 2023). This can aid in its plant growth promotion properties and allow its use as a biocontrol agent. Other strains of *P. frigoritolerans* showed nematicidal and fungicidal activity due to production of koranimine and other compounds, corroborating our results (Montecillo and Bae, 2022b; Y. Wang et al. 2022). We manually confirmed the absence of T7-IITJ resistance to several antibiotics, viz., kanamycin, ampicillin, gentamycin, and rifampicin, by observing growth up to a week (data not shown). On the other hand, antibiotic-resistance genes in the T7-IITJ genome (Table S9) were not associated with mobile genetic elements (Table S10), which warrants safe use of this strain as a biofertilizer or a biocontrol agent (Chen, Yu and He 2023).

*P. frigoritolerans* strains are known to colonize the internal tissues of plants, thereby promoting stress alleviation by various means (Li *et al*. 2018; Durlik *et al*. 2020; Wang *et al*. 2022b). Generally, PGPRs induce plant genes for enhanced nutrient uptake and stress tolerance, either as biofilms on the root surface or as endophytes (Kasim *et al*. 2016; Calvo *et al*. 2019; Abiala *et al*. 2023). Hence, we decided to study the role of T7-IITJ in regulating global gene expression in *A. thaliana* seedlings. Transcriptome analysis revealed the upregulation of several pathways contributing to the growth promotion of *A. thaliana* under drought stress due to T7-IITJ (Table 4, Table S6). Photosynthesis is vital for plant growth, compromised under drought stress (Zargar *et al*. 2017). Our gene enrichment analysis showed upregulation of 11 photosynthetic pathway genes in *A. thaliana,* which can explain the enhanced growth under PEG upon T7-IITJ inoculation, involving increased photosynthetic functions. These genes were involved in an extensive network of functionally interacting genes (Fig. S3). This correlates with the increased chlorophyll content observed in T7-IITJ-inoculated plants (Fig. 4). Previously, inoculation with *Bacillus* species significantly upregulated chlorophyll-binding proteins in *A. thaliana* (Chang *et al*. 2023) and led to improvement of chlorophyll content and photosynthesis rate in other plants (Jang *et al*. 2018; Abiala *et al*. 2023). Auxin and SA pathways were significantly enriched in our study, which can directly correlate with the root development of *A. thaliana* under drought stress. While our biochemical tests revealed that T7-IITJ could produce auxin (Fig. 2), the transcriptome indicated that the bacterium upregulated many plant genes involved in auxin response (Table 3, Table S6). Phytohormone production by PGPRs is a well-studied area (Li et al. 2023), and previously, *B. subtilis* showed auxin and SA pathway enrichment in *A. thaliana* (Chang *et al*. 2023). The super pathways of geranylgeranyl diphosphate biosynthesis via MEV or MEP were enriched in our study. These pathways supply precursors of the phytohormones: gibberellic acid and abscisic acid (ABA) for growth and drought response, respectively, and isoprenoids related to photosynthesis (Rodríguez-Concepción *et al*. 2004; Morrone *et al*. 2009). Several species of *Azospirillum sp.* produce different forms of gibberellic acid, increasing plant growth (Lucangeli and Bottini 1997; Cassán *et al*. 2011; Castillo *et al*. 2015). The MEV pathway is also the hallmark of plant-symbiotic bacteria signaling (Venkateshwaran *et al*. 2015). Pathways related to ubiquinone and other terpenoid quinone biosynthesis were upregulated in our study. Ubiquinone is an essential component of the electron transport chain, expressed in response to stress. Wheat seedlings inoculated with *Bacillus sp.* showed ubiquinone-enriched pathways (Zhao *et al*. 2022). T7-IITJ enriched the pyridine nucleotide cycling pathway, essential for redox homeostasis and plant growth under stress (Noctor, Queval and Gakière 2006). Enrichment of pathways related to iron ion homeostasis in the upregulated gene set suggests the role of T7-IITJ in facilitating iron acquisition and sequestration in plant cells from iron-deficient soils (Table S2), correlating with our biochemical detection of siderophore production by this bacterium (Fig. S2), and increased accumulation of iron in plant tissues (Fig. 6). Enrichment of iron pathways were previously documented in *A. thaliana* upon inoculation with *Bacillus megaterium* (Liu *et al*. 2023).

While we observed the upregulation of some drought and ABA-inducible genes in our transcriptome (Table 3), we also found the downregulation of many other ABA- and stress-responsive genes of *A. thaliana*, viz. *KIN1*, *dehydrins* like *RAB18*, and several *LEA* paralogs upon inoculation by T7-IITJ (Supplemental Tables 5, 7, 8), which is puzzling since this bacterium elevated drought stress tolerance. Previously, an endophytic bacterium, *Acinetobacter baumannii* MZ30V92, downregulated the drought-responsive genes in *Zea mays* (Sandhya and Ali 2019). Drought stress supports the colonization of microbes in plant roots (Vargas et al. 2014), and in some cases, they activate drought-responsive genes like *ERD15*, *RAB18,* and *DREB*s (Timmusk 2003; Vaishnav and Choudhary 2019) while in other, repress those genes but still alleviate the stress. For example, *Gluconacetobacter diazotrophicus*-inoculated sugarcane and *Rhizobium leguminosarum* LET4910-inoculated wheat showed downregulation of DREBs compared to the non-inoculated plants (Vargas *et al*. 2014; Barquero *et al*. 2022). Inoculation of *Pseudomonas chlororaphis* O6 induced systemic drought tolerance but downregulated drought-responsive genes *ABF3*, *ABI1*, *dehydrins, HNL4*, and *LEA3* in *A. thaliana* (Cho, Kang and Kim 2013). The explanation provided by these researchers for this phenomenon is that the endophytes pre-sensitize or prime the plants by downregulating a subset of ABA and drought-responsive genes and reducing the tissue levels of the drought hormones to result in stress alleviation. Other studies indicated that genes other than the classical ABA and drought-responsive genes could be responsible for increased growth under drought stress due to PGPRs. The study of *A. thaliana* mutants with altered signaling pathways also indicated that ABA was not the primary hormone responsible for drought tolerance. Rather, JA could be (Cho *et al*. 2012; Cho, Kang and Kim 2013). In another study, inoculation of *Enterobacter cloacae* UW4 upregulated the growth-responsive genes but downregulated ethylene-responsive genes in canola roots (Hontzeas, Saleh and Glick 2004). A similar observation was documented upon inoculation of *P. fluorescens* FPT9601-T5, which upregulated the auxin-responsive genes but downregulated the ethylene-responsive genes, essential components of drought tolerance in *A. thaliana* (Wang *et al*. 2005). Hence, T7-IITJ improved drought tolerance in *A. thaliana* through the upregulation of different biochemical pathways involved in JA signaling, auxin response, photosynthesis, enhanced nutrient uptake, secondary metabolite biosynthesis, etc., to alleviate drought stress while downregulating a subset of drought-responsive genes.

It can be concluded from this study that *Peribacillus frigoritolerans* T7-IITJ is a promising PGPR for developing biofertilizers to improve the growth and drought tolerance of plants in arid regions. However, the efficacy of this isolate needs to be tested under field conditions. This local isolate can be a better alternative to biofertilizer consortia imported from outside states that fail to acclimatize under the arid conditions of Rajasthan and, hence, cannot give the desired results. Furthermore, the whole genome sequence of T7-IITJ unraveled in the present study will also pave the way for the future improvement of its biofertilization and biocontrol potential by genetic engineering and genome editing.

## Statements and Declarations

### Significance and Impact of the Study

This study strongly suggests the potential of *P. frigoritolerans* T7-IITJ, a Thar desert rhizobacterium, to develop biofertilizers and biocontrol agents for sustainable agriculture under drought, while unravelling the genome-wide mechanisms of plant growth promotion.

### Ethical approval

This study did not involve any animal or human subjects and doesn’t require ethical approval. This is an original work not published or under review in other journals. The authors declare that no fabrication, plagiarism, or misappropriation was done in the work.

### Data availability

The 16S rDNA sequences of *Enterobacter cloacae* C1P-IITJ, *Kalamiella piersonii* J4-IITJ, and *Peribacillus frigoritolerans* T7-IITJ are available at NCBI GenBank with accession numbers OQ991942, OQ992202, and OQ991918, respectively. The whole genome sequence of T7-IITJ is available at NCBI with an accession number JAUPFL000000000. The raw transcriptome sequences are available at the NCBI SRA database with a BioProject accession number PRJNA1020649.

### Author contributions

DM performed most of the experiments. MA and AS designed and supervised the experiments. PS and NSC carried out bacterial genome sequencing and some biochemical tests. NJ and DV helped in plant phenotyping. MM and NJ helped in rhizosphere soil sampling and analysis. RSS and PY performed PPI network analysis. AS prepared the figures. DM and AS drafted the paper with inputs from all the remaining authors.

## Supplementary Data

**Fig. S1.** PCR amplification and deduplication analysis of 16S rDNA fragments from genomic DNA of rhizobacterial isolates. **A** PCR amplicons of 16S rDNA fragments amplified from the genomic DNA of the isolates C1P, J4, and T7 are shown in a 1% agarose gel. L: 100 bp DNA ladder. **B** A deduplication analysis performed by restriction fragment length polymorphism (RFLP), i.e., digesting the purified PCR fragment with HaeIII, a four-base cutting restriction enzyme. Out of the samples with similar band patterns, denoted by the same number below, only one was used for Sanger sequencing.

**Fig. S2.** Nitrate reduction, siderophore production, and antifungal activity of *Peribacillus frigoritolerans* T7-IITJ. **A** Nitrate reduction test results are shown as the nitrite concentration produced in µg/ml (see Methods). The bar indicates an average of three biological replicates with standard error. The asterisk indicates significant differences between bacterial and control samples (*P* < 0.05, Student’s *t*-test). **B** Siderophore production of the bacterium, on CAS medium, is seen as an orange halo around the bacterial colony due to the chelation of iron from the blue dye in the medium (see Methods). The mean halo diameter of six colonies is shown below, with standard error. **C** The diameters of the zones of growth inhibition (ZOI) of *Rhizoctonia solani* and *Fusarium oxysporum* around disks inoculated with T7-IITJ in LB agar plates spread with the fungal cultures (see Methods) are plotted. Sterile PBS was inoculated on control disks (C). Bars indicate averages of three colonies (1, 2, and 3) with standard error.

**Fig. S3.** Protein-protein interaction network of genes differentially expressed by *Peribacillus frigoritolerans* T7-IITJ. The association between functionally interacting proteins was identified, and enrichment analysis was conducted using the STRING database with all the differentially expressed genes with -1 < log_2_FC > 1 (see Table S5) and visualized using Cytoscape 3.10.1. The node color indicates upregulated genes (in pink) and downregulated genes (in green), and the colored circle around the nodes represents different enriched pathways to which these nodes are linked. **A** The PPI network for upregulated genes. **B** The PPI network for downregulated genes. **C** A key to the node size representing the degree of the nodes, and a color key of the nodes representing the log2FC value of genes, are shown. The color chart represents different enriched pathways along with their annotation identity.

**Fig. S4.** Seedling growth assay of *Tephrosia purpurea* inoculated with *Peribacillus frigoritolerans* T7-IITJ. Plant seeds were collected locally from the IIT Jodhpur campus. Surface sterilized seeds were imbibed in distilled water for 2-3 d and transferred to 250 ml ¼^th^ Hoagland media, pH 5.8, under hydroponic growth conditions. One milliliter of T7-IITJ resuspended in phosphate-buffered saline (PBS) was added as inoculum. Sterile PBS without bacteria was added as a mock. The seedlings were photographed after 10 d. Plants were considered dead if the shoot withered after 14 d. The survival rates and root lengths in 10% PEG are presented in the graphs. Bars indicate an average of three independent experiments, six seeds each. Asterisks indicate significant differences between inoculated and mock experiments (* *P* < 0.06, \*\**P* < 0.05, Student’s *t*-test).

**Fig. S5.** Germination assay of *Triticum aestivum* (43^rd^ ESWYT 103) inoculated with *Peribacillus frigoritolerans* T7-IITJ. Surface sterilized seeds were placed on cotton wool soaked in 20 ml distilled water (control) or 5% PEG-6000 in Petri dishes. One milliliter of T7-IITJ resuspended in phosphate-buffered saline (PBS) was added as inoculum. Sterile PBS without bacteria was added as a mock. The seedlings were photographed after 5 d. Sprouted seeds with visible roots were germinated. The germination rates and root and shoot lengths of germinated seedlings are shown as a graph. Bars indicate an average of three independent experiments, 20 seeds each. Asterisks indicate significant differences between inoculated and mock experiments (*P* < 0.05, Student’s *t*-test).

**Fig. S6.** Germination assay of *Setaria italica* (GP-125) inoculated with *Peribacillus frigoritolerans* T7-IITJ. Surface sterilized seeds were placed on cotton wool soaked in 20 ml distilled water (control) or 5% PEG-6000 in Petri dishes. One milliliter of T7-IITJ resuspended in phosphate-buffered saline (PBS) was added as inoculum. Sterile PBS without bacteria was added as a mock. The seedlings were photographed after 10 d. The germination rates and root and shoot lengths of germinated seedlings are shown as a graph. Bars indicate an average of three independent experiments, 20 seeds each. Asterisks indicate significant differences between inoculated and mock experiments (*P* < 0.05, Student’s *t*-test).

**Table S1.** Oligonucleotides used in the quantitative real-time PCR analysis

**Table S2.** Soil analysis results of sampling sites of Thar desert rhizobacteria

**Table S3.** Nucleotide identity between species closely related to *Peribacillus frigoritolerans* T7-IITJ

**Table S4.** Tetra correlation between species closely related to *Peribacillus frigoritolerans* T7-IITJ

**Table S5.** Genes differentially expressed in *Arabidopsis thaliana* seedlings inoculated with *Peribacillus frigoritolerans* T7-IITJ

**Table S6.** Gene Ontology enrichment of genes induced in *Arabidopsis thaliana* seedlings inoculated with *Peribacillus frigoritolerans* T7-IITJ

**Table S7.** Enrichment of biological pathways within genes repressed in *Arabidopsis thaliana* seedlings inoculated with *Peribacillus frigoritolerans* T7-IITJ

**Table S8.** Gene Ontology enrichment of genes repressed in Arabidopsis thaliana seedlings inoculated with *Peribacillus frigoritolerans* T7-IITJ

**Table S9.** Antibiotic resistance genes of *Peribacillus frigoritolerans* T7-IITJ

**Table S10.** Mobilome in *Peribacillus frigoritolerans* T7-IITJ

