## Supplemental files for "*Peribacillus frigoritolerans* T7-IITJ, a potential biofertilizer, induces plant growth-promoting genes of *Arabidopsis thaliana*"

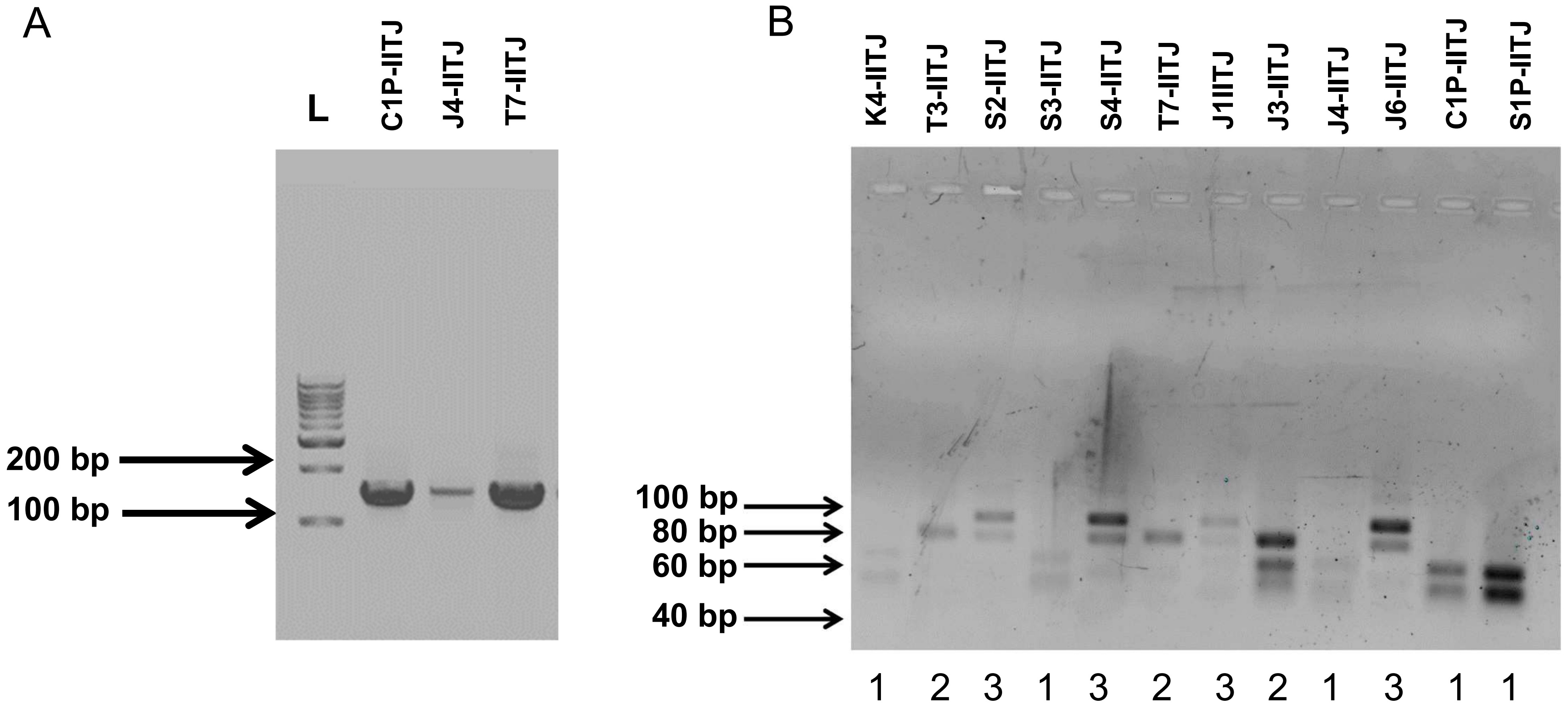


**Supplemental Fig. S1. PCR amplification and deduplication analysis of 16S rDNA fragments from genomic DNA of rhizobacterial isolates. A** PCR amplicons of 16S rDNA fragments amplified from the genomic DNA of the isolates C1P, J4 and T7 are shown in an 1% agarose gel. L: 100 bp DNA ladder. **B** A deduplication analysis performed by restriction fragment length polymorphism (RFLP), i.e., digesting the purified PCR fragment with HaeIII, a four-base cutting restriction enzyme. Out of the samples with similar band patterns, denoted by the same number below, only one was used for Sanger sequencing.

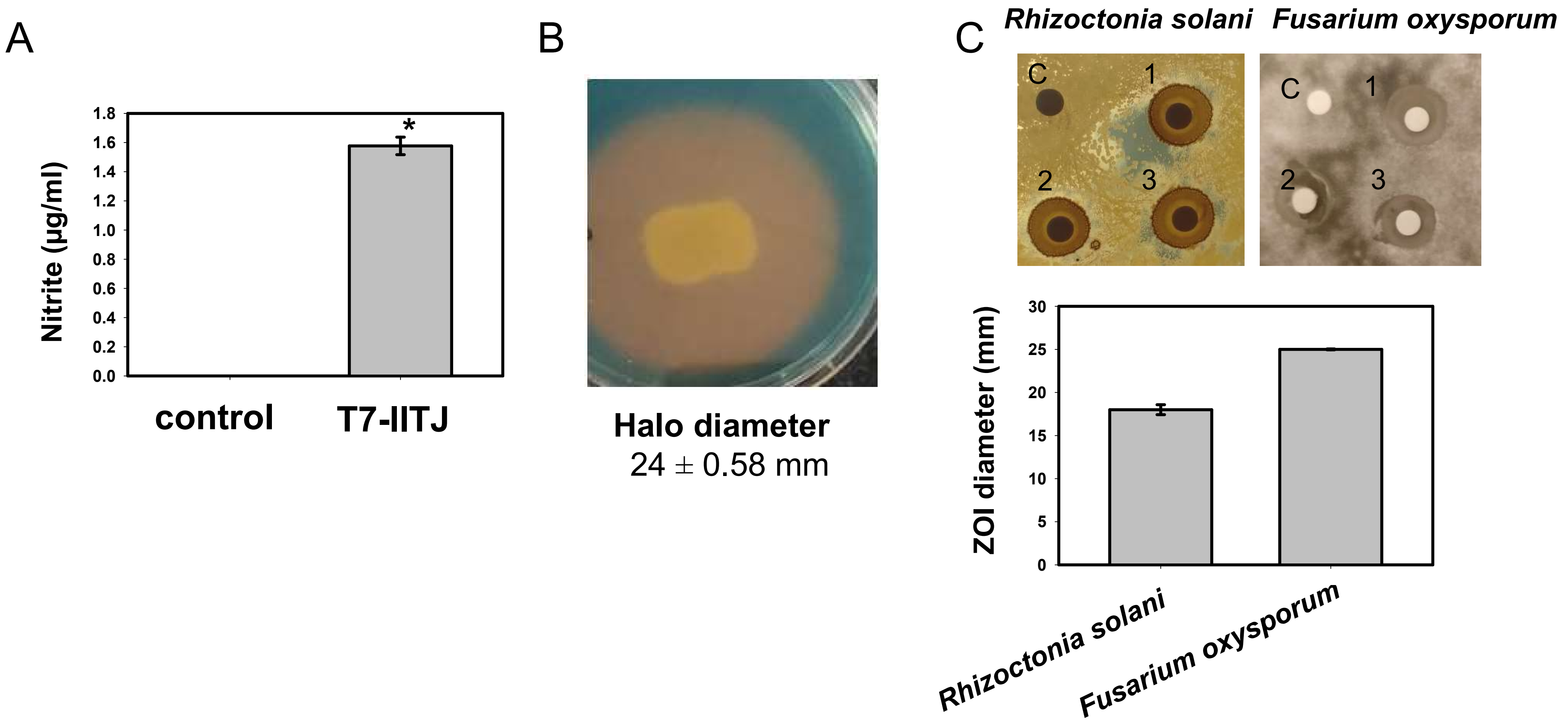


**Supplemental Fig. S2. Nitrate reduction, siderophore production and antifungal activity of *Peribacillus frigoritolerans* T7-IITJ. A** Nitrate reduction test results are shown as the nitrite concentration produced in µg/ml (see Methods). The bar indicates an average of three biological replicates with standard error. The asterisk indicates significant differences between bacterial and control samples ($P < 0.05$, Student's $t$-test). **B** Siderophore production of the bacterium, on CAS medium, is seen as an orange halo around the bacterial colony, due to chelation of iron from the blue dye in the medium (seed Methods). The mean halo diameter of six colonies are shown below, with standard error. **C** The diameters of the zones of growth inhibition (ZOI) of *Rhizoctonia solani* and *Fusarium oxysporum*, around disks inoculated with T7-IITJ in LB agar plates spread with the fungal cultures (see Methods), are plotted. Sterile PBS was inoculated on control disks (C). Bars indicate averages of three colonies (1, 2, and 3) with standard error.

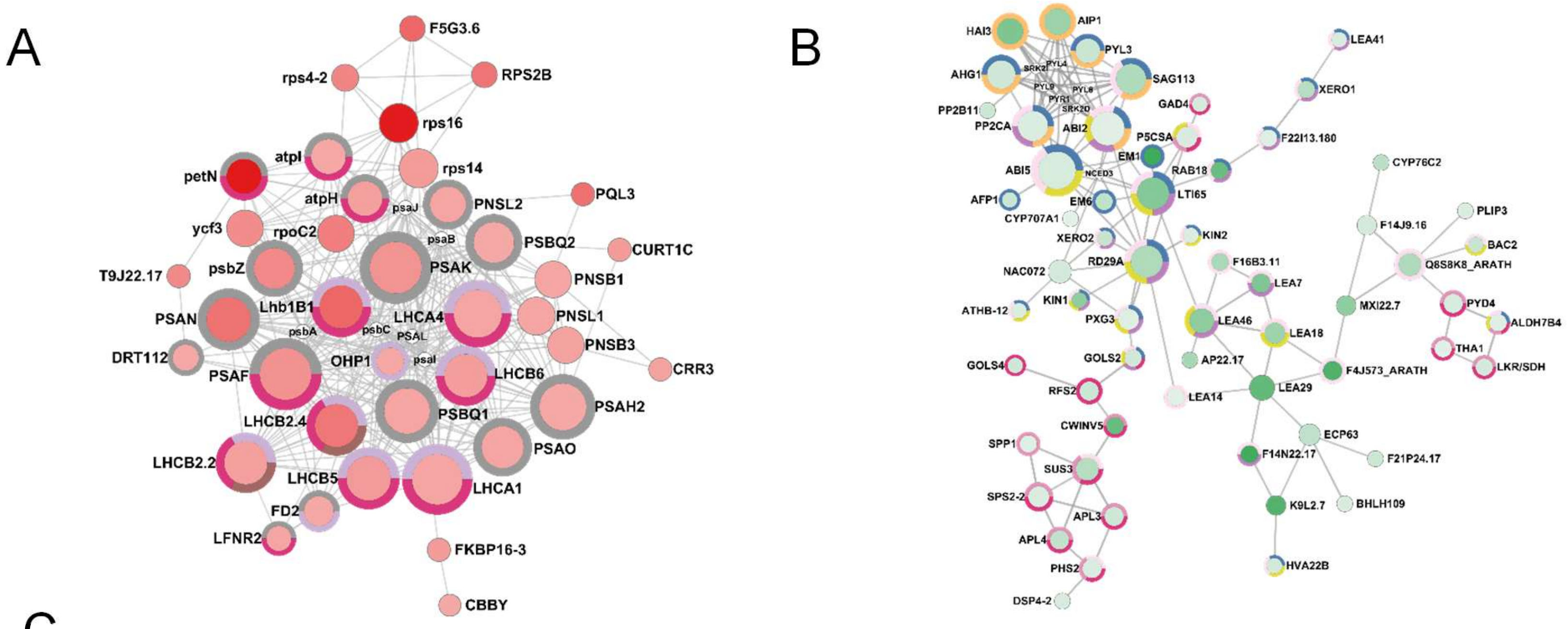


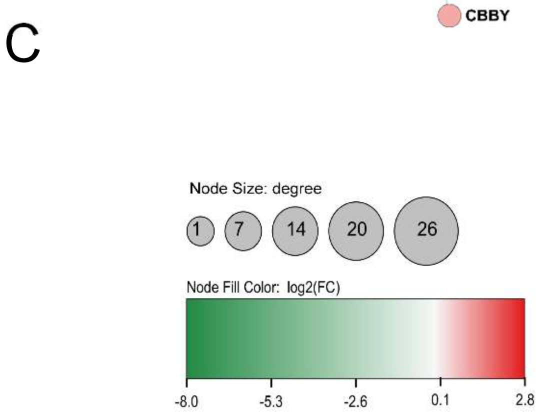


| Category | Color | Annotation ID | Description |
|---|---|---|---|
| GO Biological Process | | GO:0009266 | Response to temperature stimulus |
| GO Biological Process | | GO:0006970 | Response to osmotic stress |
| GO Biological Process | | GO:0009628 | Response to abiotic stimulus |
| GO Biological Process | | GO:0009737 | Response to abscisic acid |
| GO Biological Process | | GO:1903428 | Positive regulation of reactive oxygen species biosynthetic process |
| GO Biological Process | | GO:0009416 | Response to light stimulus |
| KEGG Pathways | | ath01100 | Metabolic pathways |
| KEGG Pathways | | ath01110 | Biosynthesis of secondary metabolites |
| KEGG Pathways | | ath04016 | MAPK signaling pathway - plant |
| KEGG Pathways | | ath00195 | Photosynthesis |

**Supplemental Fig. S3. Protein-protein interaction network of genes differentially expressed by *Peribacillus frigoritolerans* T7-IITJ.** The association between functionally interacting proteins was identified by a predicted protein-protein interaction (PPI) network, and enrichment analysis, conducted using the STRING database with all the differentially expressed genes with -1 < $\log_2$FC > 1 (see Supplemental Table 5) and visualized using Cytoscape 3.10.1. The node color indicates upregulated genes (in pink) and downregulated genes (in green), and the colored circle around the nodes represents different enriched pathways to which these nodes are linked. **A.** The PPI network for upregulated genes. **B.** The PPI network for downregulated genes. **C** A key to the node size representing the degree of the nodes, and a color key of the nodes representing the $\log_2$FC value of genes, are shown. The color chart represents different enriched pathways along with their annotation identity.

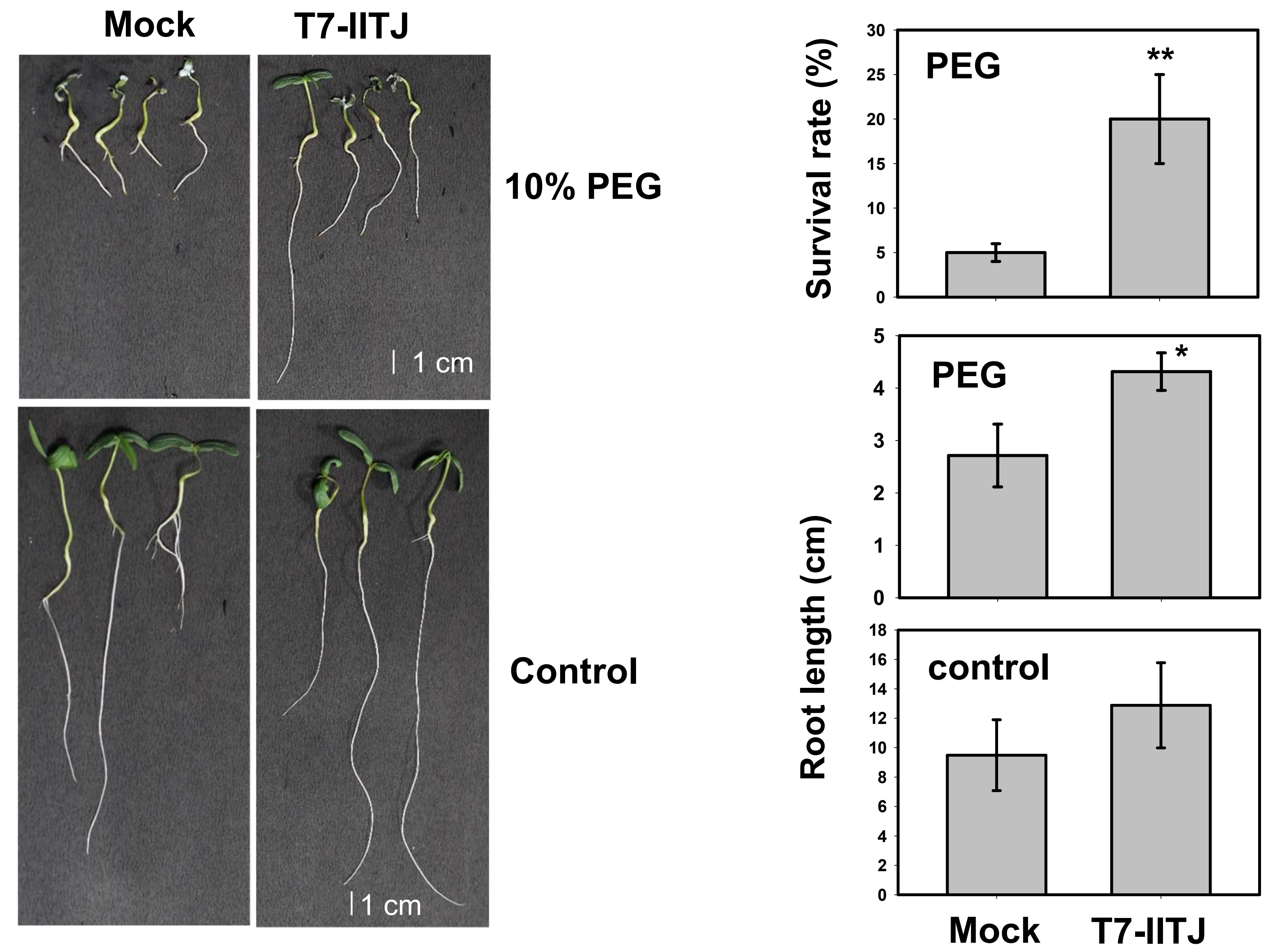


**Supplemental Fig. S4. Seedling growth assay of *Tephrosia purpurea* inoculated with *Peribacillus frigoritolerans* T7-IITJ**. Plant seeds were collected locally from the IIT Jodhpur campus. Surface sterilized seeds were imbibed in distilled water for 2-3 d and transferred to 250 ml ¼$^{th}$ Hoagland media, pH 5.8 under hydroponic growth conditions. One milliliter of T7-IITJ resuspended in phosphate-buffered saline (PBS) was added as inoculum. Sterile PBS without bacteria was added as mock. The seedlings were photographed after 10 d. Plants were considered dead if the shoot withered after 14 d. The survival rates and root lengths in 10% PEG are presented in the graphs. Bars indicate average of three independent experiments, 6 seeds each. Asterisks indicate significant differences between inoculated and mock experiments (* $P < 0.06$, **$P < 0.05$, Student's *t*-test).

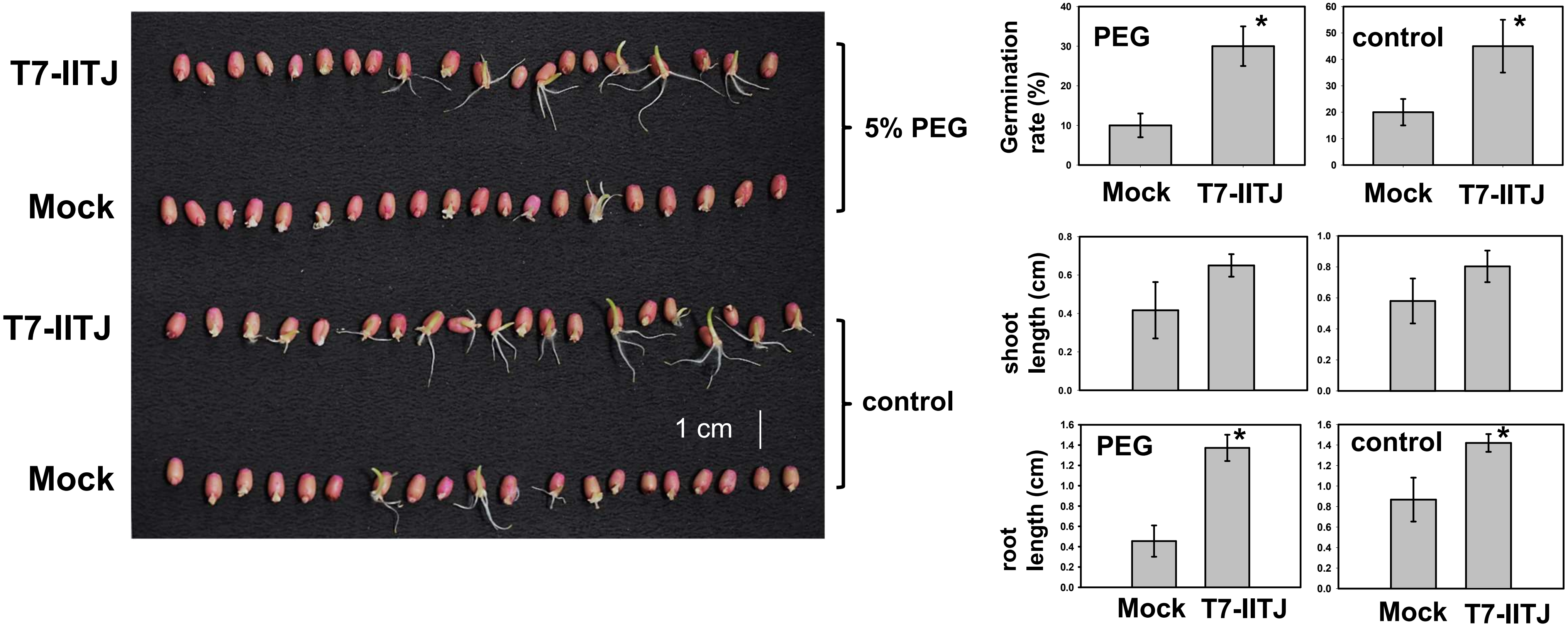


**Supplemental Fig. S5. Germination assay of *Triticum aestivum* (43rd ESWYT 103) inoculated with *Peribacillus frigoritolerans* T7-IITJ.** Surface sterilized seeds were placed on cotton wool soaked in 20 ml distilled water (control) or 5% PEG-6000 in petri dishes. One milliliter of T7-IITJ resuspended in phosphate-buffered saline (PBS) was added as inoculum. Sterile PBS without bacteria was added as mock. The seedlings were photographed after 5 d. Sprouted seeds with visible root were considered to be germinated. The germination rates, as well as root and shoot lengths of germinated seedlings are shown as a graph. Bars indicate average of three independent experiments, 20 seeds each. Asterisks indicate significant differences between inoculated and mock experiments ($P < 0.05$, Student's $t$-test).

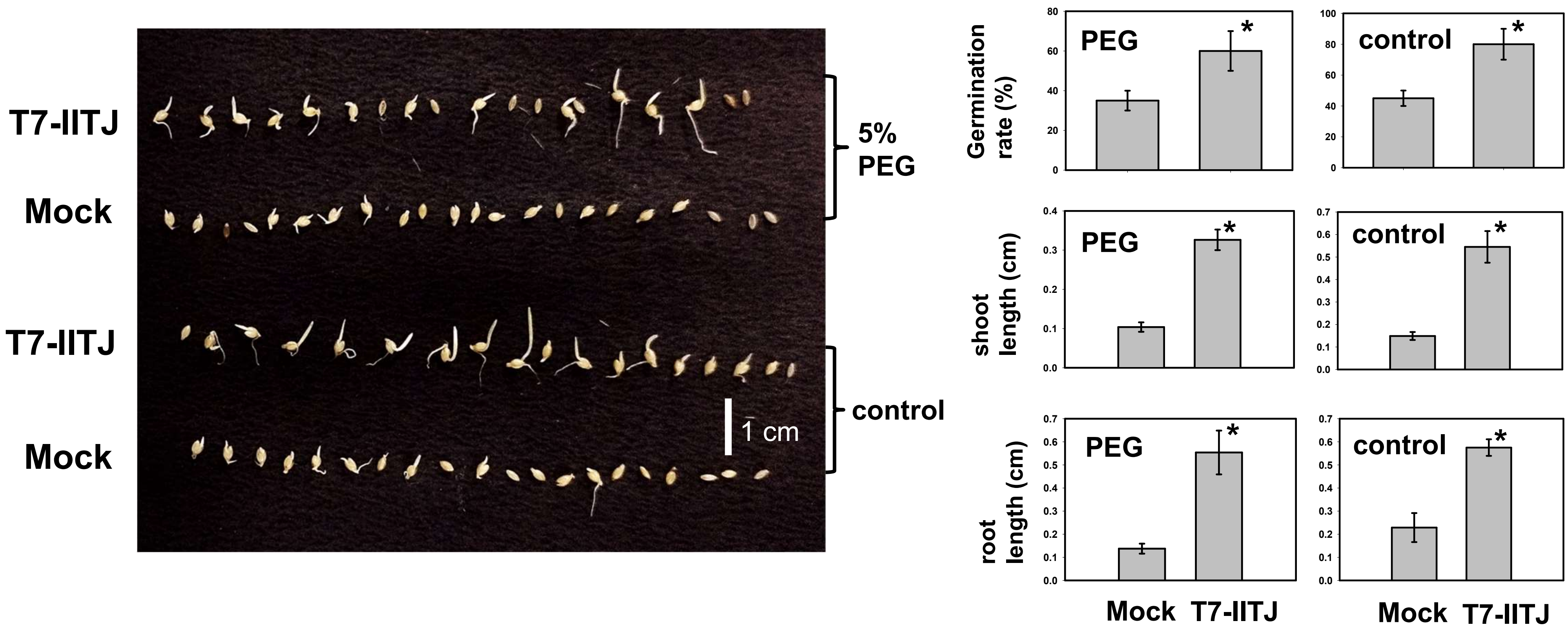


**Supplemental Fig. S6. Germination assay of *Setaria italica* (GP-125) inoculated with *Peribacillus frigoritolerans* T7-IITJ.** Surface sterilized seeds were placed on cotton wool soaked in 20 ml distilled water (control) or 5% PEG-6000 in Petri dishes. One milliliter of T7-IITJ resuspended in phosphate-buffered saline (PBS) was added as inoculum. Sterile PBS without bacteria was added as a mock. The seedlings were photographed after 10 d. The germination rates and root and shoot lengths of germinated seedlings are shown as a graph. Bars indicate an average of three independent experiments, 20 seeds each. Asterisks indicate significant differences between inoculated and mock experiments ($P < 0.05$, Student's $t$-test).

**Supplemental Table S1. Oligonucleotides used in the quantitative real-time PCR analysis**

| Oligonucleotide Name | Sequence (5'->3') |
|---|---|
| AT3G25190RT_Fw | GGTTAAGTCGAGCGTCAGGG |
| AT3G25190RT_Rv | TGGAAAGAAGGACCACATGAAAGA |
| AT5G46890RT_Fw | GATCTTGAAGCCGCGGTCTG |
| AT5G46890RT_Rv | TCTCTAGGCACATTTGAAACCAGAT |
| AT4G19030RT_Fw | ACCGATAAACCATTGCGAGAAATCA |
| AT4G19030RT_Rv | ACTCCATCAACTCCACCCATGT |
| AT1G52070RT_Fw | ACATATGGGTTTGTGTCAGGTACC |
| AT1G52070RT_Rv | CCACCATCCCGCCCATAGAA |
| AT4G12510RT_Fw | GTCTCTGCACCGCCCTAAAG |
| AT4G12510RT_Rv | AGGAACCTTCTTGCCACAAACA |
| AT5G46900RT_Fw | GAAGCCGCGGTCTGTCTTTG |
| AT5G46900RT_Rv | AGGACAACGTTCAAAGAAATGGGA |
| *AtUBQ1* _qPCR_Fw | TCGTAAGTACAATCAGGATAAGATG |
| *AtUBQ1* _qPCR_Rv | CACTGAAACAAGAAAAACAAACCCT |

Oligonucleotide primers used in the study for real time quantitative PCR are shown

**Supplemental Table S2. Soil analysis results of sampling sites of Thar desert rhizobacteria**

| Standard and its unit | pH (Acidity/ Alkalinity) | Electron Conductivity (Total dissolved salts; dS/m) | Organic Carbon (%) | Available Phosphorous ($P_2O_5$; Kg/ha) | Nitrogen (Kg/ha) | Available Potash ( $K_2O$; Kg/ha) | Available Zinc (ppm) | Available Iron (ppm) | Available Copper (ppm) | Available Manganese (ppm) | Available Molybdenum (ppm) | Available Boron (ppm) |
|---|---|---|---|---|---|---|---|---|---|---|---|---|
| **Standard values** | Normal (7.0-8.5), Basic (8.5-9.5), Highly Basic (> 9.5) | Normal (till 1.5), Saline (1.5-3.0), Highly saline (> 3.0) | Low (< 0.5), Medium (0.5-0.75), High (> 0.75) | Low (< 23.0), Medium (23.0-56.0), High (> 56.0) | Low (< 240) Medium (240-480) High (> 480) | Low (< 144), Medium (142-336), High (> 336) | Low (< 0.6), Normal (> 0.6) | Low (1.0-3.0) Normal (4.0-6.0) High (> 6.0) | Low (< 0.2), Normal (> 0.2) | Low (< 0.2), Normal (> 0.2) | Low (0-0.1) Normal (0.1-0.5) High (> 0.5) | Low (0.1-1.0) Normal (1-2) High (> 2.0) |
| **Desert soil** | 8.6 | 0.18 (Normal) | 0.21 (Low) | 36 (Medium) | 150-200 (Low) | 212 (Medium) | 0.38 (Low) | 3.42 (Low) | 0.52 (Normal) | 5.26 (Normal) | Not detected | Not detected |
| **Rhizosoil 1 (*Tephrosia purpurea*)** | 8.7 | 0.18 (Normal) | 0.17 (Low) | 44 (Medium) | 150-200 (Low) | 240 (Medium) | 0.32 (Low) | 3.18 (Low) | 0.42 (Normal) | 5.2 (Normal) | Not detected | Not detected |
| **Rhizosoil 2 (*Prosopis cineraria*)** | 8.5 | 0.22 (Normal) | 0.13 (Low) | 36 (Medium) | 150-200 (Low) | 339 (High) | 0.34 (Low) | 3.26 (Low) | 0.46 (Normal) | 5.16 (Normal) | Not detected | Not detected |
| **Desert soil + soil-rite mix (3:1)** | 8.6 | 0.25 (Normal) | 0.19 (Low) | 36 (Medium) | 150-200 (Low) | 226 (Medium) | 0.36 (Low) | 3.34 (Low) | 0.48 (Normal) | 5.2 (Normal) | Not detected | Not detected |

The results of soil analysis are shown. This includes pH, electron conductivity/ soil salinity, % organic carbon, and nutrients P, K, Zn, Cu, and Mn, estimated by The Soil Admin Lab, Jodhpur. Rhizosoils were collected around the roots of *Prosopis cineraria* and *Tephrosia purpurea*. Desert soil indicates soil collected from the same site, where there were no plants, at approximately the same depth as the rhizosoils (10-15 cm from the top). The composition of a mixture of desert soil and soil-rite (3:1), used in the plant growth promotion asay (Fig. 2), is also shown. Soil rite consists of perlite, peat moss and vermiculite 1:1:1. Mean values of three technical replicates are presented. The standard values of pH and nutrients are also shown for comparison.

**Supplemental Table S3. Nucleotide identity between species closely related to *Peribacillus frigoritolerans* T7-IITJ**

| | ***Peribacillus frigoritolerans T7-IITJ*** | ***Peribacillus* AS _1** | ***Peribacillus* BBB004** | ***Peribacillus faecalis* AGMB02131** | ***Peribacillus frigoritolerans* CF13** | ***Peribacillus frigoritolerans* FD2** | ***Peribacillus frigoritolerans* HMB20428** | ***Peribacillus frigoritolerans* LN4** | ***Peribacillus frigoritolerans* MER73** | ***Peribacillus* sp.SI84** | ***Peribacillus simplex MSL17322*** |
|---|---|---|---|---|---|---|---|---|---|---|---|
| ***Peribacillus frigoritolerans T7-IITJ*** | * | 94.35 | 92.56 | 68.85 | **99.29** | 68.2 | 94.78 | 94.72 | 94.63 | 82.76 | 93.36 |
| ***Peribacillus* AS_1** | 93.92 | * | 91.72 | 67.7 | **96.21** | 67.49 | **96.28** | **96.25** | **96.22** | 82.07 | 92.5 |
| ***Peribacillus* BBB004** | 91.98 | 91.63 | * | 68.06 | 91.79 | 67.63 | 91.85 | 91.83 | 91.89 | 82.02 | 92.54 |
| ***Peribacillus faecalis* AGMB02131** | 68.04 | 67.97 | 68.14 | * | 68.01 | 67.39 | 68.03 | 68.07 | 68.07 | 67.87 | 68.17 |
| ***Peribacillus frigoritolerans* CF13** | **99.29** | **96.3** | 92.05 | 68.64 | * | 68.06 | **99.24** | **98** | **96.74** | 82.38 | 92.91 |
| ***Peribacillus frigoritolerans* FD2** | 67.85 | 67.87 | 67.84 | 67.53 | 67.94 | * | 67.88 | 67.89 | 68.17 | 67.54 | 67.85 |
| ***Peribacillus frigoritolerans* HMB20428** | 94.56 | **96.39** | 92.2 | 68.73 | **94.51** | 68.32 | * | **98.1** | **96.84** | 82.6 | 93.09 |
| ***Peribacillus frigoritolerans* LN4** | 94.62 | **96.51** | 92.13 | 68.6 | **98.11** | 68.3 | **98.18** | * | **96.82** | 82.54 | 93.05 |
| ***Peribacillus frigoritolerans* MER73** | 94.24 | **96.33** | 91.95 | 68.09 | **96.72** | 67.89 | **96.76** | **96.57** | * | 82.08 | 92.67 |
| ***Peribacillus* SI84** | 82.71 | 82.53 | 82.48 | 67.67 | 82.62 | 67.25 | 82.62 | 82.59 | 82.54 | * | 82.91 |
| ***Peribacillus simplex* MSL17322** | 92.97 | 92.58 | 92.73 | 68.26 | 92.72 | 67.81 | 92.8 | 92.78 | 92.69 | 82.5 | * |

Average nucleotide identity between species closely related to *Peribacillus frigoritolerans* T7-IITJ, calculated using Prokka 1.12. J-species software (http://jspecies.ribohost.com/jspeciesws/).

**Supplementary Table 4. Tetra correlation between species closely related to *Peribacillus frigoritolerans* T7-IITJ**

| Microbes | Z-Score |
|---|---|
| *Peribacillus frigoritolerans* FJAT-2396 | 0.99852 |
| *Bacillus* sp. FJAT-22058 | 0.99837 |
| *Bacillus* sp. FJAT-21352 | 0.99831 |
| *Peribacillus frigoritolerans* DSM 8801 | 0.99821 |
| *Peribacillus simplex* P558 | 0.99802 |
| *Bacillus* sp. Soil745 | 0.99778 |
| *Peribacillus simplex* NBRC 15720 = DSM 1321 NBRC 15720 | 0.99701 |
| *Peribacillus simplex* NBRC 15720 = DSM 1321 | 0.99684 |
| *Peribacillus simplex* SH-B26 | 0.99459 |
| *Peribacillus muralis* DSM 16288 | 0.98306 |
| *Peribacillus muralis* G25-68 | 0.98279 |
| *Peribacillus simplex* VanAntwerpen02 | 0.98028 |
| *Priestia abyssalis* DSM 25875 | 0.95442 |
| *Bacillus* sp. SG-1 | 0.95239 |
| *Bacillus* sp. Soil768D1 | 0.94772 |
| *Peribacillus butanolivorans* DSM 18926 | 0.94698 |
| *Bacillus sp* . Leaf13 | 0.946 |
| *Peribacillus glennii* V44-8 | 0.94501 |
| *Siminovitchia fordii* DSM 16014 = CIP 108821 DSM 16014 | 0.94383 |
| *Peribacillus cavernae* L5 | 0.94364 |
| *Sporosarcina globispora* DSM 4 | 0.93946 |
| *Peribacillus saganii* V47-23a | 0.93924 |
| *Cytobacillus oceanisediminis* CGMCC 1.10115 | 0.93881 |
| *Bacillus salipaludis* WN066 | 0.93757 |
| *Falsibacillus albus* GY 10110 | 0.93572 |
| *Siminovitchia fortis* DSM 16012 | 0.93303 |
| *Mesobacillus harenae* Y40 | 0.93108 |
| *Cytobacillus oceanisediminis 2691* | 0.93052 |
| *Bacillus salacetis* SKP7-4 | 0.92965 |
| *Neobacillus endophyticus* BRMEA1 | 0.929 |
| *Siminovitchia acidinfaciens* 3-2-2 | 0.92863 |
| *Bacillus methanolicus* MGA3 | 0.92755 |
| *Bacillus methanolicus* MGA3 | 0.92711 |
| *Mesobacillus zeae* JJ-247 | 0.92691 |
| *Neobacillus bataviensis* LMG 21833 | 0.9268 |
| *Mesobacillus zeae* JJ-247 | 0.92669 |
| *Bacillus acidicola* FJAT-2406 | 0.92613 |

| | |
|---|---|
| *Neobacillus drentensis* NBRC 102427 | 0.92552 |
| *Neobacillus citreus* FJAT-50051 | 0.92539 |
| *Bacillus* sp. 2_A_57_CT2 | 0.92522 |
| *Bacillus oleivorans* JC228 | 0.92519 |
| *Mesobacillus foraminis* CV53 | 0.92499 |
| *Mesobacillus foraminis* CV53 | 0.92483 |
| *Neobacillus rhizophilus* FJAT-49825 | 0.92469 |
| *Bacillus methanolicus* PB1 | 0.92392 |
| *Neobacillus drentensis* FJAT-10044 | 0.92389 |
| *Bacillus renqingensis* REN2 | 0.92378 |
| *Neobacillus* soli NBRC 102451 | 0.92365 |
| *Cytobacillus horneckiae* 1P01SC | 0.92362 |
| *Bacillus massiliglaciei* Marseille-P2600 | 0.92355 |
| *Neobacillus vireti* DSM 15602 | 0.92347 |
| *Neobacillus vireti* LMG 21834 | 0.92341 |
| *Neobacillus* soli DSM 15604 | 0.9233 |
| *Cytobacillus eiseniae* FJAT-2352 | 0.92286 |
| *Metabacillus dongyingensis* BY2G20 | 0.92248 |
| *Neobacillus kokaensis* LOB 377 | 0.92227 |
| *Neobacillus fumarioli NBRC* 102428 | 0.92152 |
| *Neobacillus massiliamazoniensis* LF1 | 0.92148 |
| *Bacillus norwichensis* Sa1BUA2 | 0.92131 |
| *Neobacillus novalis* NBRC 102450 | 0.92122 |
| *Bacillus benzoevorans* DSM 5391 | 0.92112 |
| *Cytobacillus firmus* NCTC10335 | 0.92083 |
| *Cytobacillus praedii* FJAT-25547 | 0.92076 |
| *Neobacillus novalis* FJAT-14227 | 0.92037 |
| *Neobacillus* terrae C11 | 0.92022 |
| *Neobacillus muris* DSM 110989 | 0.92022 |
| *Neobacillus cucumis* DSM 101566 | 0.9194 |
| *Bacillus* sp. FJAT-21945 | 0.91903 |
| *Cytobacillus gottheilii* FJAT-2394 | 0.91875 |
| *Cytobacillus firmus* DS1 | 0.91873 |
| *Neobacillus mesonae* FJAT-13985 | 0.91813 |
| *Bacillus mediterraneensis* Marseille-P2366 | 0.91797 |
| *Cytobacillus solani* FJAT-18043 | 0.91779 |
| *Peribacillus alkalitolerans* KCTC 33631 | 0.91772 |
| *Cytobacillus firmus* NBRC 15306 | 0.91743 |
| *Siminovitchia terrae* LMG 29736 | 0.91688 |
| *Bacillus pakistanensis* DSM 24834 | 0.91638 |

| | |
|---|---|
| *Bacillus mesophilum* IITR-54 | 0.91631 |
| *Neobacillus niacini* NBRC 15566 | 0.91605 |
| *Metabacillus arenae* IB182487 | 0.91604 |
| *Neobacillus dielmonensis* null | 0.91597 |
| *Metabacillus idriensis* DSM 19097 | 0.91595 |
| *Metabacillus idriensis* DSM-19097 | 0.91588 |
| Bacillus sp. SA1-12 | 0.91558 |
| *Neobacillus niacini* DSM 2923 | 0.91508 |
| *Bacillus massiliigabonensis* Marseille-P2639 | 0.915 |
| *Oceanobacillus massiliensis* str. N'diop Ndiop | 0.91469 |
| *Bacillus testis* SIT10 | 0.91459 |
| *Peribacillus psychrosaccharolyticus* ATCC 23296 | 0.91337 |
| *Cytobacillus firmus* LK28 | 0.91278 |
| *Aeribacillus pallidus* 8 | 0.91268 |
| *Falsibacillus pallidus* DSM 25281 | 0.91266 |
| *Lysinibacillus yapensis* YLB-03 | 0.91262 |
| *Aeribacillus pallidus* 8m3 | 0.91228 |
| *Bacillus sp.* FJAT-27231 | 0.91205 |
| *Rossellomorea arthrocnemi* EAR8 | 0.91199 |
| *Neobacillus rhizosphaerae* CIP 111895 | 0.91178 |
| *Peribacillus loiseleuriae* FJAT-27997 | 0.9116 |
| *Cytobacillus depressus* BZ1 | 0.91158 |
| *Aeribacillus composti* KCTC 33824 | 0.91102 |

Tetra correlation among *Peribacillus frigoritolerans* with other *Peribacillus* species highlighted by a wide distribution of z-scores

**Supplemental Table S5. Genes differentially expressed in *Arabidopsis thaliana* seedlings inoculated with *Peribacillus frigoritolerans* T7-IITJ**

| AGI Code | $log_2(FC)$ | *P* -value | FDR | Gene Symbol | Araport 11 Short Description |
|---|---|---|---|---|---|
| AT4G06477 | 6.78 | 4.44E-06 | 9.30E-05 | | transposable_element_gene |
| AT5G46900 | 5.42 | 5.99E-26 | 2.29E-23 | | Bifunctional inhibitor/lipid-transfer protein/seed storage 2S albumin superfamily protein |
| AT1G14960 | 5.27 | 4.93E-06 | 1.02E-04 | | Polyketide cyclase/dehydrase and lipid transport superfamily protein |
| ArthCt096 | 5.14 | 1.47E-04 | 1.86E-03 | trnY | tRNA-Tyr |
| AT1G21140 | 4.63 | 1.05E-16 | 1.59E-14 | VTL1 | Vacuolar iron transporter homolog 1 |
| AT4G12510 | 4.49 | 1.37E-19 | 2.89E-17 | AZI5 | Encodes a member of the AZI family of lipid transfer proteins. |
| AT5G46890 | 4.40 | 9.86E-52 | 2.40E-48 | MQD22.2 | Bifunctional inhibitor/lipid-transfer protein/seed storage 2S albumin superfamily protein |
| AT2G33790 | 3.91 | 3.49E-09 | 1.56E-07 | ATAGP30 | Pollen Ole e 1 allergen protein containing 14.6% proline residues |
| AT5G44417 | 3.55 | 1.03E-15 | 1.38E-13 | | Pseudogene of FAD-binding Berberine family protein |
| ArthCt097 | 3.46 | 2.74E-06 | 6.10E-05 | trnE | tRNA-Glu |
| AT3G25190 | 3.30 | 6.79E-61 | 4.40E-57 | VTL5 | Vacuolar iron transporter homolog 2.1 |
| ArthCt099 | 3.23 | 1.52E-05 | 2.70E-04 | trnfM | tRNA-Met |
| AT5G57530 | 3.09 | 4.67E-04 | 4.85E-03 | XTH12 | xyloglucan endotransglucosylase/hydrolase 12 |
| AT2G04480 | 2.96 | 1.91E-07 | 5.81E-06 | T1O3.11 | Uncharacterized protein At2g04480 |
| AT1G52070 | 2.93 | 2.22E-23 | 7.21E-21 | F5F19.13 | Mannose-binding lectin superfamily protein |
| AT5G26130 | 2.91 | 1.63E-03 | 1.33E-02 | ATCAPE8 | CAP (Cysteine-rich secretory proteins, Antigen 5, and Pathogenesis-related 1 protein) superfamily protein |
| AT4G25220 | 2.87 | 8.39E-04 | 7.85E-03 | G3PP2 | Encodes a member of the phosphate starvation-induced glycerol-3-phosphate permease gene family |
| AT2G43870 | 2.86 | 1.76E-07 | 5.39E-06 | F18O19.2 | Pectin lyase-like superfamily protein |
| ArthCp015 | 2.85 | 7.00E-05 | 9.96E-04 | petN | cytochrome b6/f complex subunit VIII |
| ArthCp004 | 2.81 | 1.91E-06 | 4.48E-05 | rps16 | ribosomal protein S16 |
| AT5G20710 | 2.78 | 3.42E-06 | 7.45E-05 | BGAL7 | beta-galactosidase 7 |
| AT3G55515 | 2.71 | 8.92E-06 | 1.71E-04 | DVL8 | Rotundifolia like 7 |
| AT3G50640 | 2.71 | 4.11E-14 | 4.39E-12 | | (thale cress) hypothetical protein |
| AT4G08400 | 2.66 | 4.07E-07 | 1.15E-05 | EXT7 | Extensin-like protein (Proline-rich extensin-like family protein) |
| AT5G28145 | 2.62 | 2.65E-04 | 3.06E-03 | | transposable_element_gene |
| AT4G13390 | 2.59 | 1.50E-05 | 2.69E-04 | EXT12 | Extensin-like protein (Proline-rich extensin-like family protein) |
| AT4G12520 | 2.59 | 9.20E-18 | 1.58E-15 | T1P17.110 | Bifunctional inhibitor/lipid-transfer protein/seed storage 2S albumin superfamily protein |
| AT2G25240 | 2.57 | 3.77E-08 | 1.36E-06 | SERPIN5 | Serine protease inhibitor (SERPIN) family protein |
| AT1G74930 | 2.57 | 2.26E-04 | 2.67E-03 | ERF018 | encodes a member of the DREB subfamily A-5 of ERF/AP2 transcription factor family |
| AT4G19030 | 2.56 | 8.69E-32 | 5.12E-29 | NLM1 | an aquaporin whose expression level is reduced by ABA, NaCl, dark, and desiccation |
| AT5G35480 | 2.52 | 1.62E-11 | 1.17E-09 | MOK9.7 | Uncharacterized protein |
| AT4G15290 | 2.52 | 2.69E-07 | 7.95E-06 | CSLB5 | Cellulose synthase-like protein B5 (AtCslB5) (EC 2.4.1.-) |
| AT5G10990 | 2.50 | 1.31E-05 | 2.40E-04 | SAUR69 | SAUR-like auxin-responsive protein family |
| AT2G04460 | 2.49 | 3.00E-03 | 2.14E-02 | T1O3.13 | transposable_element_gene |
| AT2G32270 | 2.46 | 5.84E-12 | 4.40E-10 | ZIP3 | Zinc transporter 3 (ZRT/IRT-like protein 3) |
| AT1G51470 | 2.40 | 2.88E-15 | 3.64E-13 | TGG5 | Encodes a myrosinase. |
| AT3G29250 | 2.39 | 2.08E-14 | 2.32E-12 | SDR4 | Short-chain dehydrogenase reductase 4 (AtSDR4) (EC 1.1.1.-) |
| AT3G44260 | 2.34 | 3.68E-05 | 5.79E-04 | CAF1A | CCR4-associated factor 1A (EC 3.1.13.4) |
| AT1G13420 | 2.30 | 1.46E-12 | 1.21E-10 | ATST4B | Arabidopsis thaliana sulfotransferase 4B |
| AT2G25160 | 2.29 | 1.10E-10 | 6.65E-09 | CYP82F1 | Cytochrome P450, family 82, subfamily F, polypeptide 1 |
| AT2G21650 | 2.26 | 4.01E-04 | 4.27E-03 | MEE3 | Maternal effect embryo arrest 3 |
| AT4G34810 | 2.25 | 1.51E-04 | 1.90E-03 | SAUR5 | SAUR-like auxin-responsive protein family |
| AT1G30760 | 2.23 | 5.46E-06 | 1.12E-04 | ATBBE-LIKE 13 | Berberine bridge enzyme-like 13 |
| AT3G14510 | 2.23 | 1.66E-05 | 2.92E-04 | | Isoprenyl diphosphate synthase5 |
| AT1G36060 | 2.22 | 5.33E-11 | 3.48E-09 | ERF55 | Encodes a member of the DREB subfamily A-6 of ERF/AP2 transcription factor family |
| AT3G63350 | 2.19 | 8.05E-04 | 7.60E-03 | HSFA7B | Heat stress transcription factor A-7b (AtHsfA7b) (AtHsf-10) |
| AT1G74500 | 2.17 | 8.80E-22 | 2.28E-19 | PRE3 | Encodes a basic helix/loop/helix transcription factor that acts downstream of MP in root initiation |
| AT1G74490 | 2.16 | 8.51E-06 | 1.64E-04 | PBL29 | Protein kinase superfamily protein |
| AT5G35490 | 2.16 | 2.33E-05 | 3.90E-04 | MRU1 | Encodes MRU1 (mto 1 responding up) |
| AT2G36885 | 2.14 | 1.49E-12 | 1.22E-10 | At2g36885 | Translation initiation factor |
| AT4G22666 | 2.13 | 1.68E-07 | 5.18E-06 | LTPG28 | Bifunctional inhibitor/lipid-transfer protein/seed storage 2S albumin superfamily protein |
| AT5G51520 | 2.12 | 3.35E-04 | 3.71E-03 | At5g51520 | (thale cress) hypothetical protein |
| AT2G44110 | 2.11 | 4.61E-06 | 9.64E-05 | MLO | MLO-like protein |
| AT4G35640 | 2.11 | 4.60E-10 | 2.51E-08 | SAT4 | Serine acetyltransferase 4 (AtSAT-4) (AtSERAT3;2) (EC 2.3.1.30) |
| AT5G55570 | 2.10 | 1.64E-06 | 3.91E-05 | MDF20.1 | Transmembrane protein |
| AT3G55970 | 2.07 | 6.25E-05 | 9.05E-04 | JRG21 | Jasmonate-regulated gene 21 |
| AT5G21150 | 2.06 | 1.46E-03 | 1.22E-02 | AGO9 | Protein argonaute 9 |
| AT5G39860 | 2.05 | 8.05E-07 | 2.12E-05 | PRE1 | Transcription factor PRE1 (Basic helix-loop-helix protein 136) |
| AT1G19900 | 2.05 | 9.51E-04 | 8.69E-03 | F6F9.4 | F6F9.4 protein |
| AT5G36910 | 2.04 | 7.24E-14 | 7.41E-12 | THI2.2 | Thionin-2.2 [Cleaved into: Thionin-2.2; Acidic protein] |
| AT2G32550 | 2.02 | 2.14E-06 | 4.94E-05 | NOT9C | Cell differentiation, Rcd1-like protein |
| AT1G59940 | 2.00 | 7.12E-04 | 6.85E-03 | ARR3 | Response regulator 3 |
| AT4G33720 | 1.96 | 5.70E-09 | 2.48E-07 | At4g33720 | (thale cress) hypothetical protein |
| AT1G68650 | 1.95 | 3.00E-06 | 6.61E-05 | At1g68650 | GDT1 family protein |
| AT1G07175 | 1.95 | 4.46E-03 | 2.91E-02 | At1g07175 | Secreted protein |
| AT5G54370 | 1.93 | 9.88E-13 | 8.32E-11 | MDK4.20 | At5g54370 (Late embryogenesis abundant (LEA) protein-like protein) (Root cap protein 2-like protein) |
| AT4G12500 | 1.91 | 3.97E-05 | 6.18E-04 | At4g12500 | pEARLI1-like lipid transfer protein 3 |
| AT5G01330 | 1.90 | 3.64E-06 | 7.86E-05 | PDC3 | Pyruvate decarboxylase 3 (AtPDC3) (EC 4.1.1.1) |
| AT5G35777 | 1.88 | 1.63E-06 | 3.91E-05 | | |
| AT4G28680 | 1.87 | 2.95E-13 | 2.76E-11 | TYRDC | L-tyrosine decarboxylase |
| AT5G17170 | 1.87 | 3.45E-36 | 2.80E-33 | ENH1 | Rubredoxin family protein (Uncharacterized protein At5g17170) |
| AT1G15460 | 1.87 | 1.39E-04 | 1.77E-03 | BOR4 | HCO3-transporter family |
| AT4G36570 | 1.86 | 5.00E-08 | 1.77E-06 | RL3 | Protein RADIALIS-like 3 (AtRL3) (Protein RAD-like 3) |
| AT2G18328 | 1.86 | 1.93E-14 | 2.18E-12 | RL4 | Protein RADIALIS-like 4 (AtRL4) (Protein RAD-like 4) |
| AT5G36180 | 1.86 | 4.48E-05 | 6.85E-04 | SCPL1 | Serine carboxypeptidase-like 1 |
| AT2G34000 | 1.85 | 5.66E-05 | 8.32E-04 | At2g34000 | (thale cress) hypothetical protein |
| AT4G22110 | 1.85 | 1.69E-05 | 2.96E-04 | At4g22110 | GroES-like zinc-binding dehydrogenase family protein |

| | | | | | |
|---|---|---|---|---|---|
| AT4G12360 | 1.85 | 8.79E-04 | 8.14E-03 | LTPG24 | Non-specific lipid transfer protein GPI-anchored 24 (AtLTPG-24) (Protein LTP-GPI-ANCHORED 24) |
| AT1G72645 | 1.85 | 1.50E-08 | 5.96E-07 | At1g72645 | Transmembrane protein |
| AT5G03130 | 1.84 | 6.85E-04 | 6.67E-03 | At5g03130 | Uncharacterized protein |
| AT2G05160 | 1.84 | 2.15E-10 | 1.23E-08 | At2g05160 | CCCH-type zinc fingerfamily protein with RNA-binding domain-containing protein |
| AT5G51720 | 1.84 | 2.16E-03 | 1.67E-02 | NEET | CDGSH iron-sulfur domain-containing protein NEET (At-NEET) |
| AT2G34430 | 1.84 | 3.05E-33 | 2.05E-30 | LHB1B1 | Chlorophyll a-b binding protein, chloroplastic |
| AT4G37610 | 1.83 | 1.04E-07 | 3.40E-06 | BT5 | BTB/POZ and TAZ domain-containing protein 5 (BTB and TAZ domain protein 5) |
| AT4G25470 | 1.83 | 1.71E-04 | 2.12E-03 | DREB1C | Dehydration-responsive element-binding protein 1C (Protein DREB1C) |
| AT3G29110 | 1.82 | 4.35E-04 | 4.57E-03 | At3g29110 | Uncharacterized protein |
| AT1G47600 | 1.82 | 4.34E-10 | 2.38E-08 | BGLU34 | Beta glucosidase 34 |
| AT2G40610 | 1.81 | 3.66E-16 | 5.20E-14 | EXPA8 | Expansin-A8 (AtEXPA8) (Alpha-expansin-8) (At-EXP8) (AtEx8) (Ath-ExpAlpha-1.11) |
| AT5G65350 | 1.79 | 6.48E-04 | 6.39E-03 | At5g65350 | Histone H3-like 5 |
| AT3G45530 | 1.77 | 1.92E-04 | 2.33E-03 | F9K21.110 | Cysteine/Histidine-rich C1 domain family protein (Uncharacterized protein F9K21.110) |
| AT4G01430 | 1.77 | 2.33E-05 | 3.90E-04 | UMAMIT29 | WAT1-related protein |
| AT5G14020 | 1.77 | 9.90E-07 | 2.54E-05 | MAC12.31 | Endosomal targeting BRO1-like domain-containing protein (Uncharacterized protein At5g14020) |
| AT1G52060 | 1.76 | 1.05E-06 | 2.68E-05 | At1g52060 | Jacalin-type lectin domain-containing protein |
| AT1G20120 | 1.76 | 4.03E-07 | 1.14E-05 | At1g20120 | GDSL esterase/lipase At1g20120 (EC 3.1.1.-) (Extracellular lipase At1g20120) |
| AT3G19430 | 1.75 | 1.24E-03 | 1.08E-02 | At3g19430 | Late embryogenesis abundant (LEA) hydroxyproline-rich glycoprotein family |
| AT3G50230 | 1.75 | 3.77E-04 | 4.06E-03 | At3g50230 | Protein kinase domain-containing protein |
| AT5G40510 | 1.74 | 1.29E-12 | 1.07E-10 | At5g40510 | Sucrose cleavage protein-like |
| AT1G04425 | 1.74 | 3.54E-07 | 1.02E-05 | | |
| AT2G01918 | 1.73 | 6.16E-08 | 2.11E-06 | At2g01918 | PQL3 |
| AT1G04540 | 1.73 | 9.07E-06 | 1.73E-04 | T1G11.21 | T1G11.21 protein |
| AT1G02650 | 1.71 | 1.95E-03 | 1.54E-02 | At1g02650 | Tetratricopeptide repeat (TPR)-like superfamily protein |
| AT4G31615 | 1.71 | 1.15E-04 | 1.51E-03 | REM2 | B3 domain-containing protein REM2 (Protein REPRODUCTIVE MERISTEM 2) |
| AT5G26290 | 1.71 | 1.67E-05 | 2.93E-04 | RAMGAP | TRAF-like family protein |
| AT4G11210 | 1.70 | 3.23E-04 | 3.60E-03 | DIR14 | Dirigent protein 14 (AtDIR14) |
| AT1G10657 | 1.70 | 7.89E-05 | 1.11E-03 | DEG17 | Transmembrane protein |
| AT5G64040 | 1.69 | 2.27E-40 | 2.21E-37 | PSAN | Photosystem I reaction center subunit PSI-N, chloroplast, putative / PSI-N, putative (PSAN) |
| AT5G26220 | 1.69 | 8.28E-05 | 1.15E-03 | GGCT2 | Gamma-glutamylcyclotransferase (EC 4.3.2.9) |
| AT1G56710 | 1.68 | 5.04E-04 | 5.17E-03 | At1g56710 | (thale cress) hypothetical protein |
| AT4G12545 | 1.68 | 8.55E-06 | 1.64E-04 | AIR1B | Putative lipid-binding protein AIR1B |
| AT5G26700 | 1.67 | 5.66E-03 | 3.50E-02 | At5g26700 | Probable germin-like protein subfamily 2 member 5 |
| AT1G58684 | 1.66 | 4.01E-04 | 4.27E-03 | At1g58684 | XW6 |
| ArthCt117 | 1.66 | 1.59E-05 | 2.81E-04 | | |
| AT1G05660 | 1.66 | 1.09E-04 | 1.45E-03 | At1g05660 | Polygalacturonase |
| AT3G27690 | 1.65 | 1.56E-31 | 8.94E-29 | LHCB23 | Chlorophyll a-b binding protein, chloroplastic |
| AT1G18300 | 1.65 | 1.95E-03 | 1.54E-02 | NUDT4 | Nudix hydrolase 4 |
| AT3G22540 | 1.64 | 7.20E-05 | 1.02E-03 | At3g22540 | (thale cress) hypothetical protein |
| AT3G59930 | 1.64 | 2.13E-03 | 1.65E-02 | At3g59930 | Defensin-like protein 206 |
| AT4G37700 | 1.60 | 1.08E-03 | 9.66E-03 | At4g37700 | Uncharacterized protein |
| AT1G58290 | 1.59 | 1.79E-32 | 1.12E-29 | HEMA1 | Glutamyl-tRNA reductase 1, chloroplastic |
| AT5G26320 | 1.59 | 6.41E-06 | 1.28E-04 | At5g26320 | MATH domain-containing protein |
| AT1G17147 | 1.59 | 2.61E-03 | 1.93E-02 | VQ1 | VQ motif-containing protein 1 (AtVQ1) |
| AT4G11610 | 1.58 | 1.49E-08 | 5.95E-07 | MCTP7 | C2 calcium/lipid-binding plant phosphoribosyltransferase family protein |
| AT3G23510 | 1.58 | 5.32E-10 | 2.86E-08 | At3g23510 | Amine oxidase domain-containing protein |
| AT5G47980 | 1.58 | 2.19E-04 | 2.59E-03 | BAHD1 | BAHD acyltransferase At5g47980 (EC 2.3.1.-) |
| AT3G27500 | 1.58 | 8.75E-04 | 8.12E-03 | At3g27500 | Cysteine/Histidine-rich C1 domain family protein |
| AT2G20670 | 1.57 | 1.03E-04 | 1.38E-03 | At2g20670 | (thale cress) hypothetical protein |
| AT3G49190 | 1.57 | 6.85E-03 | 4.04E-02 | WSD4 | Wax ester synthase/diacylglycerol acyltransferase 4 |
| AT2G47750 | 1.57 | 2.95E-08 | 1.09E-06 | GH3.9 | Putative indole-3-acetic acid-amido synthetase GH3.9 |
| AT4G33790 | 1.57 | 3.68E-05 | 5.79E-04 | CER4 | Fatty acyl-CoA reductase |
| AT3G45860 | 1.55 | 2.19E-03 | 1.68E-02 | CRK4 | Cysteine-rich receptor-like protein kinase 4 |
| AT4G33880 | 1.55 | 7.48E-03 | 4.32E-02 | RSL2 | Transcription factor RSL2 |
| AT1G23390 | 1.55 | 9.04E-04 | 8.32E-03 | At1g23390 | F-box/Kelch repeat-containing F-box family protein |
| ArthCp012 | 1.54 | 3.50E-04 | 3.85E-03 | | |
| AT3G46400 | 1.54 | 1.05E-03 | 9.42E-03 | At3g46400 | Leucine-rich repeat protein kinase family protein |
| AT3G01330 | 1.54 | 1.37E-07 | 4.31E-06 | E2FF | E2F transcription factor-like E2FF |
| AT4G26530 | 1.53 | 5.73E-15 | 6.93E-13 | FBA5 | Fructose-bisphosphate aldolase 5, cytosolic |
| AT1G67035 | 1.53 | 1.89E-04 | 2.30E-03 | F1O19.9 | F1O19.9 protein |
| AT5G10570 | 1.53 | 4.70E-03 | 3.03E-02 | BHLH61 | Transcription factor bHLH61 |
| AT3G23880 | 1.53 | 7.49E-09 | 3.20E-07 | At3g23880 | F-box/kelch-repeat protein At3g23880 |
| AT4G00880 | 1.52 | 3.39E-04 | 3.75E-03 | SAUR31 | AT4g00880 protein |
| ArthCt089 | 1.51 | 5.78E-07 | 1.58E-05 | | |
| AT1G73540 | 1.51 | 1.70E-03 | 1.37E-02 | At1g73540 | Nudix hydrolase domain-containing protein |
| AT5G36920 | 1.51 | 2.14E-03 | 1.66E-02 | STMP9 | Secreted transmembrane peptide 9 |
| AT3G32040 | 1.50 | 8.06E-04 | 7.61E-03 | At3g32040 | Geranylgeranyl pyrophosphate synthase 12, chloroplastic |
| AT3G10150 | 1.48 | 1.10E-05 | 2.06E-04 | PAP16 | Purple acid phosphatase 16 |
| AT4G36060 | 1.48 | 6.77E-04 | 6.62E-03 | BHLH11 | Basic helix-loop-helix |
| AT2G32100 | 1.48 | 5.53E-08 | 1.93E-06 | OFP16 | Transcription repressor OFP16 |
| AT3G08375 | 1.48 | 3.06E-04 | 3.44E-03 | | |
| AT3G47070 | 1.47 | 4.27E-19 | 8.38E-17 | F13I12.120 | Thylakoid soluble phosphoprotein |
| AT2G29290 | 1.47 | 4.53E-11 | 3.02E-09 | At2g29290 | Tropinone reductase homolog At2g29290 |
| AT3G16440 | 1.47 | 4.81E-17 | 7.67E-15 | MLP-300B | Myrosinase-binding protein-like protein-300B |
| AT5G42700 | 1.46 | 5.40E-05 | 7.98E-04 | MJB21.7 | AP2/B3-like transcriptional factor family protein |
| AT4G33120 | 1.46 | 2.70E-05 | 4.45E-04 | At4g33120 | Uncharacterized protein |
| ArthCp024 | 1.46 | 2.34E-04 | 2.75E-03 | | |
| AT1G75620 | 1.46 | 9.36E-04 | 8.57E-03 | At1g75620 | F10A5.18 |
| AT5G42510 | 1.46 | 1.13E-04 | 1.50E-03 | DIR1 | Dirigent protein 1 |
| AT3G62960 | 1.45 | 1.67E-04 | 2.07E-03 | GRXC14 | Glutaredoxin-C14 |

| | | | | | |
|---|---|---|---|---|---|
| AT4G13410 | 1.43 | 5.92E-06 | 1.20E-04 | ATCSLA15 | Nucleotide-diphospho-sugar transferases superfamily protein |
| AT3G55630 | 1.43 | 3.39E-18 | 6.15E-16 | At3g55630 | Folylpolyglutamate synthase |
| AT4G00680 | 1.43 | 8.05E-04 | 7.60E-03 | At4g00680 | ADF-H domain-containing protein |
| AT1G27140 | 1.42 | 4.99E-04 | 5.13E-03 | GSTU14 | Glutathione S-transferase U14 |
| AT4G12870 | 1.42 | 6.67E-04 | 6.54E-03 | At4g12870 | Gamma interferon responsive lysosomal thiol |
| AT4G34400 | 1.42 | 5.04E-03 | 3.19E-02 | At4g34400 | B3 domain-containing protein At4g34400 |
| AT3G18450 | 1.42 | 5.20E-08 | 1.83E-06 | PCR5 | Protein PLANT CADMIUM RESISTANCE 5 |
| AT2G25200 | 1.42 | 5.70E-04 | 5.74E-03 | At2g25200 | At2g25200 |
| AT4G25820 | 1.42 | 1.84E-11 | 1.31E-09 | XTH14 | Xyloglucan endotransglucosylase/hydrolase protein 14 |
| AT1G29600 | 1.42 | 3.34E-03 | 2.34E-02 | At1g29600 | Zinc finger C-x8-C-x5-C-x3-H type family protein |
| AT2G42365 | 1.42 | 9.24E-04 | 8.47E-03 | | |
| ArthCp019 | 1.41 | 1.92E-03 | 1.52E-02 | | |
| AT1G75250 | 1.41 | 1.25E-04 | 1.63E-03 | RL6 | Protein RADIALIS-like 6 |
| AT2G26500 | 1.41 | 1.11E-23 | 3.65E-21 | petM | Cytochrome b6f complex subunit |
| AT3G16670 | 1.41 | 8.49E-05 | 1.17E-03 | At3g16670 | |
| AT4G19980 | 1.40 | 7.38E-08 | 2.48E-06 | At4g19980 | |
| AT1G65490 | 1.40 | 4.17E-16 | 5.79E-14 | STMP5 | Transmembrane protein |
| AT5G45820 | 1.40 | 3.51E-05 | 5.58E-04 | CIPK20 | CBL-interacting serine/threonine-protein kinase 20 |
| AT2G43920 | 1.40 | 3.94E-03 | 2.65E-02 | At2g43920 | HOL2 |
| AT2G34620 | 1.40 | 9.41E-17 | 1.43E-14 | MTERF10 | Mitochondrial transcription termination factor family protein |
| AT1G14220 | 1.40 | 5.16E-05 | 7.70E-04 | At1g14220 | |
| AT5G35525 | 1.40 | 1.73E-03 | 1.39E-02 | PCR3 | Protein PLANT CADMIUM RESISTANCE 3 |
| AT4G29180 | 1.39 | 2.45E-03 | 1.83E-02 | RHS16 | Root hair specific 16 |
| AT1G53520 | 1.38 | 1.90E-15 | 2.43E-13 | FAP3 | Fatty-acid-binding protein 3, chloroplastic |
| AT5G48110 | 1.38 | 1.75E-04 | 2.16E-03 | TPS20 | Inactive terpenoid synthase 20, chloroplastic |
| AT5G63270 | 1.38 | 1.50E-03 | 1.25E-02 | At5g63270 | |
| AT5G47330 | 1.38 | 4.82E-03 | 3.09E-02 | At5g47330 | Uncharacterized protein |
| AT5G51800 | 1.38 | 4.36E-03 | 2.86E-02 | MIO24.6 | Gb\|AAB80672.1 |
| AT5G42630 | 1.37 | 9.23E-11 | 5.70E-09 | KAN4 | Probable transcription factor KAN4 |
| AT4G02270 | 1.37 | 1.31E-09 | 6.56E-08 | SRPP | Protein SEED AND ROOT HAIR PROTECTIVE PROTEIN |
| ArthCp023 | 1.37 | 6.65E-04 | 6.53E-03 | | |
| AT1G68520 | 1.37 | 2.10E-06 | 4.88E-05 | At1g68520 | Uncharacterized protein |
| AT5G07325 | 1.36 | 1.66E-03 | 1.35E-02 | | |
| AT1G13430 | 1.36 | 1.98E-04 | 2.39E-03 | STO9 | Cytosolic sulfotransferase 9 |
| AT4G12830 | 1.36 | 2.08E-10 | 1.20E-08 | At4g12830 | AT4g12830/T20K18_180 |
| AT4G08410 | 1.36 | 6.56E-06 | 1.31E-04 | EXT8 | Extensin-like protein |
| AT1G06350 | 1.36 | 1.09E-03 | 9.70E-03 | At1g06350 | Delta-9 desaturase-like 4 protein |
| AT1G13610 | 1.35 | 5.10E-04 | 5.22E-03 | F21F23.4 | F21F23.4 protein |
| AT4G01270 | 1.35 | 2.30E-05 | 3.86E-04 | At4g01270 | RING-type domain-containing protein |
| AT4G30320 | 1.35 | 1.75E-04 | 2.15E-03 | At4g30320 | |
| AT1G72920 | 1.35 | 4.15E-03 | 2.76E-02 | At1g72920 | Similar to part of disease resistance protein |
| AT1G13260 | 1.33 | 6.95E-05 | 9.90E-04 | RAV1 | AP2/ERF and B3 domain-containing transcription factor RAV1 |
| AT3G22210 | 1.33 | 8.19E-03 | 4.64E-02 | At3g22210 | |
| AT3G46880 | 1.33 | 7.75E-04 | 7.37E-03 | T6H20.90 | Uncharacterized protein T6H20.90 |
| AT2G04790 | 1.33 | 1.00E-10 | 6.16E-09 | At2g04790 | PTB domain engulfment adapter |
| AT5G61590 | 1.33 | 1.91E-05 | 3.28E-04 | ERF107 | Ethylene-responsive transcription factor ERF107 |
| AT3G49790 | 1.33 | 4.74E-05 | 7.16E-04 | At3g49790 | |
| AT5G24105 | 1.32 | 9.67E-06 | 1.84E-04 | AGP41 | Arabinogalactan protein 41 |
| AT2G27402 | 1.32 | 3.50E-04 | 3.85E-03 | At2g27402 | Uncharacterized protein |
| AT3G01500 | 1.32 | 1.32E-08 | 5.35E-07 | At3g01500 | Carbonic anhydrase |
| AT4G01883 | 1.32 | 3.33E-08 | 1.21E-06 | At4g01883 | Coenzyme Q-binding protein COQ10 START domain-containing protein |
| AT1G02820 | 1.32 | 1.11E-08 | 4.56E-07 | LEA2 | Late embryogenis abundant protein 2 |
| AT4G26320 | 1.31 | 1.62E-04 | 2.02E-03 | AGP13 | Arabinogalactan protein 13 |
| AT3G06020 | 1.31 | 1.71E-06 | 4.03E-05 | FAF4 | Protein FANTASTIC FOUR 4 |
| AT4G34800 | 1.31 | 5.20E-03 | 3.28E-02 | At4g34800 | Uncharacterized protein |
| AT3G50560 | 1.31 | 6.19E-17 | 9.55E-15 | At3g50560 | AT3g50560/T20E23_160 |
| AT4G15165 | 1.31 | 4.40E-03 | 2.89E-02 | PAC2 | Putative proteasome subunit alpha type-4-B |
| AT5G59080 | 1.30 | 1.57E-07 | 4.88E-06 | At5g59080 | Uncharacterized protein |
| AT5G16080 | 1.30 | 1.31E-03 | 1.12E-02 | CXE17 | Probable carboxylesterase 17 |
| AT5G62280 | 1.30 | 1.09E-07 | 3.53E-06 | At5g62280 | |
| AT1G31330 | 1.30 | 2.09E-25 | 7.81E-23 | PSAF | Photosystem I reaction center subunit III, chloroplastic |
| AT1G69523 | 1.30 | 8.83E-05 | 1.21E-03 | At1g69523 | Methyltransferase type 11 domain-containing protein |
| AT3G61898 | 1.30 | 2.99E-03 | 2.14E-02 | At3g61898 | Transmembrane protein |
| AT2G37380 | 1.30 | 1.17E-08 | 4.78E-07 | MAKR3 | Probable membrane-associated kinase regulator 3 |
| AT1G73830 | 1.30 | 2.02E-07 | 6.13E-06 | At1g73830 | BHLH domain-containing protein |
| AT2G20750 | 1.30 | 4.69E-08 | 1.68E-06 | EXPB1 | Expansin B1 |
| AT5G60680 | 1.29 | 3.09E-04 | 3.47E-03 | At5g60680 | Senescence regulator |
| AT1G12740 | 1.29 | 3.96E-05 | 6.17E-04 | CYP87A2 | Cytochrome P450, family 87, subfamily A, polypeptide 2 |
| AT1G30380 | 1.28 | 3.33E-21 | 8.19E-19 | At1g30380 | PSI-K |
| AT3G10680 | 1.28 | 6.34E-04 | 6.27E-03 | At3g10680 | SHSP domain-containing protein |
| AT5G19190 | 1.27 | 1.60E-10 | 9.36E-09 | At5g19190 | |
| AT1G12010 | 1.26 | 2.20E-05 | 3.71E-04 | At1g12010 | 1-aminocyclopropane-1-carboxylate oxidase 3 |
| AT2G24400 | 1.26 | 8.06E-04 | 7.61E-03 | At2g24400 | SAUR-like auxin-responsive protein family |
| AT5G04190 | 1.26 | 1.79E-04 | 2.20E-03 | PKS4 | Protein PHYTOCHROME KINASE SUBSTRATE 4 |
| AT4G04955 | 1.26 | 8.63E-13 | 7.41E-11 | ALN | allantoinase |
| AT4G08390 | 1.25 | 5.30E-17 | 8.32E-15 | SAPX | L-ascorbate peroxidase |
| AT5G45680 | 1.25 | 7.69E-18 | 1.35E-15 | FKBP13 | Peptidyl-prolyl cis-trans isomerase FKBP13, chloroplastic |
| AT1G31950 | 1.25 | 4.11E-05 | 6.37E-04 | At1g31950 | Terpenoid cyclases/Protein prenyltransferases superfamily protein |
| AT5G08050 | 1.25 | 4.73E-23 | 1.41E-20 | RIQ1 | Uncharacterized protein F13G24.250 |
| AT1G09450 | 1.25 | 1.04E-03 | 9.37E-03 | HASPIN | Serine/threonine-protein kinase haspin homolog |

| | | | | | |
|---|---|---|---|---|---|
| AT1G01390 | 1.24 | 1.36E-05 | 2.46E-04 | At1g01390 | Glycosyltransferase |
| AT5G40730 | 1.24 | 5.18E-14 | 5.48E-12 | AGP24 | Arabinogalactan protein 24 |
| AT1G20480 | 1.24 | 1.11E-03 | 9.87E-03 | 4CLL2 | 4-coumarate--CoA ligase-like 2 |
| AT3G14850 | 1.24 | 1.20E-05 | 2.21E-04 | TBL41 | Protein trichome birefringence-like 41 |
| AT4G19130 | 1.24 | 3.47E-03 | 2.42E-02 | RPA1E | Replication protein A subunit |
| AT1G50110 | 1.24 | 2.69E-05 | 4.44E-04 | BCAT6 | Branched-chain-amino-acid aminotransferase |
| AT1G50050 | 1.23 | 3.17E-14 | 3.46E-12 | At1g50050 | CAP |
| AT3G16150 | 1.23 | 4.50E-07 | 1.25E-05 | At3g16150 | Probable isoaspartyl peptidase/L-asparaginase 2 |
| AT5G17350 | 1.23 | 4.75E-03 | 3.06E-02 | At5g17350 | |
| AT4G36880 | 1.23 | 6.79E-03 | 4.02E-02 | At4g36880 | |
| AT1G70460 | 1.23 | 3.19E-05 | 5.15E-04 | At1g70460 | non-specific serine/threonine protein kinase |
| AT1G13650 | 1.23 | 5.87E-06 | 1.19E-04 | At1g13650 | Uncharacterized protein |
| AT2G25880 | 1.22 | 6.39E-11 | 4.10E-09 | AUR2 | Aurora kinase |
| AT1G62770 | 1.22 | 1.83E-07 | 5.61E-06 | PMEI9 | Pectinesterase inhibitor 9 |
| AT2G29890 | 1.22 | 2.26E-08 | 8.49E-07 | At2g29890 | Putative villin 1 VLN1 |
| AT1G49010 | 1.22 | 1.87E-09 | 8.98E-08 | MYBS1 | At1g49010 |
| AT1G75030 | 1.22 | 6.41E-03 | 3.85E-02 | At1g75030 | TLP-3 |
| AT1G21500 | 1.22 | 5.56E-17 | 8.65E-15 | At1g21500 | AT1G21500 protein |
| AT5G06250 | 1.21 | 7.64E-03 | 4.40E-02 | DPA4 | AP2/B3-like transcriptional factor family protein |
| AT3G54770 | 1.21 | 1.11E-03 | 9.84E-03 | ARP1 | RNA-binding |
| AT2G43375 | 1.21 | 6.51E-04 | 6.42E-03 | | |
| AT1G06830 | 1.21 | 8.51E-03 | 4.78E-02 | GRXS11 | Monothiol glutaredoxin-S11 |
| AT1G75166 | 1.20 | 3.19E-03 | 2.25E-02 | | |
| AT1G53860 | 1.20 | 4.19E-03 | 2.78E-02 | At1g53860 | At1g53860 |
| AT2G27830 | 1.20 | 1.75E-05 | 3.06E-04 | At2g27830 | Uncharacterized protein |
| AT1G07270 | 1.20 | 2.64E-03 | 1.94E-02 | CDC6B | Cell division control protein |
| AT3G44970 | 1.20 | 1.32E-03 | 1.13E-02 | At3g44970 | Cytochrome P450 superfamily protein |
| AT3G06840 | 1.20 | 2.17E-05 | 3.67E-04 | At3g06840 | |
| AT3G58850 | 1.20 | 5.48E-06 | 1.12E-04 | PAR2 | Transcription factor PAR2 |
| AT3G14260 | 1.20 | 2.69E-04 | 3.09E-03 | At3g14260 | |
| AT5G56550 | 1.19 | 1.78E-05 | 3.09E-04 | OXS3 | Protein OXIDATIVE STRESS 3 |
| AT2G40300 | 1.19 | 3.08E-06 | 6.77E-05 | FER4 | Ferritin-4, chloroplastic |
| AT3G46490 | 1.19 | 2.49E-03 | 1.86E-02 | F12A12.10 | Uncharacterized protein F12A12.10 |
| AT5G24180 | 1.19 | 6.27E-03 | 3.78E-02 | At5g24180 | Gb\|AAD29063.1 |
| AT5G56840 | 1.19 | 1.72E-03 | 1.39E-02 | At5g56840 | |
| AT4G36410 | 1.19 | 1.20E-05 | 2.21E-04 | UBC17 | Probable ubiquitin-conjugating enzyme E2 17 |
| AT2G30230 | 1.19 | 5.75E-03 | 3.53E-02 | At2g30230 | |
| AT5G53030 | 1.19 | 1.25E-03 | 1.08E-02 | MNB8.9 | Uncharacterized protein |
| AT5G63160 | 1.19 | 5.09E-12 | 3.90E-10 | BT1 | BTB and TAZ domain protein 1 |
| AT4G10340 | 1.18 | 7.82E-22 | 2.08E-19 | LHCB5 | Chlorophyll a-b binding protein CP26, chloroplastic |
| AT5G10150 | 1.18 | 7.02E-13 | 6.12E-11 | SOK2 | UPSTREAM OF FLC protein |
| AT5G21940 | 1.18 | 3.57E-05 | 5.64E-04 | O3L1 | Protein OXIDATIVE STRESS 3 LIKE 1 |
| AT3G05730 | 1.18 | 2.14E-05 | 3.62E-04 | At3g05730 | |
| AT1G78450 | 1.18 | 6.92E-04 | 6.72E-03 | At1g78450 | Uncharacterized protein At1g78450 |
| AT5G38410 | 1.18 | 2.26E-13 | 2.16E-11 | RBCS3B | Ribulose bisphosphate carboxylase small subunit, chloroplastic |
| AT1G29025 | 1.18 | 5.75E-03 | 3.53E-02 | At1g29025 | F1K23.2 |
| AT3G59250 | 1.18 | 4.21E-04 | 4.44E-03 | At3g59250 | F-box/LRR-repeat protein At3g59250 |
| AT3G62430 | 1.18 | 1.94E-03 | 1.54E-02 | At3g62430 | F-box protein At3g62430 |
| AT3G42725 | 1.18 | 6.35E-03 | 3.82E-02 | At3g42725 | Membrane lipoprotein |
| AT4G34970 | 1.18 | 1.38E-05 | 2.50E-04 | ADF9 | Actin-depolymerizing factor 9 |
| AT2G43560 | 1.18 | 3.59E-19 | 7.21E-17 | At2g43560 | peptidylprolyl isomerase |
| AT1G52220 | 1.18 | 5.64E-19 | 1.09E-16 | CURT1C | CURVATURE THYLAKOID protein |
| AT5G02350 | 1.17 | 4.76E-04 | 4.94E-03 | At5g02350 | Cysteine/Histidine-rich C1 domain family protein |
| ArthCp020 | 1.17 | 3.69E-07 | 1.05E-05 | | |
| AT1G68120 | 1.17 | 4.29E-03 | 2.83E-02 | At1g68120 | GAGA-binding transcriptional activator |
| AT3G63470 | 1.17 | 6.82E-03 | 4.03E-02 | SCPL40 | Serine carboxypeptidase-like 40 |
| AT1G16640 | 1.17 | 1.18E-03 | 1.04E-02 | At1g16640 | B3 domain-containing protein At1g16640 |
| AT5G14150 | 1.17 | 1.95E-04 | 2.35E-03 | At5g14150 | DUF642 domain-containing protein |
| AT3G20470 | 1.17 | 7.78E-04 | 7.39E-03 | At3g20470 | Uncharacterized protein |
| AT1G55960 | 1.17 | 8.75E-14 | 8.86E-12 | At1g55960 | Polyketide cyclase/dehydrase and lipid transport superfamily protein |
| AT3G07350 | 1.17 | 2.48E-03 | 1.85E-02 | At3g07350 | Uncharacterized protein |
| AT3G19320 | 1.17 | 6.24E-03 | 3.77E-02 | At3g19320 | Leucine-rich repeat |
| AT1G49910 | 1.17 | 2.44E-04 | 2.85E-03 | BUB3.2 | Mitotic checkpoint protein BUB3.2 |
| AT1G58150 | 1.17 | 5.37E-03 | 3.36E-02 | T18I24.6 | Phosphoglycerate kinase |
| AT5G33370 | 1.17 | 4.22E-05 | 6.52E-04 | At5g33370 | GDSL esterase/lipase At5g33370 |
| AT1G16060 | 1.16 | 9.70E-04 | 8.82E-03 | ADAP | ARIA-interacting double AP2 domain protein |
| AT4G24175 | 1.16 | 1.06E-06 | 2.69E-05 | At4g24175 | Kinesin-like protein |
| AT5G56120 | 1.16 | 2.43E-07 | 7.31E-06 | At5g56120 | Uncharacterized protein |
| AT3G45430 | 1.16 | 4.25E-03 | 2.81E-02 | LECRK15 | Probable L-type lectin-domain containing receptor kinase I.5 |
| AT1G19450 | 1.15 | 3.13E-08 | 1.14E-06 | At1g19450 | |
| AT5G57780 | 1.15 | 5.71E-12 | 4.32E-10 | P1R1 | At5g57780 |
| AT5G02160 | 1.15 | 1.25E-12 | 1.05E-10 | T7H20_210 | Uncharacterized protein T7H20_210 |
| AT1G54820 | 1.15 | 9.98E-07 | 2.55E-05 | At1g54820 | Protein kinase superfamily protein |
| AT1G13300 | 1.15 | 5.50E-05 | 8.11E-04 | At1g13300 | HTH myb-type domain-containing protein |
| AT4G32690 | 1.15 | 8.21E-06 | 1.59E-04 | GLB3 | Two-on-two hemoglobin-3 |
| AT1G11460 | 1.15 | 6.36E-03 | 3.83E-02 | At1g11460 | WAT1-related protein At1g11460 |
| AT1G15820 | 1.15 | 8.83E-18 | 1.53E-15 | Lhcb6 | Chlorophyll a-b binding protein, chloroplastic |
| AT1G66800 | 1.14 | 7.49E-03 | 4.32E-02 | At1g66800 | NAD |
| AT1G05300 | 1.14 | 3.58E-03 | 2.47E-02 | ZIP5 | Zinc transporter 5 |
| AT5G58750 | 1.14 | 9.77E-05 | 1.33E-03 | PRISE | NAD |

| | | | | | |
|---|---|---|---|---|---|
| AT4G30610 | 1.13 | 7.16E-07 | 1.90E-05 | SCPL24 | Serine carboxypeptidase 24 |
| AT1G12900 | 1.13 | 1.23E-14 | 1.44E-12 | GAPA-2 | Glyceraldehyde 3-phosphate dehydrogenase A subunit 2 |
| AT5G09978 | 1.13 | 8.76E-04 | 8.12E-03 | PEP7 | Elicitor peptide 7 |
| AT4G25050 | 1.13 | 2.01E-17 | 3.34E-15 | ACP4 | Acyl carrier protein 4 |
| AT3G29670 | 1.13 | 2.01E-04 | 2.42E-03 | PMAT2 | Phenolic glucoside malonyltransferase 2 |
| AT2G27420 | 1.13 | 6.68E-06 | 1.33E-04 | At2g27420 | Cysteine proteinase |
| AT3G01550 | 1.13 | 1.25E-04 | 1.63E-03 | PPT2 | Phosphoenolpyruvate |
| AT3G13590 | 1.12 | 1.79E-03 | 1.43E-02 | At3g13590 | Phorbol-ester/DAG-type domain-containing protein |
| AT1G33840 | 1.12 | 6.20E-03 | 3.75E-02 | At1g33840 | Protein LURP-one-related 15 |
| AT1G75040 | 1.12 | 1.87E-11 | 1.32E-09 | At1g75040 | Pathogenesis-related protein 5 |
| AT1G52245 | 1.12 | 2.70E-03 | 1.97E-02 | F9I5.13 | Dynein light chain |
| AT5G52900 | 1.12 | 9.81E-04 | 8.92E-03 | MAKR6 | Probable membrane-associated kinase regulator 6 |
| AT4G11280 | 1.12 | 6.97E-04 | 6.76E-03 | ACS6 | 1-aminocyclopropane-1-carboxylate synthase 6 |
| AT2G05070 | 1.12 | 6.27E-16 | 8.53E-14 | LHCB2.2 | Chlorophyll a-b binding protein 2.2, chloroplastic |
| ArthCp009 | 1.12 | 4.73E-03 | 3.05E-02 | | |
| AT5G49270 | 1.12 | 1.53E-04 | 1.92E-03 | COBL9 | COBRA-like protein 9 |
| AT3G20940 | 1.11 | 1.31E-03 | 1.12E-02 | CYP705A30 | Cytochrome P450, family 705, subfamily A, polypeptide 30 |
| AT4G39970 | 1.11 | 2.25E-10 | 1.29E-08 | At4g39970 | Haloacid dehalogenase-like hydrolase domain-containing protein At4g39970 |
| AT5G56100 | 1.11 | 4.62E-05 | 7.02E-04 | MDA7.16 | Glycine-rich protein / oleosin |
| AT1G68110 | 1.11 | 7.92E-04 | 7.50E-03 | At1g68110 | Putative clathrin assembly protein At1g68110 |
| AT5G05580 | 1.11 | 1.74E-03 | 1.40E-02 | FAD8 | Fatty acid desaturase 8 |
| AT5G38420 | 1.11 | 4.75E-13 | 4.22E-11 | RBCS-2B | Ribulose bisphosphate carboxylase small subunit 2B, chloroplastic |
| AT4G21500 | 1.11 | 4.96E-04 | 5.10E-03 | At4g21500 | At4g21500 |
| AT1G23360 | 1.10 | 6.67E-06 | 1.33E-04 | MENG | 2-phytyl-1,4-beta-naphthoquinone methyltransferase, chloroplastic |
| AT3G56090 | 1.10 | 8.82E-09 | 3.70E-07 | At3g56090 | Ferritin |
| AT1G67780 | 1.10 | 1.51E-03 | 1.26E-02 | At1g67780 | Zinc-finger domain of monoamine-oxidase A repressor R1 protein |
| AT4G40090 | 1.10 | 1.31E-07 | 4.14E-06 | AGP3 | Classical arabinogalactan protein 3 |
| AT5G67190 | 1.10 | 1.46E-08 | 5.82E-07 | ERF010 | Ethylene-responsive transcription factor ERF010 |
| AT5G23870 | 1.10 | 4.53E-05 | 6.92E-04 | PAE9 | Pectin acetylesterase |
| AT5G56540 | 1.10 | 9.50E-07 | 2.46E-05 | AGP14 | Arabinogalactan protein 14 |
| AT1G48330 | 1.10 | 1.89E-05 | 3.25E-04 | At1g48330 | |
| AT5G25810 | 1.09 | 2.93E-07 | 8.55E-06 | TINY | Ethylene-responsive transcription factor TINY |
| AT1G76110 | 1.09 | 1.71E-10 | 9.92E-09 | HMGB9 | High mobility group B protein 9 |
| AT1G09475 | 1.09 | 3.92E-03 | 2.64E-02 | | |
| AT2G18300 | 1.09 | 8.55E-08 | 2.84E-06 | HBI1 | Basic helix-loop-helix |
| AT1G43670 | 1.09 | 2.10E-18 | 3.92E-16 | CYFBP | Fructose-1,6-bisphosphatase, cytosolic |
| AT1G52050 | 1.09 | 5.30E-05 | 7.87E-04 | JAL8 | Jacalin-related lectin 8 |
| AT4G37685 | 1.09 | 3.63E-03 | 2.49E-02 | At4g37685 | Uncharacterized protein |
| AT3G16250 | 1.09 | 1.12E-10 | 6.74E-09 | PNSB3 | Photosynthetic NDH subunit of subcomplex B 3, chloroplastic |
| AT4G17480 | 1.09 | 7.37E-03 | 4.27E-02 | dl4775c | Thioesterase like protein |
| AT3G61870 | 1.08 | 3.12E-16 | 4.56E-14 | F21F14.40 | AT3g61870/F21F14_40 |
| AT2G01590 | 1.08 | 3.93E-09 | 1.74E-07 | CRR3 | Chlororespiratory reduction 3 |
| AT1G77870 | 1.08 | 5.99E-03 | 3.65E-02 | MUB5 | Membrane-anchored ubiquitin-fold protein 5 |
| AT5G23840 | 1.08 | 5.57E-05 | 8.20E-04 | MRO11.12 | MD-2-related lipid recognition domain-containing protein |
| AT3G30122 | 1.08 | 8.19E-04 | 7.69E-03 | | |
| AT1G30520 | 1.08 | 1.30E-06 | 3.22E-05 | AAE14 | Acyl-activating enzyme 14 |
| AT2G24610 | 1.08 | 4.30E-03 | 2.84E-02 | CNGC14 | Cyclic nucleotide-gated channel 14 |
| AT2G02410 | 1.08 | 1.20E-06 | 2.99E-05 | At2g02410 | Uncharacterized protein At2g02410 |
| AT5G62420 | 1.08 | 7.79E-03 | 4.46E-02 | At5g62420 | NADP-dependent oxidoreductase domain-containing protein |
| AT4G21445 | 1.07 | 2.06E-08 | 7.84E-07 | At4g21445 | Receptor-interacting protein |
| AT3G09580 | 1.07 | 2.01E-11 | 1.41E-09 | At3g09580 | |
| AT1G09812 | 1.07 | 4.06E-03 | 2.71E-02 | At1g09812 | Transmembrane protein |
| AT5G17670 | 1.07 | 1.01E-13 | 1.01E-11 | At5g17670 | GPI inositol-deacylase |
| AT5G02680 | 1.07 | 1.34E-03 | 1.14E-02 | At5g02680 | Uncharacterized protein |
| AT3G06145 | 1.06 | 2.00E-04 | 2.41E-03 | At3g06145 | At3g06145 |
| AT5G18660 | 1.06 | 1.49E-14 | 1.72E-12 | DVR | Divinyl chlorophyllide a 8-vinyl-reductase, chloroplastic |
| AT1G52342 | 1.06 | 2.60E-14 | 2.88E-12 | At1g52342 | Uncharacterized protein |
| AT3G15520 | 1.06 | 6.54E-07 | 1.76E-05 | At3g15520 | Cyclophilin-like peptidyl-prolyl cis-trans isomerase family protein |
| AT1G20020 | 1.06 | 1.73E-11 | 1.24E-09 | FNR2 | Ferredoxin--NADP reductase, chloroplastic |
| AT4G30250 | 1.06 | 3.46E-04 | 3.82E-03 | At4g30250 | P-loop containing nucleoside triphosphate hydrolases superfamily protein |
| AT1G22900 | 1.06 | 7.54E-03 | 4.35E-02 | DIR11 | Dirigent protein 11 |
| AT1G61795 | 1.06 | 6.28E-03 | 3.79E-02 | At1g61795 | CRIB domain-containing protein |
| AT2G27480 | 1.06 | 1.60E-03 | 1.32E-02 | At2g27480 | Calcium-binding EF-hand family protein |
| AT1G05820 | 1.06 | 5.40E-03 | 3.38E-02 | SPPL5 | SIGNAL PEPTIDE PEPTIDASE-LIKE 5 |
| AT1G14150 | 1.06 | 3.31E-10 | 1.83E-08 | PNSL2 | Photosynthetic NDH subunit of lumenal location 2, chloroplastic |
| AT1G65370 | 1.06 | 1.77E-05 | 3.08E-04 | At1g65370 | At1g65370/T8F5_15 |
| AT1G74453 | 1.06 | 1.69E-04 | 2.10E-03 | | |
| AT3G47470 | 1.06 | 3.02E-15 | 3.79E-13 | At3g47470 | Chlorophyll a-b binding protein, chloroplastic |
| AT2G40020 | 1.05 | 7.90E-08 | 2.64E-06 | At2g40020 | Nucleolar histone methyltransferase-related protein |
| AT4G21280 | 1.05 | 3.99E-13 | 3.56E-11 | At4g21280 | PSBQA |
| AT1G08380 | 1.05 | 4.56E-17 | 7.34E-15 | PSAO | Photosystem I subunit O |
| AT3G17640 | 1.05 | 1.59E-05 | 2.82E-04 | At3g17640 | Leucine-rich repeat-containing N-terminal plant-type domain-containing protein |
| AT1G32540 | 1.05 | 8.19E-04 | 7.69E-03 | At1g32540 | Zinc finger LSD1-type domain-containing protein |
| AT3G54890 | 1.05 | 2.86E-17 | 4.71E-15 | LHCA1 | Chlorophyll a-b binding protein, chloroplastic |
| AT3G47675 | 1.05 | 8.00E-03 | 4.55E-02 | At3g47675 | Ternary complex factor MIP1 leucine-zipper domain-containing protein |
| AT1G15980 | 1.05 | 4.54E-08 | 1.63E-06 | PNSB1 | Photosynthetic NDH subunit of subcomplex B 1, chloroplastic |
| AT5G27350 | 1.05 | 8.96E-11 | 5.57E-09 | At5g27350 | Major facilitator superfamily |
| AT2G40000 | 1.05 | 2.89E-03 | 2.08E-02 | HSPRO2 | Nematode resistance protein-like HSPRO2 |
| AT2G39470 | 1.04 | 2.27E-12 | 1.85E-10 | PNSL1 | Photosynthetic NDH subunit of lumenal location 1, chloroplastic |
| AT5G54130 | 1.04 | 1.59E-15 | 2.04E-13 | At5g54130 | Endonuclease/exonuclease/phosphatase domain-containing protein |

| | | | | | |
|---|---|---|---|---|---|
| AT3G16660 | 1.04 | 1.01E-03 | 9.14E-03 | At3g16660 | Pollen Ole e 1 allergen and extensin family protein |
| AT5G52570 | 1.04 | 1.34E-07 | 4.24E-06 | BETA-OHASE | Beta-carotene 3-hydroxylase 2, chloroplastic |
| AT1G49880 | 1.04 | 3.14E-04 | 3.51E-03 | ERV1 | Sulfhydryl oxidase |
| AT1G12250 | 1.04 | 8.65E-13 | 7.41E-11 | TL203 | Pentapeptide repeat-containing protein |
| AT1G52827 | 1.04 | 6.14E-04 | 6.09E-03 | CYSTM2 | Protein CYSTEINE-RICH TRANSMEMBRANE MODULE 2 |
| AT4G35030 | 1.03 | 1.01E-04 | 1.37E-03 | At4g35030 | Protein kinase superfamily protein |
| AT2G40330 | 1.03 | 7.52E-03 | 4.34E-02 | PYL6 | Abscisic acid receptor PYL6 |
| AT3G56000 | 1.03 | 2.60E-03 | 1.92E-02 | CSLA14 | Probable glucomannan 4-beta-mannosyltransferase 14 |
| AT1G60600 | 1.03 | 1.54E-10 | 9.07E-09 | ABC4 | 2-carboxy-1,4-naphthoquinone phytyltransferase, chloroplastic |
| AT2G37810 | 1.03 | 5.63E-03 | 3.49E-02 | At2g37810 | Cysteine/Histidine-rich C1 domain family protein |
| AT1G03180 | 1.03 | 6.52E-03 | 3.90E-02 | F15K9.21 | F15k9.21 |
| AT5G58770 | 1.03 | 9.74E-07 | 2.50E-05 | At5g58770 | Dehydrodolichyl diphosphate synthase 2 |
| AT1G16445 | 1.03 | 3.29E-04 | 3.66E-03 | At1g16445 | S-adenosyl-L-methionine-dependent methyltransferases superfamily protein |
| AT1G77590 | 1.03 | 2.53E-10 | 1.44E-08 | LACS9 | Long chain acyl-CoA synthetase 9 |
| AT4G33260 | 1.03 | 7.02E-04 | 6.80E-03 | CDC20-2 | Cell division cycle 20.2, cofactor of APC complex |
| AT4G16880 | 1.03 | 1.12E-03 | 9.95E-03 | dl4470c | Disease resistance RPP5 like protein |
| AT5G11070 | 1.03 | 1.25E-09 | 6.28E-08 | At5g11070 | Uncharacterized protein |
| AT5G36790 | 1.02 | 2.72E-04 | 3.12E-03 | PGLP1B | Phosphoglycolate phosphatase 1B, chloroplastic |
| AT2G05540 | 1.02 | 1.04E-13 | 1.03E-11 | At2g05540 | Glycine-rich protein |
| AT5G03905 | 1.02 | 1.87E-07 | 5.72E-06 | At5g03905 | Iron-sulfur cluster biosynthesis family protein |
| AT1G62780 | 1.02 | 8.41E-15 | 9.91E-13 | At1g62780 | Dimethylallyl, adenosine tRNA methylthiotransferase |
| AT2G45180 | 1.02 | 8.30E-05 | 1.15E-03 | At2g45180 | At2g45180 |
| AT4G38860 | 1.02 | 3.75E-06 | 8.05E-05 | SAUR16 | Protein SMALL AUXIN UP-REGULATED RNA 16 |
| AT3G52480 | 1.02 | 1.33E-04 | 1.70E-03 | At3g52480 | |
| AT3G27660 | 1.02 | 3.91E-04 | 4.17E-03 | At3g27660 | Oleosin |
| AT5G24420 | 1.02 | 3.93E-05 | 6.13E-04 | PGL5 | Probable 6-phosphogluconolactonase 5 |
| AT1G07450 | 1.02 | 1.64E-03 | 1.34E-02 | At1g07450 | Tropinone reductase homolog At1g07450 |
| AT1G20340 | 1.02 | 3.35E-14 | 3.64E-12 | DRT112 | Plastocyanin major isoform, chloroplastic |
| AT5G07580 | 1.02 | 1.64E-06 | 3.91E-05 | ERF106 | Ethylene-responsive transcription factor ERF106 |
| AT1G08115 | 1.02 | 2.55E-03 | 1.89E-02 | | |
| AT4G28270 | 1.02 | 2.38E-09 | 1.12E-07 | RMA2 | E3 ubiquitin-protein ligase RMA2 |
| ArthCp010 | 1.01 | 7.11E-03 | 4.16E-02 | | |
| AT5G02120 | 1.01 | 3.54E-11 | 2.41E-09 | OHP1 | Light-harvesting complex-like protein OHP1, chloroplastic |
| AT3G48360 | 1.01 | 9.36E-06 | 1.78E-04 | BT2 | BTB and TAZ domain protein 2 |
| AT5G65730 | 1.01 | 7.72E-08 | 2.59E-06 | XTH6 | Probable xyloglucan endotransglucosylase/hydrolase protein 6 |
| AT1G52230 | 1.01 | 1.69E-11 | 1.22E-09 | PSAH2 | Photosystem I reaction center subunit VI-2, chloroplastic |
| AT3G07800 | 1.01 | 4.91E-05 | 7.38E-04 | TK1A | Thymidine kinase a |
| AT4G05180 | 1.01 | 5.61E-14 | 5.86E-12 | PSBQ2 | Oxygen-evolving enhancer protein 3-2, chloroplastic |
| AT2G32500 | 1.01 | 6.23E-06 | 1.25E-04 | At2g32500 | Stress responsive alpha-beta barrel domain protein |
| AT5G60660 | 1.01 | 4.43E-08 | 1.59E-06 | PIP2-4 | Probable aquaporin PIP2-4 |
| AT4G00080 | 1.01 | 3.81E-03 | 2.58E-02 | At4g00080 | |
| AT1G35180 | 1.00 | 2.09E-04 | 2.50E-03 | T32G9.28 | Uncharacterized protein T32G9.28 |
| AT1G80440 | 1.00 | 8.74E-07 | 2.28E-05 | At1g80440 | F-box/kelch-repeat protein At1g80440 |
| AT1G60950 | 1.00 | 4.08E-15 | 5.05E-13 | FD2 | Ferredoxin-2, chloroplastic |
| AT4G33000 | 1.00 | 4.05E-06 | 8.58E-05 | At4g33000 | Calcineurin B-like protein |
| AT3G48420 | 1.00 | 4.83E-13 | 4.27E-11 | CBBY | CBBY-like protein |
| AT2G02500 | 1.00 | 6.72E-06 | 1.33E-04 | ISPD | 2-C-methyl-D-erythritol 4-phosphate cytidylyltransferase, chloroplastic |
| AT5G57345 | 1.00 | 2.71E-08 | 1.01E-06 | ATOXR | At5g57345 |
| AT5G18130 | -1.00 | 8.30E-06 | 1.61E-04 | MRG7.9 | Transmembrane protein |
| AT4G15990 | -1.00 | 6.97E-06 | 1.38E-04 | At4g15990 | |
| AT1G35115 | -1.00 | 2.48E-09 | 1.16E-07 | | |
| AT5G65750 | -1.01 | 8.55E-12 | 6.32E-10 | MPA24.10 | oxoglutarate dehydrogenase |
| AT3G29575 | -1.01 | 3.04E-06 | 6.69E-05 | AFP3 | Ninja-family protein AFP3 |
| AT1G60730 | -1.01 | 6.94E-09 | 2.99E-07 | At1g60730 | NAD |
| AT4G16835 | -1.01 | 7.74E-03 | 4.45E-02 | DYW10 | Pentatricopeptide repeat-containing protein At4g16835, mitochondrial |
| AT4G34860 | -1.01 | 1.26E-07 | 4.02E-06 | INVB | Probable alkaline/neutral invertase B |
| AT4G32920 | -1.01 | 4.32E-11 | 2.90E-09 | At4g32920 | AT4g32920/F26P21_40 |
| AT2G32120 | -1.01 | 8.78E-10 | 4.60E-08 | HSP70-8 | Heat shock 70 kDa protein 8 |
| AT1G01470 | -1.01 | 2.13E-09 | 1.01E-07 | LEA14 | Probable desiccation-related protein LEA14 |
| AT2G46270 | -1.01 | 7.29E-13 | 6.33E-11 | At2g46270 | BZIP domain-containing protein |
| AT1G19020 | -1.01 | 3.80E-03 | 2.57E-02 | At1g19020 | CDP-diacylglycerol-glycerol-3-phosphate 3-phosphatidyltransferase |
| AT1G67360 | -1.01 | 4.78E-12 | 3.71E-10 | At1g67360 | REF/SRPP-like protein At1g67360 |
| AT1G51090 | -1.01 | 9.71E-04 | 8.83E-03 | ATHMAD1 | Heavy metal transport/detoxification superfamily protein |
| AT4G25810 | -1.01 | 2.19E-03 | 1.68E-02 | At4g25810 | XTR6 |
| AT4G19090 | -1.01 | 3.73E-03 | 2.55E-02 | At4g19090 | Transmembrane protein, putative |
| AT1G23760 | -1.02 | 3.61E-03 | 2.49E-02 | PGL1 | Polygalacturonase 1 beta-like protein 1 |
| AT2G12190 | -1.02 | 1.03E-04 | 1.38E-03 | At2g12190 | Cytochrome P450 superfamily protein |
| AT2G45220 | -1.02 | 3.82E-06 | 8.19E-05 | PME17 | Probable pectinesterase/pectinesterase inhibitor 17 [Includes: Pectinesterase inhibitor 17 |
| AT5G66080 | -1.02 | 1.01E-06 | 2.58E-05 | APD9 | Protein phosphatase 2C family protein |
| AT1G61890 | -1.02 | 9.31E-11 | 5.73E-09 | At1g61890 | Protein DETOXIFICATION |
| AT1G47405 | -1.02 | 1.80E-04 | 2.21E-03 | | |
| AT3G48020 | -1.02 | 3.23E-05 | 5.20E-04 | At3g48020 | |
| AT5G17420 | -1.02 | 5.02E-03 | 3.19E-02 | CESA7 | Cellulose synthase A catalytic subunit 7 [UDP-forming] |
| AT1G68690 | -1.02 | 8.07E-08 | 2.69E-06 | PERK9 | non-specific serine/threonine protein kinase |
| AT3G24927 | -1.02 | 7.95E-09 | 3.38E-07 | | |
| AT1G77000 | -1.03 | 3.68E-05 | 5.79E-04 | SKP2B | RNI-like superfamily protein |
| AT1G68570 | -1.03 | 2.44E-09 | 1.14E-07 | NPF31 | Major facilitator superfamily protein |
| AT1G51140 | -1.03 | 1.43E-07 | 4.49E-06 | BHLH122 | Transcription factor bHLH122 |
| AT2G20825 | -1.03 | 8.48E-04 | 7.92E-03 | ULT2 | Protein ULTRAPETALA 2 |
| AT5G10300 | -1.03 | 1.24E-04 | 1.62E-03 | MES5 | Methylesterase |

| | | | | | |
|---|---|---|---|---|---|
| AT1G15310 | -1.04 | 1.75E-04 | 2.15E-03 | SRP-54A | Signal recognition particle subunit SRP54 1 |
| AT1G78610 | -1.04 | 1.22E-10 | 7.28E-09 | MSL6 | Mechanosensitive ion channel protein 6 |
| AT4G14370 | -1.04 | 1.32E-04 | 1.69E-03 | DL3225C | Disease resistance protein |
| AT1G62280 | -1.04 | 3.89E-04 | 4.15E-03 | SLAH1 | S-type anion channel SLAH1 |
| AT1G72800 | -1.04 | 3.32E-03 | 2.33E-02 | At1g72800 | RRM domain-containing protein |
| AT1G67365 | -1.04 | 3.76E-05 | 5.90E-04 | | |
| AT1G58270 | -1.04 | 6.18E-06 | 1.24E-04 | At1g58270 | Expressed protein |
| AT5G01670 | -1.04 | 7.21E-09 | 3.10E-07 | At5g01670 | NAD |
| AT4G37970 | -1.04 | 2.06E-06 | 4.80E-05 | CAD6 | Cinnamyl alcohol dehydrogenase 6 |
| AT4G19230 | -1.05 | 2.66E-06 | 5.95E-05 | CYP707A1 | Cytochrome P450, family 707, subfamily A, polypeptide 1 |
| AT1G76590 | -1.05 | 2.10E-07 | 6.34E-06 | At1g76590 | PLATZ transcription factor family protein |
| AT3G14470 | -1.05 | 5.67E-08 | 1.97E-06 | RPPL1 | Putative disease resistance RPP13-like protein 1 |
| AT5G23220 | -1.05 | 9.92E-05 | 1.34E-03 | NIC3 | Nicotinamidase 3 |
| AT5G49330 | -1.06 | 5.43E-04 | 5.49E-03 | At5g49330 | PFG3 |
| AT4G30460 | -1.06 | 1.42E-05 | 2.55E-04 | At4g30460 | Glycine-rich protein |
| AT5G01900 | -1.06 | 6.20E-03 | 3.75E-02 | WRKY62 | Probable WRKY transcription factor 62 |
| AT2G36770 | -1.06 | 2.22E-03 | 1.70E-02 | UGT73C4 | UDP-glycosyltransferase 73C4 |
| AT4G35560 | -1.06 | 5.88E-07 | 1.60E-05 | At4g35560 | Uncharacterized protein At4g35560 |
| AT2G39430 | -1.07 | 3.26E-04 | 3.63E-03 | DIR9 | Dirigent protein 9 |
| AT5G45310 | -1.07 | 2.53E-05 | 4.18E-04 | At5g45310 | Uncharacterized protein |
| AT4G00700 | -1.07 | 1.56E-05 | 2.77E-04 | MCTP9 | Multiple C2 domain and transmembrane region protein 9 |
| AT4G23150 | -1.07 | 8.00E-03 | 4.55E-02 | CRK7 | Cysteine-rich receptor-like protein kinase 7 |
| AT3G20300 | -1.07 | 1.63E-10 | 9.51E-09 | At3g20300 | Uncharacterized protein At3g20300 |
| AT5G06230 | -1.07 | 5.79E-04 | 5.80E-03 | TBL9 | TRICHOME BIREFRINGENCE-LIKE 9 |
| AT5G07010 | -1.08 | 1.53E-04 | 1.92E-03 | SOT15 | Cytosolic sulfotransferase 15 |
| AT2G22470 | -1.08 | 2.87E-06 | 6.38E-05 | AGP2 | Classical arabinogalactan protein 2 |
| AT5G37670 | -1.08 | 5.19E-03 | 3.27E-02 | HSP15.7 | 15.7 kDa heat shock protein, peroxisomal |
| AT5G13910 | -1.08 | 8.85E-10 | 4.63E-08 | LEP | Ethylene-responsive transcription factor LEP |
| AT5G61865 | -1.09 | 1.62E-03 | 1.33E-02 | At5g61865 | Uncharacterized protein |
| AT5G18270 | -1.09 | 1.46E-07 | 4.56E-06 | ANAC087 | NAC domain containing protein 87 |
| AT2G20770 | -1.09 | 1.18E-07 | 3.79E-06 | GCL2 | LanC-like protein GCL2 |
| AT1G22370 | -1.09 | 4.88E-05 | 7.34E-04 | UGT85A5 | UDP-glycosyltransferase 85A5 |
| AT5G01720 | -1.09 | 1.41E-09 | 6.97E-08 | At5g01720 | RNI-like superfamily protein |
| AT2G32190 | -1.10 | 5.35E-03 | 3.35E-02 | ATHCYSTM4 | Cysteine-rich/transmembrane domain A-like protein |
| AT4G23670 | -1.10 | 9.28E-12 | 6.83E-10 | At4g23670 | Bet v I/Major latex protein domain-containing protein |
| AT3G57780 | -1.10 | 1.20E-06 | 2.99E-05 | At3g57780 | Nucleolar-like protein |
| AT2G01390 | -1.10 | 6.78E-05 | 9.70E-04 | At2g01390/At2g01 | Pentatricopeptide repeat-containing protein At2g01390 |
| AT3G05640 | -1.11 | 1.85E-05 | 3.20E-04 | At3g05640 | Probable protein phosphatase 2C 34 |
| AT1G58360 | -1.11 | 3.20E-13 | 2.97E-11 | AAP1 | Amino acid permease 1 |
| AT1G52120 | -1.11 | 1.17E-06 | 2.92E-05 | JAL12 | Jacalin-related lectin 12 |
| AT3G02410 | -1.11 | 6.66E-03 | 3.96E-02 | ICME-LIKE2 | Alpha/beta-Hydrolases superfamily protein |
| AT1G08650 | -1.12 | 7.63E-11 | 4.85E-09 | PPCK1 | Phosphoenolpyruvate carboxylase kinase 1 |
| AT4G01360 | -1.12 | 1.67E-06 | 3.97E-05 | A_IG002N01.26 | A_IG002N01.26 protein |
| AT1G19960 | -1.12 | 5.50E-05 | 8.11E-04 | At1g19960 | Uncharacterized protein |
| AT3G47110 | -1.12 | 4.19E-05 | 6.48E-04 | At3g47110 | Putative receptor-like protein kinase At3g47110 |
| AT5G49645 | -1.12 | 1.54E-03 | 1.28E-02 | At5g49645 | Uncharacterized protein |
| AT5G37540 | -1.12 | 3.50E-04 | 3.85E-03 | At5g37540 | Peptidase A1 domain-containing protein |
| AT4G29190 | -1.12 | 2.97E-07 | 8.65E-06 | At4g29190 | Zinc finger CCCH domain-containing protein 49 |
| AT1G08630 | -1.12 | 1.59E-05 | 2.81E-04 | THA1 | Threonine aldolase 1 |
| AT3G03470 | -1.12 | 5.69E-06 | 1.16E-04 | CYP89A9 | Cytochrome P450 89A9 |
| AT3G16990 | -1.13 | 6.20E-05 | 9.01E-04 | TENA_E | Bifunctional TENA-E protein |
| AT4G21650 | -1.13 | 1.44E-03 | 1.21E-02 | SBT3.13 | Subtilisin-like protease SBT3.13 |
| AT4G10500 | -1.13 | 4.50E-04 | 4.69E-03 | DLO1 | Protein DMR6-LIKE OXYGENASE 1 |
| AT3G50760 | -1.13 | 2.19E-08 | 8.30E-07 | GATL2 | Probable galacturonosyltransferase-like 2 |
| AT1G80120 | -1.13 | 8.78E-08 | 2.91E-06 | At1g80120 | Protein LURP-one-related 5 |
| AT3G49120 | -1.13 | 4.80E-04 | 4.97E-03 | PER34 | Peroxidase 34 |
| AT2G21640 | -1.13 | 6.67E-07 | 1.79E-05 | At2g21640 | Uncharacterized protein |
| AT1G53330 | -1.14 | 1.70E-04 | 2.10E-03 | At1g53330 | Pentatricopeptide repeat |
| AT1G10220 | -1.14 | 1.58E-03 | 1.30E-02 | At1g10220 | ZCF37 |
| AT5G19110 | -1.14 | 7.52E-03 | 4.34E-02 | At5g19110 | Eukaryotic aspartyl protease family protein |
| AT3G08500 | -1.14 | 2.92E-03 | 2.10E-02 | MYB83 | Transcription factor MYB83 |
| AT3G17110 | -1.15 | 6.42E-06 | 1.28E-04 | | |
| AT2G30550 | -1.15 | 2.73E-12 | 2.18E-10 | DALL3 | Alpha/beta-Hydrolases superfamily protein |
| AT1G70830 | -1.15 | 1.84E-05 | 3.17E-04 | MLP28 | MLP-like protein 28 |
| AT5G64430 | -1.16 | 2.20E-09 | 1.05E-07 | At5g64430 | PB1 domain-containing protein |
| AT5G20420 | -1.16 | 4.13E-04 | 4.37E-03 | CLSY2 | SNF2 domain-containing protein CLASSY 2 |
| AT5G13930 | -1.16 | 2.98E-10 | 1.68E-08 | CHS | Chalcone synthase |
| AT3G51750 | -1.16 | 3.05E-06 | 6.71E-05 | At3g51750 | Uncharacterized protein |
| AT2G39050 | -1.16 | 2.22E-12 | 1.81E-10 | At2g39050 | Hydroxyproline-rich glycoprotein family protein |
| AT4G35190 | -1.16 | 8.56E-04 | 7.97E-03 | LOG5 | Cytokinin riboside 5'-monophosphate phosphoribohydrolase |
| AT3G02370 | -1.16 | 4.87E-04 | 5.02E-03 | F16B3.1 | F16B3.1 protein |
| AT5G57050 | -1.17 | 3.55E-13 | 3.24E-11 | At5g57050 | protein-serine/threonine phosphatase |
| AT5G53390 | -1.18 | 8.25E-03 | 4.67E-02 | WSD11 | Wax ester synthase/diacylglycerol acyltransferase 11 |
| AT1G18710 | -1.18 | 4.99E-05 | 7.48E-04 | MYB47 | MYB transcription factor |
| AT2G28650 | -1.18 | 2.61E-04 | 3.02E-03 | At2g28650 | Exocyst subunit Exo70 family protein |
| AT4G28460 | -1.19 | 5.86E-03 | 3.58E-02 | PIP1 | PAMP-induced secreted peptide 1 |
| AT1G24580 | -1.19 | 8.42E-22 | 2.21E-19 | At1g24580 | At1g24580 |
| AT1G54100 | -1.19 | 1.08E-08 | 4.47E-07 | ALDH7B4 | Aldehyde dehydrogenase family 7 member B4 |
| AT1G26240 | -1.20 | 3.72E-17 | 6.03E-15 | EXT19 | Proline-rich extensin-like family protein |
| AT1G56120 | -1.20 | 1.51E-03 | 1.26E-02 | At1g56120 | Leucine-rich repeat transmembrane protein kinase |

| | | | | | |
|---|---|---|---|---|---|
| AT4G31405 | -1.20 | 1.71E-03 | 1.38E-02 | At4g31405 | Uncharacterized protein |
| AT3G51860 | -1.21 | 3.06E-04 | 3.44E-03 | CAX3 | Vacuolar cation/proton exchanger |
| AT1G80130 | -1.21 | 2.96E-05 | 4.84E-04 | At1g80130 | |
| AT2G43050 | -1.21 | 1.21E-07 | 3.88E-06 | PME16 | Probable pectinesterase/pectinesterase inhibitor 16 [Includes: Pectinesterase inhibitor 16 |
| AT1G61420 | -1.22 | 9.03E-04 | 8.32E-03 | At1g61420 | S-locus lectin protein kinase family protein |
| AT1G63240 | -1.22 | 3.14E-10 | 1.75E-08 | At1g63240 | MBD domain-containing protein |
| AT3G19620 | -1.22 | 7.04E-04 | 6.82E-03 | BXL5 | Probable beta-D-xylosidase 5 |
| AT1G09045 | -1.22 | 2.80E-03 | 2.03E-02 | | |
| AT3G04040 | -1.22 | 3.33E-04 | 3.70E-03 | At3g04040 | |
| AT1G13340 | -1.23 | 1.84E-05 | 3.17E-04 | ISTL6 | Regulator of Vps4 activity in the MVB pathway protein |
| AT2G28840 | -1.23 | 1.50E-20 | 3.44E-18 | XBAT31 | RING-type domain-containing protein |
| AT2G12400 | -1.23 | 1.16E-17 | 1.98E-15 | At2g12400 | Uncharacterized protein At2g12400 |
| AT4G38410 | -1.23 | 3.79E-03 | 2.57E-02 | At4g38410 | At4g38410 |
| AT1G78955 | -1.23 | 7.70E-05 | 1.08E-03 | CAMS1 | Camelliol C synthase 1 |
| AT2G33380 | -1.24 | 1.31E-03 | 1.12E-02 | PXG3 | Probable peroxygenase 3 |
| AT3G53230 | -1.24 | 2.48E-15 | 3.15E-13 | CDC48D | Cell division control protein 48 homolog D |
| AT1G02520 | -1.24 | 9.46E-13 | 8.07E-11 | ABCB11 | ABC transporter B family member 11 |
| AT3G14560 | -1.24 | 2.76E-04 | 3.15E-03 | At3g14560 | Uncharacterized protein |
| AT3G48450 | -1.24 | 6.86E-12 | 5.11E-10 | At3g48450 | |
| AT1G51420 | -1.25 | 1.69E-05 | 2.96E-04 | SPP1 | Sucrose-phosphatase |
| AT3G14590 | -1.25 | 3.05E-10 | 1.70E-08 | NTMC2T62 | Calcium-dependent lipid-binding |
| AT1G23200 | -1.25 | 1.67E-03 | 1.36E-02 | PME6 | Probable pectinesterase/pectinesterase inhibitor 6 [Includes: Pectinesterase inhibitor 6 |
| AT5G53990 | -1.25 | 1.00E-03 | 9.11E-03 | UGT79B9 | UDP-glycosyltransferase 79B9 |
| AT5G03190 | -1.25 | 2.41E-18 | 4.47E-16 | At5g03190 | |
| AT3G28270 | -1.26 | 4.69E-07 | 1.30E-05 | At3g28270 | UPF0496 protein At3g28270 |
| AT5G57010 | -1.26 | 8.60E-03 | 4.82E-02 | At5g57010 | Uncharacterized protein At5g57010/MHM17_13 |
| AT2G30210 | -1.26 | 1.12E-15 | 1.46E-13 | At2g30210 | Laccase |
| AT1G02390 | -1.26 | 4.45E-10 | 2.43E-08 | GPAT2 | Probable glycerol-3-phosphate acyltransferase 2 |
| AT2G19810 | -1.27 | 3.47E-09 | 1.56E-07 | At2g19810 | Zinc finger CCCH domain-containing protein 20 |
| AT5G66390 | -1.27 | 5.56E-14 | 5.85E-12 | PER72 | Peroxidase 72 |
| AT5G42180 | -1.27 | 1.24E-06 | 3.08E-05 | At5g42180 | peroxidase |
| AT3G03640 | -1.27 | 3.31E-16 | 4.80E-14 | BGLU25 | Probable inactive beta-glucosidase 25 |
| AT1G77200 | -1.29 | 1.58E-09 | 7.78E-08 | ERF037 | Ethylene-responsive transcription factor ERF037 |
| AT4G40065 | -1.29 | 1.58E-03 | 1.30E-02 | | |
| AT1G68470 | -1.30 | 2.95E-12 | 2.34E-10 | GT17 | Probable xyloglucan galactosyltransferase GT17 |
| AT1G54040 | -1.30 | 4.68E-05 | 7.08E-04 | ESP | Epithiospecifier protein |
| AT3G15400 | -1.30 | 1.19E-05 | 2.21E-04 | ATA20 | ATA20 protein |
| AT3G15350 | -1.30 | 3.28E-17 | 5.37E-15 | At3g15350 | |
| AT3G48520 | -1.31 | 7.09E-04 | 6.85E-03 | CYP94B3 | Cytochrome P450 94B3 |
| AT3G63060 | -1.31 | 2.16E-05 | 3.65E-04 | At3g63060 | RING-type E3 ubiquitin transferase |
| AT1G29240 | -1.31 | 4.04E-10 | 2.23E-08 | At1g29240 | |
| AT5G49990 | -1.31 | 3.84E-23 | 1.19E-20 | NAT5 | Nucleobase-ascorbate transporter 5 |
| AT1G09505 | -1.31 | 2.28E-11 | 1.59E-09 | | |
| AT5G44990 | -1.32 | 7.92E-03 | 4.52E-02 | At5g44990 | Glutathione S-transferase family protein |
| AT1G61080 | -1.32 | 7.45E-04 | 7.13E-03 | At1g61080 | Hydroxyproline-rich glycoprotein family protein |
| AT1G17020 | -1.32 | 2.52E-06 | 5.68E-05 | SRG1 | Protein SRG1 |
| AT5G39580 | -1.32 | 1.28E-19 | 2.74E-17 | MIJ24.50 | peroxidase |
| AT2G41870 | -1.32 | 6.55E-15 | 7.82E-13 | REM4.2 | Remorin 4.2 |
| AT1G30935 | -1.32 | 8.16E-05 | 1.14E-03 | FBX12 | F-box only protein 12 |
| AT5G19890 | -1.32 | 7.66E-07 | 2.02E-05 | At5g19890 | Peroxidase |
| AT1G71490 | -1.32 | 4.61E-03 | 2.99E-02 | PCMP-E67 | Pentatricopeptide repeat-containing protein At1g71490 |
| AT1G16410 | -1.33 | 3.34E-16 | 4.81E-14 | CYP79F1 | Dihomomethionine N-hydroxylase |
| AT5G24240 | -1.33 | 2.64E-08 | 9.85E-07 | PI4KG3 | Phosphatidylinositol 4-kinase gamma 3 |
| AT1G47590 | -1.33 | 1.97E-03 | 1.55E-02 | PUP20 | Putative purine permease 20 |
| AT1G14490 | -1.33 | 3.40E-05 | 5.43E-04 | AHL28 | AT-hook motif nuclear-localized protein |
| AT3G53840 | -1.33 | 1.58E-03 | 1.30E-02 | WAKL15 | Wall-associated receptor kinase-like 15 |
| AT5G17460 | -1.33 | 1.26E-12 | 1.05E-10 | At5g17460 | Uncharacterized protein |
| AT1G45616 | -1.33 | 3.09E-03 | 2.20E-02 | RLP6 | Receptor-like protein 6 |
| AT1G53470 | -1.34 | 4.78E-11 | 3.16E-09 | MSL4 | Mechanosensitive ion channel protein 4 |
| AT4G16146 | -1.34 | 4.18E-03 | 2.78E-02 | At4g16146 | cAMP-regulated phosphoprotein 19-related protein |
| AT2G28815 | -1.35 | 6.02E-04 | 5.99E-03 | At2g28815 | Uncharacterized protein |
| AT4G33710 | -1.35 | 3.13E-06 | 6.86E-05 | At4g33710 | SCP domain-containing protein |
| AT5G15190 | -1.35 | 1.10E-09 | 5.64E-08 | F8M21_80 | Uncharacterized protein F8M21_80 |
| AT4G30830 | -1.35 | 1.45E-05 | 2.61E-04 | At4g30830 | GTD-binding domain-containing protein |
| AT3G61890 | -1.35 | 2.21E-09 | 1.05E-07 | ATHB-12 | Homeobox-leucine zipper protein ATHB-12 |
| AT1G44414 | -1.36 | 3.87E-03 | 2.62E-02 | At1g44414 | Zinc-ribbon 15 domain-containing protein |
| AT2G22590 | -1.36 | 4.82E-08 | 1.71E-06 | UGT91A1 | UDP-glycosyltransferase 91A1 |
| AT2G43570 | -1.36 | 4.93E-12 | 3.79E-10 | CHI | Endochitinase CHI |
| AT2G30670 | -1.36 | 1.00E-09 | 5.19E-08 | At2g30670 | Tropinone reductase homolog At2g30670 |
| AT3G48240 | -1.37 | 8.21E-05 | 1.14E-03 | At3g48240 | PB1 domain-containing protein |
| AT4G09820 | -1.37 | 8.20E-05 | 1.14E-03 | TT8 | Transcription factor TT8 |
| AT5G15970 | -1.37 | 8.94E-11 | 5.57E-09 | KIN2 | Stress-induced protein KIN2 |
| AT5G09570 | -1.38 | 4.75E-09 | 2.08E-07 | At5g09570 | |
| AT5G02550 | -1.38 | 7.67E-09 | 3.26E-07 | At5g02550 | |
| AT3G11410 | -1.39 | 5.62E-08 | 1.96E-06 | PP2CA | Protein phosphatase 2C 37 |
| AT2G27300 | -1.39 | 2.11E-03 | 1.64E-02 | NTL8 | NAC domain-containing protein 40 |
| AT5G24640 | -1.39 | 6.94E-08 | 2.36E-06 | At5g24640 | Uncharacterized protein |
| AT3G11370 | -1.40 | 4.79E-06 | 9.97E-05 | At3g11370 | DC1 domain-containing protein |
| AT3G46970 | -1.40 | 9.83E-13 | 8.31E-11 | PHS2 | Alpha-glucan phosphorylase 2, cytosolic |
| AT4G25990 | -1.40 | 5.79E-11 | 3.74E-09 | CIL | CCT motif family protein |

| | | | | | |
|---|---|---|---|---|---|
| AT5G46460 | -1.41 | 5.14E-04 | 5.25E-03 | PCMP-H49 | Pentatricopeptide repeat-containing protein At5g46460, mitochondrial |
| AT3G48510 | -1.41 | 2.66E-04 | 3.06E-03 | At3g48510 | |
| AT1G07367 | -1.41 | 2.24E-11 | 1.56E-09 | | |
| AT1G70800 | -1.42 | 1.24E-04 | 1.61E-03 | CAR6 | Protein C2-DOMAIN ABA-RELATED 6 |
| AT5G11110 | -1.43 | 1.77E-22 | 5.00E-20 | SPS2 | Probable sucrose-phosphate synthase 2 |
| AT1G29395 | -1.43 | 3.57E-12 | 2.81E-10 | At1g29390 | Uncharacterized protein |
| AT3G58150 | -1.43 | 1.18E-04 | 1.54E-03 | F9D24.60 | Uncharacterized protein F9D24.60 |
| AT4G15150 | -1.44 | 7.77E-08 | 2.60E-06 | dl3620c | Glycine-rich protein |
| AT2G47550 | -1.44 | 7.74E-06 | 1.51E-04 | PME20 | Probable pectinesterase/pectinesterase inhibitor 20 [Includes: Pectinesterase inhibitor 20 |
| AT1G17745 | -1.44 | 9.50E-13 | 8.07E-11 | PGDH2 | D-3-phosphoglycerate dehydrogenase 2, chloroplastic |
| AT2G29220 | -1.44 | 1.34E-04 | 1.72E-03 | LECRK31 | Probable inactive L-type lectin-domain containing receptor kinase III.1 |
| AT5G24770 | -1.45 | 4.48E-11 | 3.00E-09 | VSP2 | Vegetative storage protein 2 |
| AT5G12020 | -1.45 | 4.46E-16 | 6.15E-14 | HSP17.6 | 17.6 kDa class II heat shock protein |
| AT3G62590 | -1.45 | 3.75E-09 | 1.67E-07 | PLIP3 | Phospholipase A1 PLIP3, chloroplastic |
| AT3G46230 | -1.45 | 1.00E-04 | 1.35E-03 | HSP17.4A | 17.4 kDa class I heat shock protein |
| AT5G43300 | -1.46 | 8.09E-03 | 4.60E-02 | GDPD3 | glycerophosphodiester phosphodiesterase |
| AT3G46080 | -1.46 | 2.85E-03 | 2.06E-02 | ZAT8 | Zinc finger protein ZAT8 |
| AT5G42655 | -1.46 | 3.90E-05 | 6.10E-04 | At5g42655 | Dirigent protein |
| AT4G11910 | -1.46 | 6.74E-04 | 6.60E-03 | SGR2 | Magnesium dechelatase SGR2, chloroplastic |
| AT4G35300 | -1.46 | 3.23E-09 | 1.47E-07 | TMT2 | Tonoplast monosaccharide transporter2 |
| AT1G16030 | -1.47 | 2.02E-12 | 1.66E-10 | HSP70-5 | Heat shock 70 kDa protein 5 |
| AT2G39800 | -1.47 | 2.86E-09 | 1.30E-07 | P5CS1 | Delta-1-pyrroline-5-carboxylate synthase [Includes: Glutamate 5-kinase |
| AT5G57500 | -1.47 | 1.50E-03 | 1.25E-02 | At5g57500 | Hexosyltransferase |
| AT5G03350 | -1.47 | 9.94E-04 | 9.02E-03 | LLP | Lectin-like protein |
| AT4G32810 | -1.47 | 2.92E-07 | 8.53E-06 | CCD8 | Carotenoid cleavage dioxygenase 8 |
| AT4G13310 | -1.48 | 2.78E-12 | 2.21E-10 | CYP71A20 | Cytochrome P450, family 71, subfamily A, polypeptide 20 |
| AT1G54575 | -1.49 | 3.57E-05 | 5.64E-04 | At1g54575 | Uncharacterized protein |
| AT5G66460 | -1.49 | 3.89E-12 | 3.05E-10 | MAN7 | Mannan endo-1,4-beta-mannosidase 7 |
| AT1G48000 | -1.50 | 6.68E-06 | 1.33E-04 | T2J15.9 | Myb-related transcription factor |
| AT3G09910 | -1.50 | 3.24E-05 | 5.22E-04 | RABC2B | RAB GTPase homolog C2B |
| AT2G30660 | -1.50 | 4.44E-05 | 6.80E-04 | At2g30660 | 3-hydroxyisobutyryl-CoA hydrolase |
| AT1G52130 | -1.50 | 3.55E-03 | 2.45E-02 | JAL13 | Jacalin-related lectin 13 |
| AT4G12290 | -1.50 | 1.35E-24 | 4.78E-22 | CUAO | Amine oxidase |
| AT1G68240 | -1.50 | 7.10E-04 | 6.85E-03 | BHLH109 | Transcription factor bHLH109 |
| AT1G70300 | -1.51 | 2.53E-22 | 7.02E-20 | POT6 | Potassium transporter 6 |
| AT1G19250 | -1.51 | 6.80E-04 | 6.64E-03 | FMO1 | Probable flavin-containing monooxygenase 1 |
| AT4G13560 | -1.51 | 7.40E-03 | 4.28E-02 | UNE15 | Late embryogenesis abundant protein |
| AT5G11190 | -1.51 | 7.12E-04 | 6.85E-03 | SHN2 | Ethylene-responsive transcription factor SHINE 2 |
| AT2G30770 | -1.52 | 1.94E-05 | 3.32E-04 | CYP71A13 | Indoleacetaldoxime dehydratase |
| AT5G58630 | -1.52 | 1.22E-03 | 1.06E-02 | TRM31 | DUF3741 domain-containing protein |
| AT1G56300 | -1.52 | 8.93E-07 | 2.33E-05 | F14G9.9 | DnaJ protein, putative |
| AT1G64660 | -1.52 | 1.68E-08 | 6.56E-07 | MGL | Methionine gamma-lyase |
| AT4G15910 | -1.53 | 4.65E-15 | 5.68E-13 | LEA41 | Late embryogenis abundant protein 41 |
| AT1G34050 | -1.53 | 4.90E-10 | 2.66E-08 | At1g34050 | Ankyrin repeat family protein |
| AT5G09930 | -1.53 | 3.47E-13 | 3.18E-11 | ABCF2 | ABC transporter F family member 2 |
| AT5G60220 | -1.54 | 5.13E-04 | 5.24E-03 | At5g60220 | Tetraspanin-3 |
| AT3G09220 | -1.54 | 2.16E-04 | 2.57E-03 | LAC7 | Laccase |
| AT2G36270 | -1.55 | 3.53E-07 | 1.02E-05 | ABI5 | Basic-leucine zipper |
| AT2G40113 | -1.55 | 4.28E-07 | 1.20E-05 | At2g40113 | Pollen Ole e 1 allergen and extensin family protein |
| AT1G17870 | -1.55 | 3.56E-18 | 6.42E-16 | EGY3 | Probable zinc metallopeptidase EGY3, chloroplastic |
| AT2G26560 | -1.56 | 5.23E-07 | 1.44E-05 | PLP2 | Patatin-like protein 2 |
| AT1G30750 | -1.56 | 3.33E-11 | 2.27E-09 | T5I8.20 | T5I8.20 protein |
| AT2G29630 | -1.56 | 3.05E-08 | 1.12E-06 | THIC | ThiaminC |
| AT1G32180 | -1.56 | 2.64E-03 | 1.94E-02 | CSLD6 | Putative cellulose synthase-like protein D6 |
| AT3G52180 | -1.56 | 2.87E-19 | 5.81E-17 | At3g52180 | Uncharacterized protein At3g52180 |
| AT1G63530 | -1.57 | 1.20E-17 | 2.04E-15 | At1g63530 | F2K11.11 |
| AT3G08860 | -1.57 | 2.86E-08 | 1.06E-06 | PYD4 | PYRIMIDINE 4 |
| AT5G08490 | -1.57 | 3.75E-03 | 2.55E-02 | SLG1 | Tetratricopeptide repeat |
| AT2G29500 | -1.57 | 3.81E-13 | 3.44E-11 | HSP17.6B | 17.6 kDa class I heat shock protein 2 |
| AT4G06195 | -1.57 | 8.06E-05 | 1.13E-03 | | |
| AT3G14225 | -1.58 | 2.48E-04 | 2.90E-03 | GLIP4 | GDSL-motif lipase 4 |
| AT3G29970 | -1.59 | 2.78E-09 | 1.28E-07 | At3g29970 | B12D protein |
| AT4G03820 | -1.59 | 1.78E-19 | 3.73E-17 | At4g03820 | Transmembrane protein, putative |
| AT4G22880 | -1.59 | 1.63E-06 | 3.91E-05 | At4g22880 | Mutant protein of leucoanthocyanidin dioxygenase |
| AT1G05340 | -1.60 | 2.85E-09 | 1.30E-07 | CYSTM1 | Protein CYSTEINE-RICH TRANSMEMBRANE MODULE 1 |
| AT5G51465 | -1.60 | 3.37E-05 | 5.40E-04 | PROSCOOP18 | Serine rich endogenous peptide 18 |
| AT2G02010 | -1.61 | 1.58E-08 | 6.17E-07 | GAD4 | Glutamate decarboxylase |
| AT5G52940 | -1.61 | 2.43E-06 | 5.51E-05 | ATDOB5 | DUF295 domain-containing protein |
| AT5G01640 | -1.62 | 1.90E-20 | 4.30E-18 | PRA1B5 | PRA1 family protein B5 |
| AT1G28260 | -1.62 | 2.56E-19 | 5.25E-17 | SMG7L | Nonsense-mediated mRNA decay factor SMG7-like |
| AT1G22990 | -1.63 | 2.54E-07 | 7.59E-06 | HIPP22 | Heavy metal transport/detoxification superfamily protein |
| AT2G28500 | -1.63 | 3.09E-08 | 1.14E-06 | At2g28500 | LOB domain-containing protein |
| AT5G53660 | -1.64 | 5.55E-05 | 8.17E-04 | GRF7 | Growth-regulating factor 7 |
| AT5G01520 | -1.64 | 5.32E-11 | 3.48E-09 | AIRP2 | E3 ubiquitin-protein ligase AIRP2 |
| AT1G64160 | -1.66 | 6.57E-03 | 3.92E-02 | At1g64160 | Dirigent protein |
| AT2G39330 | -1.66 | 1.49E-05 | 2.67E-04 | JAL23 | Jacalin-related lectin 23 |
| AT1G53080 | -1.66 | 2.00E-05 | 3.43E-04 | At1g53080 | Lectin-like protein At1g53080 |
| AT1G11210 | -1.66 | 1.58E-06 | 3.81E-05 | At1g11210 | DUF4408 domain-containing protein |
| AT4G33150 | -1.66 | 9.34E-20 | 2.02E-17 | LKR | Lysine-ketoglutarate reductase/saccharopine dehydrogenase bifunctional enzyme |
| AT5G13320 | -1.66 | 2.69E-13 | 2.54E-11 | PBS3 | Auxin-responsive GH3 family protein |

| | | | | | |
|---|---|---|---|---|---|
| AT1G14520 | -1.68 | 3.33E-05 | 5.34E-04 | MIOX1 | Inositol oxygenase |
| AT1G09500 | -1.68 | 1.27E-04 | 1.64E-03 | At1g09500 | At1g09500/F14J9_16 |
| AT2G18050 | -1.68 | 7.25E-10 | 3.84E-08 | HIS1-3 | Histone H1-3 |
| AT4G19390 | -1.69 | 3.34E-23 | 1.05E-20 | At4g19390 | AT4g19390/T5K18_170 |
| AT1G13520 | -1.69 | 5.00E-09 | 2.19E-07 | F13B4.3 | F13B4.3 protein |
| AT2G41730 | -1.69 | 4.86E-19 | 9.44E-17 | At2g41730 | Calcium-binding site protein |
| AT1G05880 | -1.69 | 2.00E-08 | 7.66E-07 | ARI12 | RING/U-box superfamily protein |
| AT2G25820 | -1.69 | 4.24E-03 | 2.81E-02 | ERF042 | Ethylene-responsive transcription factor ERF042 |
| AT1G24600 | -1.70 | 1.65E-06 | 3.92E-05 | At1g24600 | |
| AT1G76960 | -1.70 | 8.46E-04 | 7.91E-03 | At1g76960 | F22K20.6 protein |
| AT1G55390 | -1.71 | 1.25E-03 | 1.08E-02 | At1g55390 | Cysteine/Histidine-rich C1 domain family protein |
| AT1G14550 | -1.71 | 3.73E-04 | 4.04E-03 | PER5 | Peroxidase 5 |
| AT1G73010 | -1.71 | 1.10E-15 | 1.45E-13 | At1g73010 | PS2 |
| AT5G62490 | -1.71 | 1.50E-06 | 3.64E-05 | HVA22B | HVA22-like protein b |
| AT1G52990 | -1.71 | 2.37E-08 | 8.88E-07 | F14G24.26 | Putative thioredoxin; 109829-109566 |
| AT4G21320 | -1.72 | 1.22E-10 | 7.28E-09 | HSA32 | Protein HEAT-STRESS-ASSOCIATED 32 |
| AT2G46680 | -1.73 | 1.97E-10 | 1.14E-08 | HB-7 | Homeobox-leucine zipper protein |
| AT1G21240 | -1.73 | 5.86E-06 | 1.19E-04 | WAK3 | Wall-associated receptor kinase-like |
| AT4G12410 | -1.74 | 3.12E-05 | 5.06E-04 | At4g12410 | SAUR-like auxin-responsive protein family |
| AT4G27410 | -1.74 | 2.09E-09 | 9.99E-08 | RD26 | NAC |
| AT1G16515 | -1.74 | 2.63E-13 | 2.49E-11 | At1g16515 | At1g16515 |
| AT2G16990 | -1.75 | 8.17E-10 | 4.29E-08 | At2g16990 | Major facilitator superfamily protein |
| AT3G03170 | -1.76 | 6.01E-11 | 3.87E-09 | O3L3 | Protein OXIDATIVE STRESS 3 LIKE 3 |
| AT1G03990 | -1.76 | 4.79E-12 | 3.71E-10 | At1g03990 | Long-chain-alcohol oxidase |
| AT5G54450 | -1.76 | 4.95E-05 | 7.42E-04 | At5g54450 | DUF295 domain-containing protein |
| AT4G07815 | -1.77 | 4.32E-05 | 6.65E-04 | | |
| AT4G22950 | -1.78 | 5.23E-04 | 5.33E-03 | AGL19 | AGAMOUS-like 19 |
| AT5G47530 | -1.78 | 2.13E-06 | 4.92E-05 | At5g47530 | Cytochrome b561 and DOMON domain-containing protein At5g47530 |
| AT4G25560 | -1.78 | 1.76E-05 | 3.06E-04 | LAF1 | Myb domain protein 18 |
| AT1G62290 | -1.78 | 1.36E-17 | 2.27E-15 | PASPA2 | Saposin-like aspartyl protease family protein |
| AT5G57510 | -1.79 | 1.44E-04 | 1.83E-03 | At5g57510 | |
| AT2G16367 | -1.79 | 1.42E-03 | 1.20E-02 | | |
| AT1G62620 | -1.80 | 5.14E-06 | 1.06E-04 | At1g62620 | Flavin-containing monooxygenase |
| AT4G13800 | -1.80 | 5.22E-10 | 2.81E-08 | ENOR3L3 | Probable magnesium transporter |
| AT5G51480 | -1.80 | 7.92E-05 | 1.11E-03 | SKS2 | Monocopper oxidase-like protein SKS2 |
| AT4G37420 | -1.81 | 6.88E-11 | 4.40E-09 | At4g37420 | Glycosyltransferase family 92 protein |
| AT4G39700 | -1.81 | 4.12E-04 | 4.36E-03 | HIPP23 | Heavy metal-associated isoprenylated plant protein 23 |
| AT1G03940 | -1.81 | 6.78E-32 | 4.12E-29 | At1g03940 | HXXXD-type acyl-transferase family protein |
| AT5G63350 | -1.81 | 8.55E-06 | 1.64E-04 | At5g63350 | |
| AT1G04570 | -1.82 | 1.19E-13 | 1.17E-11 | At1g04570 | Major facilitator superfamily protein |
| AT1G31750 | -1.83 | 4.00E-16 | 5.60E-14 | F27M3_5 | At1g31750 |
| AT2G37770 | -1.84 | 5.35E-05 | 7.91E-04 | AKR4C9 | NADPH-dependent aldo-keto reductase, chloroplastic |
| AT1G56600 | -1.85 | 7.56E-13 | 6.53E-11 | GOLS2 | Galactinol synthase 2 |
| AT5G22540 | -1.87 | 1.92E-04 | 2.33E-03 | MQJ16.8 | At5g22540 |
| AT4G31710 | -1.87 | 1.92E-07 | 5.85E-06 | GLR24 | Glutamate receptor |
| AT5G15500 | -1.88 | 1.50E-18 | 2.84E-16 | At5g15500 | Ankyrin repeat family protein |
| AT1G01250 | -1.88 | 1.99E-14 | 2.24E-12 | At1g01250 | |
| AT3G28510 | -1.89 | 1.43E-10 | 8.43E-09 | At3g28510 | AAA-ATPase At3g28510 |
| AT5G51760 | -1.89 | 1.75E-05 | 3.05E-04 | AHG1 | protein-serine/threonine phosphatase |
| AT1G60470 | -1.90 | 4.56E-11 | 3.03E-09 | GOLS4 | Galactinol synthase 4 |
| AT3G57010 | -1.90 | 1.07E-13 | 1.06E-11 | SSL8 | Protein STRICTOSIDINE SYNTHASE-LIKE 8 |
| AT2G29300 | -1.91 | 5.64E-08 | 1.96E-06 | At2g29300 | NAD |
| AT1G60970 | -1.91 | 5.22E-05 | 7.77E-04 | At1g60970 | Coatomer subunit zeta-1 |
| AT4G39210 | -1.92 | 8.66E-27 | 3.58E-24 | At4g39210 | Glucose-1-phosphate adenylyltransferase |
| AT5G01380 | -1.93 | 1.24E-05 | 2.28E-04 | GT-3A | Trihelix transcription factor GT-3a |
| AT2G24130 | -1.93 | 9.10E-05 | 1.24E-03 | At2g24130 | Leucine-rich receptor-like protein kinase family protein |
| AT3G48850 | -1.94 | 3.98E-04 | 4.24E-03 | MPT2 | Mitochondrial phosphate carrier protein 2, mitochondrial |
| AT1G47130 | -1.95 | 1.33E-04 | 1.70E-03 | At1g47130 | Uncharacterized protein |
| AT1G12064 | -1.96 | 7.73E-04 | 7.36E-03 | At1g12064 | Transmembrane protein |
| AT5G64750 | -1.96 | 2.47E-21 | 6.25E-19 | ABR1 | Ethylene-responsive transcription factor ABR1 |
| AT3G14360 | -1.96 | 1.16E-11 | 8.42E-10 | OBL1 | Triacylglycerol lipase OBL1 |
| AT2G35345 | -1.96 | 6.95E-04 | 6.75E-03 | At2g35345 | Uncharacterized protein |
| AT2G23830 | -1.97 | 3.79E-03 | 2.57E-02 | PVA31 | Vesicle-associated protein 3-1 |
| AT2G46950 | -1.99 | 3.01E-05 | 4.90E-04 | CYP709B2 | Cytochrome P450, family 709, subfamily B, polypeptide 2 |
| AT3G27250 | -1.99 | 1.35E-15 | 1.75E-13 | At3g27250 | |
| AT4G37710 | -2.00 | 2.50E-12 | 2.02E-10 | VQ29 | VQ motif-containing protein 29 |
| AT3G57520 | -2.01 | 2.96E-26 | 1.17E-23 | SIP2 | galactinol--sucrose galactosyltransferase |
| AT1G80110 | -2.01 | 3.01E-11 | 2.07E-09 | PP2B11 | F-box protein PP2-B11 |
| AT2G23110 | -2.02 | 3.73E-04 | 4.04E-03 | At2g23110 | Late embryogenesis abundant protein, LEA-18 |
| AT1G05100 | -2.03 | 1.37E-08 | 5.51E-07 | MAPKKK18 | Mitogen-activated protein kinase kinase kinase 18 |
| AT1G66390 | -2.04 | 2.17E-04 | 2.58E-03 | MYB90 | Transcription factor MYB90 |
| AT4G13300 | -2.04 | 8.43E-05 | 1.17E-03 | TPS13 | |
| AT2G38465 | -2.05 | 1.15E-05 | 2.14E-04 | At2g38465 | |
| AT3G50390 | -2.05 | 1.04E-04 | 1.40E-03 | F11C1_230 | Uncharacterized protein F11C1_230 |
| AT1G02230 | -2.05 | 5.73E-04 | 5.75E-03 | NAC004 | NAC domain-containing protein 4 |
| AT1G67100 | -2.05 | 1.72E-14 | 1.97E-12 | LBD40 | LOB domain-containing protein 40 |
| AT1G55775 | -2.06 | 1.66E-10 | 9.67E-09 | At1g55775 | Heavy metal-associated domain protein |
| AT3G62730 | -2.09 | 1.24E-05 | 2.28E-04 | At3g62730 | |
| AT1G69260 | -2.09 | 2.69E-28 | 1.21E-25 | AFP1 | Ninja-family protein AFP1 |
| AT5G41280 | -2.11 | 2.72E-09 | 1.26E-07 | CRRSP57 | Cysteine-rich repeat secretory protein 57 |

| | | | | | |
|---|---|---|---|---|---|
| AT3G29590 | -2.12 | 1.07E-21 | 2.73E-19 | 5MAT | Malonyl-CoA:anthocyanidin 5-O-glucoside-6"-O-malonyltransferase |
| AT1G26208 | -2.13 | 1.30E-04 | 1.67E-03 | | |
| AT1G73000 | -2.15 | 3.15E-05 | 5.10E-04 | PYL3 | Abscisic acid receptor PYL3 |
| AT1G53540 | -2.15 | 3.44E-27 | 1.45E-24 | HSP17.6C | 17.6 kDa class I heat shock protein 3 |
| AT1G79900 | -2.16 | 3.75E-08 | 1.36E-06 | BAC2 | Mitochondrial arginine transporter BAC2 |
| AT1G73040 | -2.17 | 6.14E-05 | 8.95E-04 | JAL19 | Jacalin-related lectin 19 |
| AT4G17470 | -2.19 | 3.03E-16 | 4.47E-14 | At4g17470 | Alpha/beta-Hydrolases superfamily protein |
| AT4G25200 | -2.19 | 6.50E-17 | 9.95E-15 | HSP23.6 | 23.6 kDa heat shock protein, mitochondrial |
| AT3G50970 | -2.19 | 3.46E-25 | 1.25E-22 | XERO2 | Dehydrin Xero 2 |
| AT1G49570 | -2.20 | 9.68E-21 | 2.30E-18 | PER10 | Peroxidase 10 |
| AT1G62710 | -2.21 | 2.21E-35 | 1.72E-32 | bVPE | Vacuolar-processing enzyme beta-isozyme |
| AT3G28210 | -2.21 | 2.70E-12 | 2.16E-10 | SAP12 | Zinc finger AN1 domain-containing stress-associated protein 12 |
| AT5G22460 | -2.23 | 5.08E-09 | 2.22E-07 | MWD9.26 | Alpha/beta-Hydrolases superfamily protein |
| AT5G12030 | -2.23 | 1.88E-30 | 9.61E-28 | HSP17.7 | 17.7 kDa class II heat shock protein |
| AT2G38340 | -2.27 | 2.66E-07 | 7.90E-06 | DREB2E | Dehydration-responsive element-binding protein 2E |
| AT4G10510 | -2.29 | 3.89E-22 | 1.07E-19 | At4g10510 | Subtilase family protein |
| AT2G29460 | -2.29 | 1.15E-09 | 5.86E-08 | GSTU4 | Glutathione S-transferase U4 |
| AT2G41190 | -2.30 | 3.07E-20 | 6.80E-18 | At2g41190 | Transmembrane amino acid transporter family protein |
| AT4G33905 | -2.30 | 2.24E-14 | 2.49E-12 | At4g33905 | |
| AT1G30250 | -2.30 | 2.28E-05 | 3.83E-04 | At1g30250 | Uncharacterized protein |
| AT4G37370 | -2.30 | 2.50E-42 | 3.03E-39 | CYP81D8 | Cytochrome P450, family 81, subfamily D, polypeptide 8 |
| AT5G17220 | -2.31 | 7.33E-18 | 1.30E-15 | GSTF12 | Glutathione S-transferase F12 |
| AT3G01570 | -2.32 | 9.66E-07 | 2.49E-05 | At3g01570 | Oleosin 5 |
| AT5G59320 | -2.33 | 3.87E-04 | 4.14E-03 | LTP3 | Non-specific lipid-transfer protein 3 |
| AT5G40790 | -2.34 | 4.86E-12 | 3.75E-10 | At5g40790 | |
| AT4G31940 | -2.36 | 5.93E-39 | 5.24E-36 | CYP82C4 | Xanthotoxin 5-hydroxylase CYP82C4 |
| AT1G09950 | -2.36 | 3.20E-11 | 2.20E-09 | RAS1 | Protein RESPONSE TO ABA AND SALT 1 |
| AT1G54160 | -2.36 | 1.39E-20 | 3.22E-18 | NFYA5 | Nuclear transcription factor Y subunit A-5 |
| AT1G52560 | -2.37 | 4.63E-12 | 3.61E-10 | HSP26.5 | 26.5 kDa heat shock protein, mitochondrial |
| AT3G03341 | -2.38 | 1.63E-06 | 3.91E-05 | At3g03341 | |
| AT2G34610 | -2.39 | 9.92E-08 | 3.26E-06 | At2g34610 | Cotton fiber protein |
| AT5G40000 | -2.41 | 3.82E-13 | 3.44E-11 | At5g40000 | AAA-ATPase At5g40000 |
| AT2G35980 | -2.41 | 7.23E-10 | 3.84E-08 | NHL10 | NDR1/HIN1-like protein 10 |
| AT3G13672 | -2.42 | 3.64E-14 | 3.91E-12 | SINA2 | TRAF-like superfamily protein |
| AT3G24750 | -2.43 | 4.02E-22 | 1.09E-19 | LAZY5 | At3g24750 |
| AT5G22355 | -2.43 | 5.62E-07 | 1.54E-05 | At5g22355 | Zinc finger PHD-type domain-containing protein |
| AT4G17030 | -2.45 | 3.23E-13 | 2.99E-11 | EXLB1 | Expansin-like B1 |
| AT5G40800 | -2.47 | 4.60E-23 | 1.40E-20 | At5g40800 | |
| AT2G21590 | -2.47 | 1.19E-27 | 5.16E-25 | At2g21590 | Probable glucose-1-phosphate adenylyltransferase large subunit, chloroplastic |
| AT1G72660 | -2.48 | 1.04E-05 | 1.96E-04 | DRG1-3 | P-loop containing nucleoside triphosphate hydrolases superfamily protein |
| AT3G03480 | -2.49 | 3.07E-03 | 2.18E-02 | CHAT | |
| AT1G03070 | -2.49 | 6.86E-41 | 7.41E-38 | LFG4 | Protein LIFEGUARD 4 |
| AT2G44780 | -2.49 | 8.43E-06 | 1.63E-04 | | |
| AT2G18540 | -2.52 | 3.49E-05 | 5.55E-04 | At2g18540 | Cupin type-1 domain-containing protein |
| AT5G07700 | -2.54 | 5.54E-08 | 1.93E-06 | MYB76 | Transcription factor MYB76 |
| AT2G40170 | -2.55 | 5.81E-07 | 1.58E-05 | EM6 | Em-like protein GEA6 |
| AT1G15520 | -2.55 | 3.52E-15 | 4.39E-13 | ABCG40 | Pleiotropic drug resistance 12 |
| AT5G40010 | -2.55 | 8.25E-11 | 5.18E-09 | At5g40010 | AAA+ ATPase domain-containing protein |
| AT4G14090 | -2.55 | 7.51E-21 | 1.80E-18 | UGT75C1 | UDP-glycosyltransferase 75C1 |
| AT5G54165 | -2.57 | 6.00E-18 | 1.07E-15 | At5g54165 | |
| AT2G36640 | -2.60 | 1.26E-03 | 1.09E-02 | ECP63 | Embryonic cell protein 63 |
| AT4G25380 | -2.60 | 1.09E-10 | 6.59E-09 | SAP10 | Zinc finger A20 and AN1 domain-containing stress-associated protein 10 |
| AT5G03210 | -2.61 | 1.87E-09 | 8.98E-08 | DIP2 | Arabidopsis thaliana genomic DNA, chromosome 5, P1 clone:MOK16 |
| AT2G34850 | -2.62 | 5.05E-16 | 6.91E-14 | At2g34850 | Putative UDP-arabinose 4-epimerase 2 |
| AT5G13170 | -2.64 | 1.08E-23 | 3.62E-21 | SWEET15 | Bidirectional sugar transporter SWEET15 |
| AT5G42800 | -2.64 | 1.86E-33 | 1.34E-30 | DFRA | Dihydroflavonol 4-reductase |
| AT4G12580 | -2.65 | 3.98E-13 | 3.56E-11 | At4g12580 | At4g12580 |
| AT3G55940 | -2.66 | 5.64E-06 | 1.15E-04 | PLC7 | Phosphoinositide phospholipase C 7 |
| AT3G12955 | -2.67 | 3.42E-16 | 4.89E-14 | SAUR74 | At3g12955 |
| AT3G55880 | -2.68 | 5.90E-57 | 1.91E-53 | SUE4 | Alpha/beta hydrolase related protein |
| AT1G72760 | -2.72 | 5.70E-14 | 5.93E-12 | At1g72760 | Protein kinase superfamily protein |
| AT3G22910 | -2.74 | 5.53E-15 | 6.73E-13 | ACA13 | Putative calcium-transporting ATPase 13, plasma membrane-type |
| AT5G54060 | -2.75 | 3.81E-19 | 7.55E-17 | A3G2XYLT | Anthocyanidin 3-O-glucoside 2'''-O-xylosyltransferase |
| AT4G33550 | -2.75 | 2.53E-21 | 6.31E-19 | At4g33550 | At4g33550 |
| AT5G57550 | -2.76 | 1.86E-11 | 1.32E-09 | XTH25 | Probable xyloglucan endotransglucosylase/hydrolase protein 25 |
| AT5G59720 | -2.76 | 9.88E-21 | 2.32E-18 | HSP18.1 | 18.1 kDa class I heat shock protein |
| AT2G45570 | -2.76 | 4.85E-08 | 1.72E-06 | CYP76C2 | Cytochrome P450 76C2 |
| AT1G56650 | -2.82 | 4.69E-44 | 6.51E-41 | MYB75 | Transcription factor MYB75 |
| AT2G35070 | -2.86 | 1.46E-07 | 4.56E-06 | At2g35065/At2g35 | Uncharacterized protein |
| AT1G14880 | -2.88 | 3.08E-07 | 8.96E-06 | PCR1 | Protein PLANT CADMIUM RESISTANCE 1 |
| AT5G45630 | -2.89 | 6.58E-04 | 6.46E-03 | At5g45630 | |
| AT1G32350 | -2.89 | 1.52E-08 | 6.00E-07 | AOX1D | Ubiquinol oxidase |
| AT4G10250 | -2.91 | 8.10E-15 | 9.61E-13 | HSP22.0 | 22.0 kDa heat shock protein |
| AT5G49620 | -2.95 | 1.67E-09 | 8.17E-08 | At5g49620 | Uncharacterized protein |
| AT3G09640 | -2.97 | 7.19E-23 | 2.09E-20 | APX2 | L-ascorbate peroxidase 2, cytosolic |
| AT4G02280 | -2.98 | 1.19E-14 | 1.40E-12 | SUS3 | Sucrose synthase 3 |
| AT3G50980 | -2.98 | 6.95E-16 | 9.39E-14 | XERO1 | Dehydrin Xero 1 |
| AT4G02360 | -2.99 | 1.02E-10 | 6.27E-09 | At4g02360 | T14P8.17 |
| AT3G28007 | -3.00 | 7.99E-14 | 8.14E-12 | At3g28007 | SWEET4 |
| AT3G53980 | -3.08 | 2.96E-25 | 1.08E-22 | At3g53980 | Bifunctional inhibitor/plant lipid transfer protein/seed storage helical domain-containing protein |

| | | | | | |
|---|---|---|---|---|---|
| AT1G30190 | -3.09 | 8.37E-23 | 2.39E-20 | At1g30190 | Cotton fiber protein |
| AT1G30220 | -3.15 | 9.72E-31 | 5.25E-28 | INT2 | Probable inositol transporter 2 |
| AT1G57590 | -3.20 | 3.27E-23 | 1.04E-20 | PAE2 | Pectin acetylesterase 2 |
| AT5G59220 | -3.24 | 3.02E-18 | 5.54E-16 | HAI1 | PP2C protein |
| AT4G01985 | -3.25 | 6.60E-28 | 2.92E-25 | At4g01985 | Transmembrane protein |
| AT5G52310 | -3.27 | 2.32E-31 | 1.29E-28 | At5g52310 | AT5G52310 protein |
| AT2G25625 | -3.28 | 5.58E-06 | 1.14E-04 | At2g25625 | Uncharacterized protein |
| AT3G56275 | -3.31 | 4.34E-15 | 5.35E-13 | | |
| AT5G40382 | -3.32 | 1.01E-07 | 3.32E-06 | At5g40382 | Putative cytochrome c oxidase subunit 5C-4 |
| AT4G27150 | -3.34 | 4.10E-06 | 8.67E-05 | AT2S2 | 2S seed storage protein 2 |
| AT5G53680 | -3.35 | 1.39E-11 | 1.01E-09 | At5g53680 | RRM domain-containing protein |
| AT1G16850 | -3.36 | 1.13E-26 | 4.57E-24 | F6I1.15 | At1g16880 |
| AT3G02480 | -3.42 | 5.91E-50 | 1.15E-46 | At3g02480 | Stress-induced protein KIN2-like |
| AT3G21660 | -3.45 | 5.10E-17 | 8.07E-15 | At3g21660 | UBX domain-containing protein |
| AT1G71000 | -3.50 | 6.90E-10 | 3.69E-08 | At1g71000 | Chaperone DnaJ-domain superfamily protein |
| AT1G07985 | -3.52 | 7.19E-23 | 2.09E-20 | At1g07985 | T6D22.8 |
| AT1G52040 | -3.58 | 1.32E-32 | 8.55E-30 | MBP1 | Myrosinase-binding protein 1 |
| AT5G43840 | -3.58 | 1.56E-07 | 4.83E-06 | HSFA6A | Heat stress transcription factor A-6a |
| AT5G05220 | -3.59 | 2.42E-20 | 5.42E-18 | At5g05220 | Uncharacterized protein |
| AT4G36600 | -3.61 | 3.10E-07 | 8.99E-06 | At4g36600 | At4g36600 |
| AT2G21820 | -3.62 | 3.49E-09 | 1.56E-07 | At2g21820 | |
| AT5G01300 | -3.69 | 8.82E-16 | 1.18E-13 | At5g01300 | PEBP |
| AT2G37870 | -3.72 | 7.11E-08 | 2.41E-06 | At2g37870 | Bifunctional inhibitor/lipid-transfer protein/seed storage 2S albumin superfamily protein |
| AT4G27670 | -3.75 | 8.19E-50 | 1.45E-46 | HSP21 | Heat shock protein 21, chloroplastic |
| AT4G09600 | -3.76 | 6.32E-06 | 1.27E-04 | GASA3 | Gibberellin-regulated protein 3 |
| AT5G66110 | -3.76 | 3.21E-41 | 3.67E-38 | HIPP27 | Heavy metal transport/detoxification superfamily protein |
| AT4G08570 | -3.77 | 3.11E-26 | 1.21E-23 | HIPP24 | Heavy metal-associated isoprenylated plant protein 24 |
| AT2G14610 | -3.83 | 2.24E-38 | 1.89E-35 | At2g14610 | Pathogenesis-related protein 1 |
| AT3G62990 | -3.83 | 6.55E-15 | 7.82E-13 | T20O10_90 | Uncharacterized protein T20O10_90 |
| AT2G35300 | -3.95 | 3.22E-11 | 2.21E-09 | LEA18 | Late embryogenesis abundant protein 18 |
| AT1G61800 | -3.96 | 4.63E-21 | 1.13E-18 | GPT2 | Glucose-6-phosphate/phosphate translocator 2 |
| AT5G04380 | -3.99 | 3.40E-14 | 3.68E-12 | At5g04380 | S-adenosyl-L-methionine-dependent methyltransferases superfamily protein |
| AT5G59310 | -4.04 | 5.91E-09 | 2.56E-07 | At5g59310 | Non-specific lipid-transfer protein |
| AT3G28290 | -4.07 | 9.35E-24 | 3.19E-21 | At3g28290 | UPF0496 protein At3g28290 |
| AT1G07430 | -4.10 | 3.54E-40 | 3.28E-37 | AIP1 | Protein phosphatase 2C 3 |
| AT1G03790 | -4.18 | 9.15E-14 | 9.17E-12 | TZF4 | Zinc finger CCCH domain-containing protein 2 |
| AT3G03620 | -4.19 | 4.20E-14 | 4.46E-12 | DTX24 | Protein DETOXIFICATION 24 |
| AT2G47770 | -4.35 | 1.02E-30 | 5.37E-28 | TSPO | Translocator protein homolog |
| AT3G28300 | -4.52 | 3.71E-16 | 5.23E-14 | At3g28300 | UPF0496 protein At3g28300 |
| AT4G24000 | -4.54 | 1.31E-09 | 6.56E-08 | CSLG2 | Cellulose synthase like G2 |
| AT5G62800 | -4.56 | 1.84E-13 | 1.78E-11 | At5g62800 | RING-type E3 ubiquitin transferase |
| AT5G15960 | -4.60 | 2.38E-30 | 1.19E-27 | KIN1 | Stress-induced protein KIN1 |
| AT1G11925 | -4.61 | 1.04E-15 | 1.38E-13 | F12F1.21 | F12F1.21 protein |
| AT5G50360 | -4.62 | 1.73E-09 | 8.38E-08 | AITR5 | Uncharacterized protein At5g50360 |
| AT1G64110 | -4.63 | 1.13E-13 | 1.11E-11 | DAA1 | At1g64110/F22C12_22 |
| AT5G06760 | -4.67 | 1.62E-14 | 1.86E-12 | LEA46 | Late embryogenesis abundant protein 46 |
| AT5G07330 | -4.92 | 3.81E-47 | 6.18E-44 | At5g07330 | |
| AT2G19900 | -4.98 | 7.10E-19 | 1.35E-16 | NADP-ME1 | Malic enzyme |
| AT5G52300 | -5.05 | 8.27E-41 | 8.46E-38 | LTI65 | CAP160 protein |
| AT2G17680 | -5.11 | 2.65E-14 | 2.91E-12 | At2g17680 | DUF241 domain protein, |
| AT1G52690 | -5.29 | 2.68E-07 | 7.93E-06 | LEA7 | Late embryogenesis abundant protein 7 |
| AT2G29380 | -5.31 | 4.55E-29 | 2.16E-26 | At2g29380 | Probable protein phosphatase 2C 24 |
| AT2G03130 | -5.41 | 9.07E-14 | 9.14E-12 | At2g03130 | Ribosomal protein L7/L12 C-terminal domain-containing protein |
| AT2G42000 | -5.62 | 1.47E-06 | 3.57E-05 | ATMT4A | Plant EC metallothionein family protein |
| AT3G12960 | -6.16 | 2.29E-28 | 1.06E-25 | SMP1 | SEED MATURATION PROTEIN 1 |
| AT5G66400 | -6.20 | 3.91E-110 | 3.81E-106 | RAB18 | Dehydrin family protein |
| AT3G13784 | -6.25 | 1.73E-45 | 2.58E-42 | At3g13784 | |
| AT3G15670 | -6.53 | 4.78E-50 | 1.03E-46 | LEA29 | Late embryogenesis abundant protein 29 |
| AT5G54740 | -6.59 | 2.20E-16 | 3.26E-14 | SESA5 | 2S seed storage protein 5 |
| AT1G68250 | -7.02 | 1.97E-24 | 6.86E-22 | At1g68250 | At1g68250 |
| AT5G44310 | -7.27 | 2.81E-35 | 2.10E-32 | At5g44310 | Late embryogenesis abundant protein |
| AT3G17520 | -7.29 | 7.16E-20 | 1.56E-17 | lea32 | LEA |
| AT5G66780 | -7.73 | 5.77E-52 | 1.60E-48 | MUD21.2 | Late embryogenesis abundant protein |
| AT2G42560 | -7.93 | 2.26E-57 | 8.80E-54 | LEA25 | Late embryogenesis abundant domain-containing protein |
| AT3G51810 | -8.03 | 2.17E-16 | 3.24E-14 | EM1 | Late embryogenic abundant protein |
| AT1G04560 | -8.76 | 2.29E-19 | 4.74E-17 | At1g04560 | |
| AT5G35660 | -9.35 | 3.50E-29 | 1.70E-26 | MJE4.15 | Glycine-rich protein family |
| AT5G22470 | -9.39 | 2.03E-33 | 1.41E-30 | PARP3 | Poly [ADP-ribose] polymerase |
| AT5G15250 | -9.79 | 5.84E-43 | 7.58E-40 | FTSH6 | FTSH protease 6 |
| AT1G27461 | -9.93 | 2.51E-169 | 4.89E-165 | At1g27461 | (thale cress) hypothetical protein |
| AT4G31830 | -10.68 | 1.31E-60 | 6.39E-57 | At4g31830 | (thale cress) hypothetical protein |

Genes differentially expressed in 14 day-old *A. thaliana* seedlings inoculated with *Peribacillus frigoritolerans* T7-IITJ versus those inoculated with no bacteria (sterile PBS) with FDR < 0.05 are shown. The genes with $log_2$(fold-change) > 1 were considered upregulated and those with $log_2$(fold-change) < -1 were considered downregulated. The Arabidopsis genome initiative (AGI) codes and Araport 11 short description of each gene are also shown. FC: fold-change

**Supplemental Table S6. Gene Ontology enrichment of genes induced in *Arabidopsis thaliana* seedlings inoculated with *Peribacillus frigoritolerans* T7-IITJ**

| Enrichment term | Input Number | Background Number | *P*-Value | Corrected *P*-Value |
|---|---|---|---|---|
| | | **GO Biological process** | | |
| **photosynthesis** | 11 | 88 | 5.51E-09 | 9.95E-07 |
| **photosynthesis, light harvesting in photosystem I** | 8 | 40 | 2.67E-08 | 3.86E-06 |
| **photosynthetic electron transport chain** | 6 | 23 | 3.98E-07 | 2.88E-05 |
| **photosystem I** | 4 | 8 | 4.75E-06 | 2.64E-04 |
| **response to low light intensity stimulus** | 4 | 16 | 4.36E-05 | 1.97E-03 |
| **photosystem II oxygen evolving complex** | 5 | 36 | 5.69E-05 | 2.42E-03 |
| **photosynthetic electron transport in photosystem I** | 4 | 25 | 1.99E-04 | 6.54E-03 |
| **photosynthesis, light harvesting in photosystem II** | 3 | 9 | 2.10E-04 | 6.60E-03 |
| **phylloquinone biosynthetic process** | 3 | 13 | 5.19E-04 | 1.39E-02 |
| **response to iron ion** | 4 | 33 | 5.20E-04 | 1.39E-02 |
| **response to light stimulus** | 10 | 306 | 1.53E-03 | 3.81E-02 |
| **response to auxin** | 11 | 388 | 2.67E-03 | 5.42E-02 |
| response to cold | 10 | 489 | 3.16E-02 | 2.38E-01 |
| response to salicylic acid | 6 | 144 | 4.32E-03 | 7.13E-02 |
| unidimensional cell growth | 5 | 188 | 4.68E-02 | 2.95E-01 |
| response to high light intensity | 4 | 73 | 7.77E-03 | 1.06E-01 |
| response to red light | 4 | 85 | 1.28E-02 | 1.51E-01 |
| response to cytokinin | 4 | 93 | 1.70E-02 | 1.67E-01 |
| actin filament depolymerization | 3 | 25 | 2.78E-03 | 5.42E-02 |
| iron ion homeostasis | 3 | 39 | 8.77E-03 | 1.15E-01 |
| fatty acid metabolic process | 3 | 49 | 1.57E-02 | 1.64E-01 |
| response to reactive oxygen species | 3 | 68 | 3.53E-02 | 2.55E-01 |
| root hair elongation | 3 | 77 | 4.75E-02 | 2.95E-01 |
| positive regulation of reactive oxygen species biosynthetic process | 2 | 6 | 2.74E-03 | 5.42E-02 |
| regulation of cytokinesis | 2 | 8 | 4.34E-03 | 7.13E-02 |
| photosynthesis, light harvesting | 2 | 10 | 6.28E-03 | 9.26E-02 |
| hydrogen peroxide transmembrane transport | 2 | 12 | 8.54E-03 | 1.14E-01 |
| water homeostasis | 2 | 14 | 1.11E-02 | 1.36E-01 |
| cellular manganese ion homeostasis | 2 | 19 | 1.88E-02 | 1.77E-01 |
| fructose 1,6-bisphosphate metabolic process | 2 | 20 | 2.06E-02 | 1.88E-01 |
| mitotic spindle organization | 2 | 21 | 2.24E-02 | 2.00E-01 |
| iron ion transport | 2 | 22 | 2.42E-02 | 2.10E-01 |
| manganese ion transmembrane transporter activity | 2 | 25 | 3.02E-02 | 2.35E-01 |
| cellular iron ion homeostasis | 2 | 25 | 3.02E-02 | 2.35E-01 |
| terpenoid biosynthetic process | 2 | 25 | 3.02E-02 | 2.35E-01 |
| defense response to insect | 2 | 26 | 3.23E-02 | 2.38E-01 |
| unsaturated fatty acid biosynthetic process | 2 | 28 | 3.67E-02 | 2.60E-01 |
| | | **GO Cellular Component** | | |
| **thylakoid** | 24 | 299 | 4.45E-14 | 3.22E-11 |
| **chloroplast thylakoid** | 22 | 311 | 5.36E-12 | 1.94E-09 |
| **chloroplast thylakoid membrane** | 25 | 459 | 3.97E-11 | 9.56E-09 |
| **chloroplast thylakoid lumen** | 9 | 68 | 8.77E-08 | 1.06E-05 |
| **plastoglobule** | 9 | 102 | 2.03E-06 | 1.34E-04 |
| **chloroplast envelope** | 24 | 768 | 2.27E-06 | 1.37E-04 |
| **chloroplast** | 97 | 6172 | 7.18E-06 | 3.46E-04 |
| **chloroplast stroma** | 22 | 856 | 1.04E-04 | 3.94E-03 |
| **thylakoid lumen** | 6 | 76 | 1.87E-04 | 6.44E-03 |
| **extrinsic component of membrane** | 5 | 50 | 2.35E-04 | 7.08E-03 |
| **PSII associated light-harvesting complex II** | 3 | 11 | 3.42E-04 | 9.90E-03 |

| | | | | |
|---|---|---|---|---|
| **NAD(P)H dehydrogenase complex (plastoquinone)** | 3 | 19 | 1.36E-03 | 3.52E-02 |
| extracellular region | 55 | 3824 | 6.41E-03 | 9.26E-02 |
| extracellular space | 4 | 129 | 4.63E-02 | 2.95E-01 |
| actin cytoskeleton | 3 | 53 | 1.91E-02 | 1.77E-01 |
| condensed nuclear chromosome, centromeric region | 2 | 6 | 2.74E-03 | 5.42E-02 |
| chromosome passenger complex | 2 | 6 | 2.74E-03 | 5.42E-02 |
| mitotic checkpoint complex | 2 | 8 | 4.34E-03 | 7.13E-02 |
| photosystem II | 2 | 9 | 5.27E-03 | 8.28E-02 |
| spindle midzone | 2 | 9 | 5.27E-03 | 8.28E-02 |
| photosystem I reaction center | 2 | 14 | 1.11E-02 | 1.36E-01 |
| structural constituent of cell wall | 2 | 17 | 1.55E-02 | 1.64E-01 |
| spindle microtubule | 2 | 23 | 2.62E-02 | 2.18E-01 |
| cell periphery | 2 | 30 | 4.13E-02 | 2.82E-01 |
| **GO Molecular Function** | | | | |
| **chlorophyll binding** | 7 | 38 | 3.31E-07 | 2.88E-05 |
| **electron transport pathway of photosynthesis activity** | 4 | 9 | 6.81E-06 | 3.46E-04 |
| **intracellular sequestering of iron ion** | 4 | 20 | 9.27E-05 | 3.72E-03 |
| oxidation-reduction process | 13 | 679 | 2.49E-02 | 2.10E-01 |
| iron ion binding | 9 | 409 | 2.73E-02 | 2.24E-01 |
| calcium ion binding | 8 | 356 | 3.26E-02 | 2.38E-01 |
| mannan synthase activity | 3 | 50 | 1.65E-02 | 1.67E-01 |
| xyloglucan:xyloglucosyl transferase activity | 3 | 57 | 2.29E-02 | 2.02E-01 |
| serine-type carboxypeptidase activity | 3 | 59 | 2.49E-02 | 2.10E-01 |
| cellulose synthase activity | 3 | 73 | 4.18E-02 | 2.83E-01 |
| xyloglucan metabolic process | 3 | 78 | 4.90E-02 | 2.95E-01 |
| peptidyl-prolyl cis-trans isomerase activity | 3 | 78 | 4.90E-02 | 2.95E-01 |
| histone serine kinase activity | 2 | 6 | 2.74E-03 | 5.42E-02 |
| ribulose-bisphosphate carboxylase activity | 2 | 6 | 2.74E-03 | 5.42E-02 |
| histone kinase activity (H3-S10 specific) | 2 | 6 | 2.74E-03 | 5.42E-02 |
| glucosinolate glucohydrolase activity | 2 | 7 | 3.49E-03 | 6.65E-02 |
| histone phosphorylation | 2 | 8 | 4.34E-03 | 7.13E-02 |
| carboxy-lyase activity | 2 | 8 | 4.34E-03 | 7.13E-02 |
| thioglucosidase activity | 2 | 10 | 6.28E-03 | 9.26E-02 |
| ferroxidase activity | 2 | 10 | 6.28E-03 | 9.26E-02 |
| nonphotochemical quenching | 2 | 15 | 1.25E-02 | 1.51E-01 |
| oxidoreductase activity | 2 | 16 | 1.40E-02 | 1.58E-01 |
| ferric iron binding | 2 | 17 | 1.55E-02 | 1.64E-01 |
| glucomannan 4-beta-mannosyltransferase activity | 2 | 17 | 1.55E-02 | 1.64E-01 |
| 2 iron, 2 sulfur cluster binding | 2 | 24 | 2.82E-02 | 2.29E-01 |
| ferrous iron binding | 2 | 25 | 3.02E-02 | 2.35E-01 |
| iron ion transmembrane transporter activity | 2 | 26 | 3.23E-02 | 2.38E-01 |
| terpene synthase activity | 2 | 28 | 3.67E-02 | 2.60E-01 |
| syncytium formation | 2 | 29 | 3.90E-02 | 2.71E-01 |
| negative regulation of translation | 2 | 29 | 3.90E-02 | 2.71E-01 |

A Gene Ontology (GO) enrichment analysis of 445 upregulated genes (see Supplementary Table 4) in *A. thaliana* seedlings inoculated with *Peribacillus frigoritolerans* T7-IITJ using the KOBAS 3.0 software is shown. The frequency of occurance of a particular pathway in the set of 445 genes (input number) versus the frequency in the global *A. thaliana* geneset (background number) determined the enrichment. Pathways with *P*-value < 0.05 are presented (Fisher's exact test) while the corrected *P*-values show Benjamini and Hochberg FDR correction (enrichment terms with FDR < 0.05 are highlighted in bold).

**Supplemental Table S7. Enrichment of biological pathways within genes repressed in *Arabidopsis thaliana* seedlings inoculated with *Peribacillus frigoritolerans* T7-IITJ**

| **Enrichment term** | **Database** | **Input Number** | **Background Number** | ***P*-value** | **Corrected *P*-value** |
|---|---|---|---|---|---|
| Metabolic pathways | KEGG PATHWAY | 53 | 3392 | 3.1E-02 | 3.0E-01 |
| Biosynthesis of secondary metabolites | KEGG PATHWAY | 33 | 1664 | 4.1E-03 | 9.5E-02 |
| Protein processing in endoplasmic reticulum | KEGG PATHWAY | 14 | 326 | 6.6E-05 | 4.8E-03 |
| Plant hormone signal transduction | KEGG PATHWAY | 11 | 448 | 2.2E-02 | 2.4E-01 |
| MAPK signaling pathway - plant | KEGG PATHWAY | 10 | 221 | 4.8E-04 | 2.0E-02 |
| Phenylpropanoid biosynthesis | KEGG PATHWAY | 9 | 241 | 3.2E-03 | 8.9E-02 |
| Starch and sucrose metabolism | KEGG PATHWAY | 8 | 227 | 7.4E-03 | 1.5E-01 |
| anthocyanidin modification (Arabidopsis) | BioCyc | 5 | 42 | 2.3E-04 | 1.2E-02 |
| Glycine, serine and threonine metabolism | KEGG PATHWAY | 5 | 109 | 1.2E-02 | 1.8E-01 |
| beta-Alanine metabolism | KEGG PATHWAY | 4 | 69 | 1.1E-02 | 1.8E-01 |
| Galactose metabolism | KEGG PATHWAY | 4 | 76 | 1.5E-02 | 2.0E-01 |
| Flavonoid biosynthesis | KEGG PATHWAY | 3 | 35 | 1.0E-02 | 1.7E-01 |
| Tropane, piperidine and pyridine alkaloid biosynthesis | KEGG PATHWAY | 3 | 51 | 2.6E-02 | 2.6E-01 |
| phospholipases | BioCyc | 3 | 57 | 3.5E-02 | 3.1E-01 |
| stachyose biosynthesis | BioCyc | 2 | 6 | 3.8E-03 | 9.2E-02 |
| FAS signaling pathway | PANTHER | 2 | 7 | 4.8E-03 | 1.1E-01 |
| traumatin and (Z)-3-hexen-1-yl acetate biosynthesis | BioCyc | 2 | 10 | 8.6E-03 | 1.7E-01 |
| soybean saponin I biosynthesis | BioCyc | 2 | 13 | 1.3E-02 | 1.8E-01 |
| flavonoid biosynthesis (in equisetum) | BioCyc | 2 | 17 | 2.1E-02 | 2.3E-01 |
| glucosinolate biosynthesis from trihomomethionine | BioCyc | 2 | 19 | 2.5E-02 | 2.6E-01 |
| glucosinolate biosynthesis from homomethionine | BioCyc | 2 | 19 | 2.5E-02 | 2.6E-01 |
| glucosinolate biosynthesis from tetrahomomethionine | BioCyc | 2 | 19 | 2.5E-02 | 2.6E-01 |
| glucosinolate biosynthesis from pentahomomethionine | BioCyc | 2 | 19 | 2.5E-02 | 2.6E-01 |
| glucosinolate biosynthesis from hexahomomethionine | BioCyc | 2 | 23 | 3.5E-02 | 3.1E-01 |
| flavonoid biosynthesis | BioCyc | 2 | 27 | 4.6E-02 | 3.5E-01 |
| Sesquiterpenoid and triterpenoid biosynthesis | KEGG PATHWAY | 2 | 26 | 4.3E-02 | 3.5E-01 |

An enrichment analysis of 503 downregulated genes (see Supplementary Table 5) in *A. thaliana* seedlings inoculated with *Peribacillus frigoritolerans* T7-IITJ using the KOBAS 3.0 software is shown. The KEGG PATHWAY and BioCyc databases were used to identify the biological pathways. The frequency of occurance of a particular pathway in the set of 445 genes (input number) versus the frequency in the global *A. thaliana* geneset (background number) determined the enrichment. Pathways with $P$-value < 0.05 are presented (Fisher's exact test), while the corrected $P$-values include Benjamini and Hochberg FDR correction.

**Supplemental Table S8. Gene Ontology enrichment of genes repressed in *Arabidopsis thaliana* seedlings inoculated with *Peribacillus frigoritolerans* T7-IITJ**

| Enrichment term | Input Number | Background Number | *P*-Value | Corrected *P*-Value |
|---|---|---|---|---|
| **GO Biological process** | | | | |
| response to abscisic acid | 44 | 583 | 4.54E-21 | 3.68E-18 |
| response to water deprivation | 37 | 455 | 7.17E-19 | 2.90E-16 |
| response to salt stress | 27 | 511 | 4.70E-10 | 1.27E-07 |
| heme binding | 20 | 492 | 4.21E-06 | 4.88E-04 |
| cellular response to hypoxia | 16 | 294 | 1.17E-06 | 2.37E-04 |
| response to oxidative stress | 16 | 417 | 7.32E-05 | 4.94E-03 |
| response to cold | 13 | 489 | 7.59E-03 | 1.54E-01 |
| response to osmotic stress | 12 | 212 | 1.74E-05 | 1.76E-03 |
| response to heat | 12 | 228 | 3.44E-05 | 3.09E-03 |
| abscisic acid-activated signaling pathway | 9 | 144 | 9.57E-05 | 5.96E-03 |
| defense response to fungus | 9 | 289 | 9.77E-03 | 1.70E-01 |
| response to wounding | 9 | 303 | 1.29E-02 | 1.83E-01 |
| response to reactive oxygen species | 8 | 68 | 3.46E-06 | 4.67E-04 |
| leaf senescence | 8 | 156 | 8.09E-04 | 2.97E-02 |
| cold acclimation | 7 | 84 | 1.09E-04 | 6.31E-03 |
| response to stress | 7 | 107 | 4.39E-04 | 1.87E-02 |
| protein self-association | 7 | 118 | 7.61E-04 | 2.94E-02 |
| protein serine/threonine phosphatase activity | 7 | 144 | 2.27E-03 | 7.34E-02 |
| protein folding | 7 | 191 | 9.77E-03 | 1.70E-01 |
| negative regulation of abscisic acid-activated signaling pathway | 6 | 71 | 3.15E-04 | 1.42E-02 |
| seed development | 6 | 95 | 1.33E-03 | 4.49E-02 |
| protein complex oligomerization | 5 | 14 | 2.34E-06 | 3.79E-04 |
| anthocyanin-containing compound biosynthetic process | 5 | 30 | 5.59E-05 | 4.52E-03 |
| response to desiccation | 5 | 41 | 2.12E-04 | 1.14E-02 |
| systemic acquired resistance | 5 | 77 | 2.98E-03 | 8.92E-02 |
| response to hydrogen peroxide | 5 | 79 | 3.30E-03 | 8.92E-02 |
| response to gibberellin | 5 | 110 | 1.22E-02 | 1.83E-01 |
| response to jasmonic acid | 5 | 153 | 4.08E-02 | 3.46E-01 |
| hyperosmotic salinity response | 4 | 92 | 2.77E-02 | 2.71E-01 |
| response to nematode | 4 | 108 | 4.48E-02 | 3.46E-01 |
| response to salt | 3 | 20 | 2.47E-03 | 7.69E-02 |
| response to mannitol | 3 | 23 | 3.53E-03 | 9.22E-02 |
| jasmonic acid metabolic process | 3 | 35 | 1.03E-02 | 1.70E-01 |
| protein folding chaperone | 3 | 37 | 1.19E-02 | 1.81E-01 |
| response to insect | 3 | 40 | 1.44E-02 | 1.92E-01 |
| pollen tube development | 3 | 43 | 1.73E-02 | 2.12E-01 |
| fruit development | 3 | 44 | 1.83E-02 | 2.22E-01 |
| seed maturation | 3 | 45 | 1.94E-02 | 2.31E-01 |
| cellular response to water deprivation | 3 | 48 | 2.27E-02 | 2.46E-01 |
| defense response to virus | 3 | 49 | 2.39E-02 | 2.55E-01 |
| plant-type primary cell wall biogenesis | 3 | 56 | 3.31E-02 | 3.05E-01 |
| positive regulation of abscisic acid-activated signaling pathway | 3 | 59 | 3.76E-02 | 3.27E-01 |

| | | | | |
|---|---|---|---|---|
| heat acclimation | 3 | 62 | 4.23E-02 | 3.46E-01 |
| response to virus | 3 | 65 | 4.73E-02 | 3.46E-01 |
| proline biosynthetic process | 2 | 11 | 1.01E-02 | 1.70E-01 |
| thiamine diphosphate biosynthetic process | 2 | 11 | 1.01E-02 | 1.70E-01 |
| anion transport | 2 | 13 | 1.33E-02 | 1.83E-01 |
| glycogen biosynthetic process | 2 | 14 | 1.51E-02 | 1.96E-01 |
| cellular response to sulfate starvation | 2 | 15 | 1.70E-02 | 2.12E-01 |
| hyperosmotic response | 2 | 17 | 2.10E-02 | 2.34E-01 |
| somatic embryogenesis | 2 | 20 | 2.78E-02 | 2.71E-01 |
| stomatal closure | 2 | 21 | 3.02E-02 | 2.88E-01 |
| cellular response to unfolded protein | 2 | 25 | 4.06E-02 | 3.46E-01 |
| defense response to insect | 2 | 26 | 4.34E-02 | 3.46E-01 |
| response to unfolded protein | 2 | 27 | 4.62E-02 | 3.46E-01 |
| starch metabolic process | 2 | 27 | 4.62E-02 | 3.46E-01 |
| **GO Cellular Component** | | | | |
| membrane | 21 | 944 | 6.21E-03 | 1.32E-01 |
| extracellular region | 60 | 3824 | 2.11E-02 | 2.34E-01 |
| **GO Molecular Function** | | | | |
| peroxidase activity | 8 | 131 | 2.70E-04 | 1.29E-02 |
| protein dephosphorylation | 8 | 157 | 8.42E-04 | 2.97E-02 |
| unfolded protein binding | 7 | 153 | 3.13E-03 | 8.92E-02 |
| protein poly-ADP-ribosylation | 2 | 6 | 3.76E-03 | 9.22E-02 |
| phospholipase activity | 3 | 24 | 3.94E-03 | 9.37E-02 |
| iron ion binding | 12 | 409 | 4.89E-03 | 1.07E-01 |
| glucose-1-phosphate adenylyltransferase activity | 2 | 10 | 8.58E-03 | 1.65E-01 |
| galactose metabolic process | 3 | 33 | 8.88E-03 | 1.67E-01 |
| mechanosensitive ion channel activity | 2 | 12 | 1.16E-02 | 1.81E-01 |
| inositol 3-alpha-galactosyltransferase activity | 2 | 13 | 1.33E-02 | 1.83E-01 |
| NAD+ ADP-ribosyltransferase activity | 2 | 13 | 1.33E-02 | 1.83E-01 |
| oxidoreductase activity, acting on paired donors, with incorporation or redı | 5 | 115 | 1.45E-02 | 1.92E-01 |
| sucrose synthase activity | 2 | 15 | 1.70E-02 | 2.12E-01 |
| oxidation-reduction process | 15 | 679 | 2.00E-02 | 2.34E-01 |
| phospholipase A1 activity | 2 | 17 | 2.10E-02 | 2.34E-01 |
| hydroquinone:oxygen oxidoreductase activity | 2 | 21 | 3.02E-02 | 2.88E-01 |
| cellulose synthase (UDP-forming) activity | 3 | 56 | 3.31E-02 | 3.05E-01 |
| monooxygenase activity | 4 | 98 | 3.36E-02 | 3.06E-01 |
| fatty acid binding | 2 | 23 | 3.52E-02 | 3.10E-01 |
| oxidoreductase activity, acting on the CH-OH group of donors, NAD or N/ | 3 | 65 | 4.73E-02 | 3.46E-01 |
| lipid metabolic process | 4 | 107 | 4.36E-02 | 3.46E-01 |

A Gene Ontology (GO) enrichment analysis of 503 downregulated genes (see Supplementary Table 5) in *A. thaliana* seedlings inoculated with *Peribacillus frigoritolerans* T7-IITJ using the KOBAS 3.0 software is shown. The frequency of occurance of a particular pathway in the set of 445 genes (input number) versus the frequency in the global *A. thaliana* geneset (background number) determined the enrichment. Pathways with *P*-value < 0.05 are presented (Fisher's exact test) while the corrected *P*-values show Benjamini and Hochberg FDR correction (enrichment terms with FDR < 0.05 are highlighted in bold).

**Supplemental Table S9. Antibiotic resistance genes of *Peribacullus frigoritolerans* T7-IITJ**

| Protein | Function | Start (bp) | Stop (bp) | Strand |
|---|---|---|---|---|
| **Antibiotic resistance genes** | | | | |
| Chloramphenicol acetyltransferase | Antibiotic inactivation | 57748 | 58395 | + |
| Fosfomycin thiol transferase | Antibiotic inactivation | 70919 | 71341 | - |
| Glycopeptide resistance gene VanW | Antibiotic target alteration | 92806 | 93291 | - |
| class A Bacillus cereus Bc beta-lactamase | Antibiotic inactivation | 134652 | 135584 | - |
| Glycopeptide resistance gene cluster VanY | Antibiotic target alteration | 649333 | 649803 | + |
| Small multidrug resistance (SMR) antibiotic efflux pump | Antibiotic efflux | 3048370 | 3048717 | - |
| Small multidrug resistance (SMR) antibiotic efflux pump | Antibiotic efflux | 3048743 | 3049072 | - |
| Glycopeptide resistance gene VanY | Antibiotic target alteration | 3609753 | 3610616 | - |
| Glycopeptide resistance gene VanR | Antibiotic target alteration | 3611824 | 3612519 | - |

Antibiotic resistance proteins encoded by the *Peribacillus frigoritolerans* T7-IITJ genome, are shown. The genes were predicted using the Comprehensive Antibiotic Resistance Database (CARD) identifier.

**Supplemental Table S10. Mobilome in *Peribacillus frigoritolerans* T7-IITJ**

| Contig | Mobile Genetic Elements | Details | Position (bp) | Length (bp) | GC Content (%) | Antibiotic Resistance Genes | Drug Class | Drugs | Virulence Gene | Virulence Description |
|---|---|---|---|---|---|---|---|---|---|---|
| 1 | Genomic_island | - | 2841421..2875463 | 34042 | 33.29 | - | - | - | - | - |
| 2 | Genomic_island | - | 153797..179435 | 25638 | 33.92 | - | - | - | - | - |
| 1 | Genomic_island | - | 88958..98937 | 9979 | 37.81 | - | - | - | - | - |
| 2 | Genomic_island | - | 759835..774066 | 14231 | 29.84 | - | - | - | - | - |
| 1 | Genomic_island | - | 133892..183945 | 50053 | 34.11 | - | - | - | - | - |
| 1 | Genomic_island | - | 1371003..1381358 | 10355 | 33.55 | - | - | - | - | - |
| 1 | Prophage | - | 1010681..1036228 | 25547 | 42.83 | - | - | - | - | - |
| 1 | Prophage | - | 3061151..3125829 | 64678 | 36.67 | - | - | - | - | - |
| 1 | Prophage | - | 2350832..2364872 | 14040 | 37.8 | - | - | - | - | - |
| 1 | Prophage | - | 3357828..3363034 | 5206 | 42.34 | - | - | - | - | - |
| 2 | Integrative_Conjugative_Elements | - | 1002146..1011212 | 9066 | 41.11 | - | - | - | - | - |
| 1 | IS/Tn | ISBce4 | 1313943..1314320 | 377 | 39.26 | - | - | - | - | - |
| 1 | IS/Tn | ISBwe2 | 1554160..1554831 | 671 | 39.2 | - | - | - | - | - |
| 1 | IScluster/Tn | ISBce14 / ISBce14 | 2851490..2852825 | 1335 | 34.76 | - | - | - | - | - |
| 2 | IScluster/Tn | ISPana1 / ISBwe1 | 1095632..1096516 | 884 | 38.12 | - | - | - | - | - |
| 2 | IScluster/Tn | ISBce14 / ISBce14 | 1223738..1224841 | 1103 | 34.63 | - | - | - | - | - |

Mobile genetic elements are shown along with their respective position (in base pairs) in the genome of *Peribacillus frigoritolerans* T7-IITJ. The mobile elements were detected by the VRprofile2 tool version 2.0.
